# Green cities and the risk for vector-borne disease transmission for humans and animals: a scoping review

**DOI:** 10.1101/2025.04.01.646559

**Authors:** Mathilde Mercat, Colombine Bartholomée, Florence Fournet, Magdalena Alcover Amengual, Maria Bourquia, Emilie Bouhsira, Anthony Cornel, Xavier Fernandez Cassi, Didier Fontenille, Adolfo Ibáñez-Justicia, Renaud Marti, Nicolas Moiroux, El Hadji Niang, Woutrina Smith, Jeroen Spitzen, Tessa M. Visser, Constantianus J.M. Koenraadt, Frederic Simard

## Abstract

**Background:** Greening cities is a nature-based strategy for sustainable urban development that integrates natural elements like plants or water bodies, to mitigate climate change impacts and enhance human well-being. However, urban green infrastructures (UGIs) can influence the distribution of disease vectors, potentially affecting vector-borne diseases (VBDs). UGIs may provide new suitable environments for urban vectors, while also creating opportunities to mitigate VBD risks through predation, competition, and dilution effects. This article examined the relationships between UGIs, vectors, and associated pathogens, impacting both human and animal health, highlighting knowledge gaps and identifying research priorities to support VBD risk mitigation measures and to guide smart urban planning and design.

**Methods:** A systematic literature search was conducted following PRISMA guidelines in three databases (Pubmed, Scopus, Web of Science). Selected articles involved (i) any aspect of a urban vector system, (ii) in UGIs, and (iii) statistical analysis of the effects of UGIs on VBD risk. Methods employed to characterize UGIs and VBDs were described and the identified impacts were summarized by vector group.

**Results:** Among the 98 articles reviewed, most addressed mosquito-pathogen systems (66), tick-pathogen systems (29), and few other vector-borne pathogen systems (3), with studies often confined to a single city or several cities within the same country and focused on one vector group. Urban vegetation generally appeared to heighten the risk of tick-borne diseases. In contrast, the influence of UGIs on the risk of mosquito-borne diseases varied depending on the vector system and on the environmental and climatic context. The diversity of indicators used to assess UGIs and VBD risks may affect the observed impact on VBD risk.

**Conclusion:** Given the increasing popularity of urban greening, it is crucial to investigate its potential implications for public health, and thereby urban planning decisions. However, the lack of standardized protocols complicates the accurate assessment of the effects of UGIs on the risk for VBD emergence and transmission and consequently, on potential mitigation measures.

## Introduction

Urbanization is an ongoing global phenomenon that brings both opportunities and challenges. By 2050, approximately 70% of the world population will live in urban areas (UN Habitat, 2022). Urbanization offers opportunities to reduce poverty and inequality by increasing access to education and healthcare (United Nations, 2020). However, the rapid growth of cities, often poorly controlled, has led to social and economic inequalities, as well as health issues, primarily affecting the most vulnerable populations (Kuddus et al., 2020). City development drives environmental disruptions by modifying land use. As urban areas are responsible for most of the world’s carbon emissions, increasing urbanization could further exacerbate pollution (UN Habitat, 2022). Urban landscapes are increasingly vulnerable to extreme events such as flooding or urban heat islands, negatively impacting human health (Heaviside et al., 2017). Climate change is expected to intensify both the frequency and severity of these events (Vautard et al., 2020). Additionally, urbanization induces habitat loss and fragmentation, leading to decreased biodiversity (E. L. Jones & Leather, 2012). Urban landscapes are a mosaic of highly heterogeneous environments (Breuste et al., 2008) that form complex interfaces connecting natural, rural, peri-urban, and urban ecosystems (Honnen & Monaghan, 2017). These interfaces increase interactions between humans, livestock, and wildlife, thereby increasing the risk of zoonotic pathogen emergence and transmission (Hassell et al., 2017; Pearce et al., 2018). High human density in urban areas and global connectivity through trade and travel further facilitate the introduction and spread of arthropod vectors and the pathogen agents they carry (Weaver & Reisen, 2010).

Greening cities is recognized as a key intervention to “make cities inclusive, safe, resilient, and sustainable,” as outlined by the United Nations (UN) Sustainable Development Goals (United Nations, 2022). Urban greening (Box 1) refers to the process of incorporating, developing, or preserving urban green infrastructures (UGIs) within urban environments through various initiatives. UGI (see Boxes 1 and 2) is an umbrella concept that includes a variety of elements such as blue spaces (e.g, lake, rivers and ponds (Hyseni et al., 2021)), green spaces (e.g, parks and green corridors, defined as interconnected landscape elements that help to protect the environment (Aly & Amer, 2010)), and urban forestry, including street and park trees. In this study, *greening* refers to the process itself, whereas *UGIs* denotes the tangible, physical components of urban nature that form the basis of greening efforts.

Urban green spaces mitigate environmental challenges and contribute to various ecosystem services. It reduces heat island effects (Gunawardena et al., 2017; Marando et al., 2022), alleviates flooding (Bai et al., 2018), and reduces air pollution by capturing carbon dioxide (Baró et al., 2014; Mexia et al., 2018). Moreover, urban green spaces restore and maintain city biodiversity (Archibald et al., 2017; Vergnes et al., 2012). Substantial evidence highlights the positive impact of urban greening on human well-being and health, including stress reduction, increased opportunities for physical activities that can lower the risk of non-communicable diseases like hypertension and obesity, and a decrease in respiratory diseases (Van Den Bosch & Ode Sang, 2017). These benefits align with recommendations from the recently established One Health High Level Expert Panel (OHHLEP), which promotes a holistic approach to health for people, animals, and the environment (WHO, 2021).

While UGIs provide ecological and health benefits, they can also create new habitats within cities that favor the establishment and spread of arthropod vectors of human and animal pathogens. This raises the risk of Vector-Borne Disease (VBD, see Box 1) emergence and transmission in urban areas (Fournet et al., 2024). Hematophagous (blood-feeding) arthropods are key players in transmitting pathogens that can impact human and animal health. Many VBDs are on the rise worldwide, with their arthropod vectors spreading beyond native ranges due to climate change, globalization, and the intensification of international trade and travel (Cuthbert et al., 2023). Arthropod vectors such as mosquitoes and ticks are particularly able to adapt to new environments, often acting as invasive species due to their specific ecological, biological, and genetic traits (Renault et al., 2024). As such, UGIs may inadvertently increase habitat suitability for these disease vectors in urban areas, thereby increasing human-vector contact in these green environments frequently used by people (Hansford et al., 2017). On the other hand, UGIs may also support the establishment of predators and competitors of urban arthropod vectors, facilitating novel trophic networks that could reduce vector populations and mitigate VBD risk (Prihandi & Nurvianto, 2022). Actually, increased biodiversity in UGIs can lead to shifts in vector population dynamics, host availability, and/or the presence of biological control agents, potentially resulting in dilution effects for pathogens (Keesing & Ostfeld, 2021).

This scoping review investigated how UGIs influence VBD risks for humans and animals. Its objective was to assess the potential future consequences of urban greening on VBD risks by specifically examining the impact of existing UGIs. The review aimed to: (i) assess the influence of different UGIs on the presence and abundance of vectors (e.g., mosquitoes, ticks, sand flies, and chigger mites) and VBD dynamics (e.g., dengue fever, West Nile fever, malaria, Lyme disease, leishmaniasis, and scrub typhus); (ii) identify UGIs characteristics such as connectivity, surface area, proximity to residential areas, and plants species that most affect vector populations and VBD dynamics; and (iii) compare the effects of UGIs on the same vector species across different locations, as well as within the same location when possible. Moreover, we identified critical knowledge gaps that require further research to inform VBD mitigation measures and guide the smart design of UGIs aiming to minimize VBD risks while fostering greener cities. This work will help city planners and policymakers in understanding the VBD risks linked to urban greening, tailored to each city’s epidemiological context, enabling them to implement landscape management strategies in a thoughtful and informed way.

### Box 1. General definitions

**Vector:** an invertebrate, typically an arthropod, that transmits an infectious agent (Duvallet et al., 2018; Wilson et al., 2017) from one host (human or animal) to another (Blanc & Gutiérrez, 2015).

**Vector Borne Disease (VBD):** human or animal diseases caused by parasites, viruses, and bacteria that are transmitted by vectors (WHO, 2020).

**Risk:** According to the United Nations Office for Disaster Risk Reduction (UNDRR) (UNDRR, 2019), a risk is a “potential loss of life, injury or destroyed assets which could occur to a system, society or a community, determined probabilistically as a function of hazard, exposure, vulnerability, and capacity.” The risk may be health-related, economic, ecological, social, or political. It can be summarized as risk = hazard x exposure x vulnerability.

**VBD risk:** According to the UNDRR disaster risk definition (UNDRR, 2019), VBD risk can be defined as a hazard linked to an infected vector x exposure to this vector x vulnerability (of the vertebrate host population).

**Urban greening:** Urban greening, also referred to as planning of green and blue urban infrastructure, is defined by the European Environment Agency (European Environment Agency (EEA), 2024) as *“a strategic approach to developing interconnected, multifunctional networks of blue and green spaces capable of delivering a wide range of environmental, social, and economic benefits, while simultaneously enhancing cities’ resilience to climate change.”* More broadly, it encompasses the incorporation, development, and preservation of green infrastructure in cities through diverse initiatives that strengthen urban ecosystems and promote human well-being. Although often framed as an environmental and public health strategy, urban greening also carries political dimensions, particularly in relation to urban planning and governance (Cooke, 2020).

**Urban green infrastructure (UGI):** Refers to the physical components of urban nature, encompassing both green spaces and blue spaces. Jones et al. (2022) introduced the term “Urban Green Infrastructure” to categorize various green and blue spaces within urban environments. Their typology resulted in nine categories, including gardens, parks, amenity areas, other public spaces (e.g., cemeteries), linear features, constructed green infrastructures (e.g., green roofs or green walls), and three additional categories for urban water spaces. In this review, the term *urban green infrastructure* is applied more broadly to also include vegetation in neglected urban spaces, such as wastelands and vacant lots (Yang et al., 2019), as well as urban agriculture farms and urban forests (see Box 2).

**Highly urbanized areas (or highly built urbanized areas):** In the articles included in this scoping review, UGIs were often compared with highly urbanized areas. According to the studies, these areas are characterized by impervious surfaces (Pedrosa et al., 2020; Schwarz et al., 2020) with dense buildings (Cox et al., 2007; Pedrosa et al., 2020; Schwarz et al., 2020), a high degree of urbanization (Vieira et al., 2020), and low vegetation cover (Abreu et al., 2022).

### Box 2. Classification of UGIs

We developed a classification system to describe urban green infrastructures (UGIs) as follows:

- **Urban green space typology (UGST):** This typology based on the framework of Jones et al. (2022), classifies urban green spaces according to their use, land cover, ownership, geometry, and management. For example, if an UGI is described as an’urban park’ or as the connectivity between urban green spaces, it is categorized under the class‘Urban green space typology’.

This classification allows detailed descriptions including:

○ **Use:** urban parks, cemeteries, zoos, urban farms, urban agriculture, urban forest, urban green spaces without any specified use, and vacant lot..
○ **Ownership:** residential areas, and publicly-owned parks.
○ **Geometry:** green roofs, and green walls.

Key characteristics of urban green spaces can be described by:

○ **Shape and size:** This refers to the structural characteristics of urban green spaces, including overall shape, perimeter, and surface area.
○ **Connectivity:** This term denotes the interconnectedness of green spaces or their connection to peri-urban environments.
○ **Relevant features:** This refers to outstanding areas within urban green spaces, such as trails and edges.
○ **Surrounding land cover:** This refers to the type of vegetation or soil present near urban green spaces. It could be the level of the canopy around urban green spaces or the percentage of shrubs/grass/and bare soil surrounding parks.
○ **Management:** This refers to maintenance approaches to the vegetation within the urban green space, such as grass-cutting or watering schedules.
○ **Distance to urban green spaces:** Distance of sampling points to urban green areas.
○ The distance within a urban green space (urban parks or urban forests) from its edge.
○ **Presence of specific animals,** like the presence of farmyard animals.

- **Urban vegetation description (UVD):** This description focuses on the detailed characterization of urban vegetation including land cover, vegetation indexes, and specific vegetation traits such as height, tree density, diversity, etc. It provides a comprehensive description of the vegetation around the sampling points across different studies. For example, vegetation described using Normalized Difference Vegetative Index (NDVI) within a 200-meter buffer around a mosquito trap will be included in this category. Key components include:

○ **Type of vegetation:** This describes vegetation based on its land cover type, including grass, long grass, deciduous, mixed or coniferous forests, woodland edge, bushlands and riparian forests.
○ **Litter composition:** This describes the composition of organic litter found within the vegetation, such as leaves and twigs.
○ **Vegetation index:** This index allows for an alternative vegetation description through remote sensing indices, the percentage of urban vegetation, tree cover percentage, canopy percentage cover, arboreal percentage cover or overall vegetation cover. The NDVI is a remote sensing index used to determine the presence of vegetation. It ranges from-1 to 1, and the higher it is, the denser the vegetation (Martinez & Labib, 2023).
○ **Vegetation characteristics:** This encompasses many aspects of vegetation traits, including height, diversity, shade, size of tree holes, tree density, vegetation diversity, and the presence of flowers.
○ **Plant species:** This involves a thorough characterization of vegetation, with a focus on the presence and attributes of specific plant species studied within the area.
○ **Distance to urban vegetation:** Distance of sampling points to urban vegetation.

## Methods

The ‘Preferred Reporting Items for Systematic Reviews and Meta-Analyses’ (PRISMA) guidelines (Page et al., 2021) were used to conduct a scoping review examining the links between UGIs, vectors, and VBD risk (Box 1). Key methodological steps are outlined in Figure 1 and detailed below. The PRISMA check-list is available in Additional file 1: Text S1.

**Figure 1.**
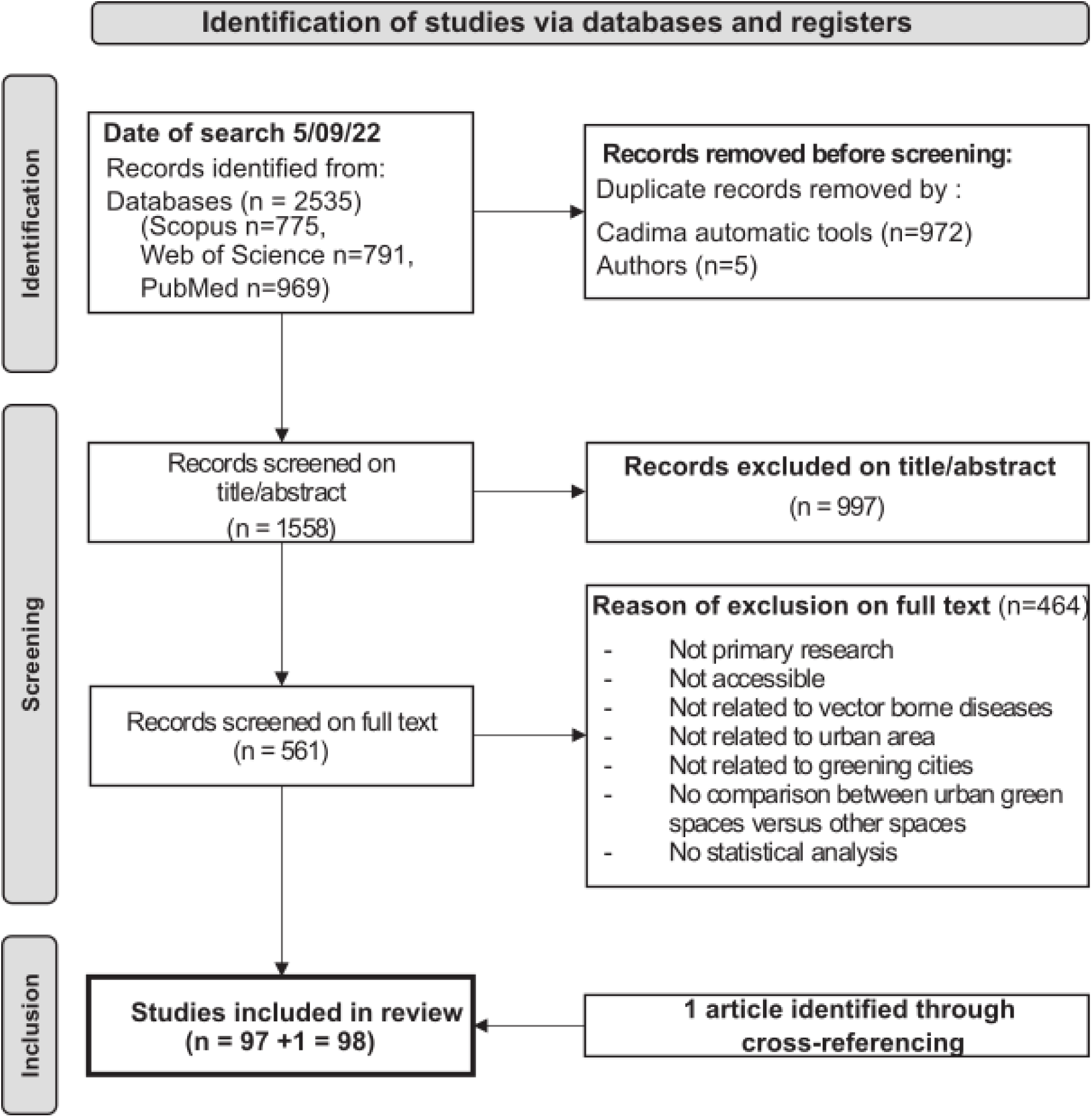
PRISMA flow diagram for systematic review. It shows the number of article identified through cross-checking references.

**Figure 2.**
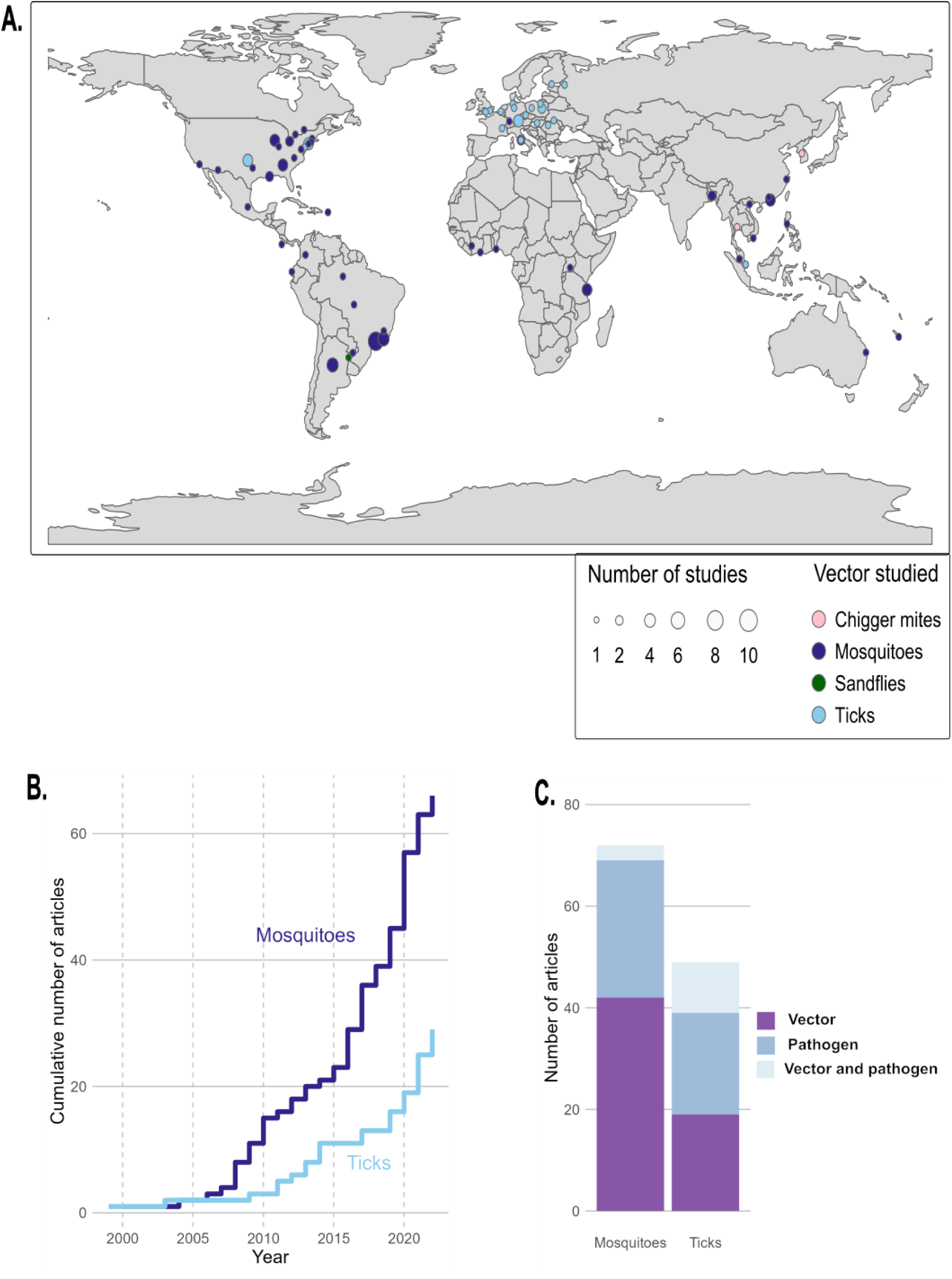
Overview of included studies. A study refers to the investigation of a specific vector type in a particular city and time by a research team; a single article may therefore include multiple studies. (A) Global distribution of studies (aggregated when they concern the same vector type in the same city). An interactive map with study references is available at at https://doi.org/10.5281/zenodo.17136279 (Bartholomée & Mercat, 2025b). (B) Cumulative number of studies by vector type and year of publication for mosquito-pathogen and tick-pathogen systems (aggregated when they share the same publication year and vector type). (C) Number of studies by data type for mosquito-pathogen and tick-pathogen systems.

### Search strategy

A comprehensive search was conducted in October 2022 using three online databases: PubMed, Scopus, and Web of Science. The search strategy combined key concepts related to “urban,” “greening,” “vector,” and “vector-borne disease.” Since the objective of this study was to assess the potential future consequences of urban greening on VBD by examining the effects of existing UGIs, the concept of *greening* was operationalized using related terms such as *forest, park*, and *vegetation*. All possible associated keywords for each key concept were included. A proximity operator was used when available to ensure that “greening” keywords referred specifically to urban areas (Additional file 2: Text S2). The query targeted titles, abstracts, and keywords, and searches were conducted using the CADIMA web tool (Kohl et al., 2018). All articles published up to October 6, 2022 (with no specified start date) were considered. Given that urban greening is a dynamic and relatively new management approach, relevant articles were expected to be limited. As a proxy for “greening cities”, all articles assessing any component of VBD risk related to any aspect of UGIs were included.

### Article selection

After the systematic database search, duplicates were automatically removed using CADIMA tool. The remaining articles underwent an initial systematic screening based on titles and abstracts, using the following inclusion criteria: articles (i) must be original research published in peer-reviewed journals in English; (ii) must address any aspect of the vector system in relation to human or animal health (e.g., vector species, pathogens, or hosts); and (iii) must focus on at least one UGI (e.g., urban park, green/blue corridor, vegetation index or similar). Subsequently, a full-text screening was conducted to further refine the selection. In this step, articles were included if they:

(i) referred specifically to urban areas; (ii) examined any aspect of UGIs (e.g., urban parks, vegetation cover indices, or tree species); and (iii) statistically analyzed the effect of different UGIs (e.g., multiple parks, varying vegetation cover, or different tree species) on VBD risk including hazard, exposure, and/or vulnerability. Commentaries, editorial reviews, opinion papers, and articles from grey literature were excluded, as well as those focusing on VBDs in urban environments without reference to green spaces. Two of the authors (MM and CB) independently reviewed both screenings (titles and abstracts, and full-text articles) for inclusion and extracted articles data, with any disagreement resolved by consensus. At the end of the selection process, the authors added one additional article. This study, cited in the *Discussion* section of one of the 98 selected articles, had not been retrieved in the systematic search because the terms “vector” or ‘vector-borne diseases’ did not appear in its title or abstract. Nevertheless, it met all other inclusion criteria and was considered relevant to the study objectives, thereby justifying its inclusion.

### Data extraction and description

From each article, the following metadata was extracted: location (continent, country, city), year of publication, year of sampling, vectors involved (i.e., groups of vectors (mosquitoes, ticks, sand flies, chigger mites), including genus, species, and life stage), as well as the disease and host of interest, when available. The methodology used for vector sampling was also documented, including the type of trap and frequency of collection, as well as the diagnostic tests employed to detect pathogens. The risk indicators from the studies were categorized into three groups: “vector data”, for indicators addressing vector presence, abundance, or density; **“**vector-borne pathogen data**”**, for indicators related to vector infection by pathogens or host disease prevalence/incidence; and **“**vector and vector-borne pathogen data**”**, for studies encompassing both vector and pathogen aspects. Metadata related to the description of UGIs were extracted as well, including the type of methodology used (e.g., general classification based on usage, remote sensing, or field description), the variables monitored (e.g., land-use classification or Normalized Difference Vegetation Index (NDVI)), and the buffer size used around sampling points, when available. The variables describing UGIs were classified into two categories: (1) urban green space typology (UGST) and (2) urban vegetation description (UVD) (Box 2).

Finally, the statistical framework used to analyze the effect of UGIs on VBD risk was also extracted from each article. All combinations of VBD risk and UGI variables (*VBD risk variable ∼ UGI variable*) tested statistically were collected. These combinations were analyzed independently, regardless of how many variables originated from the same article. In cases where a large proportion of variables were reported in a single article, each tested association was still treated as an independent interaction. For each interaction, the effect of UGI on VBD risk was classified into four categories: (i) harmful, when a variable describing the UGI increases VBD risk, (ii) beneficial, when a variable describing the UGI decreases VBD risk, (iii) variable when the effect of the UGI variable on VBD risk varies across sampling sites, independently of site characteristics, or (iv) no association when there was no significant relationship between the UGI variable and VBD risk. When the same variable was tested in different articles, all results were reported. For example, if ‘urban park’ had ‘beneficial’ effect on the VBD risk in one article but showed ‘no association’ in another, both outcomes were presented separately. The “variable” category refers to cases where the same UGI variable had opposite effects depending on the sampling point within a single study.

The data extracted from all of the articles can be found in Additional file 3: Dataset S1. All data and additional files are available online (Bartholomée & Mercat, 2025). Figures 3 to 5 summarize the combinations of UGIs and VBDs risk, according to the vector-pathogen system studied and the UGI or VBD indicators used. The number of variables included, and the number of articles from which they were extracted, are indicated. A variable reported in multiple articles is counted multiple times. The proportions shown on the x-axis represent the number of variables, within each UGI category (UGST or UVD) and vector-pathogen system, that were classified as harmful, beneficial, variable or showing no association.

**Figure 3.**
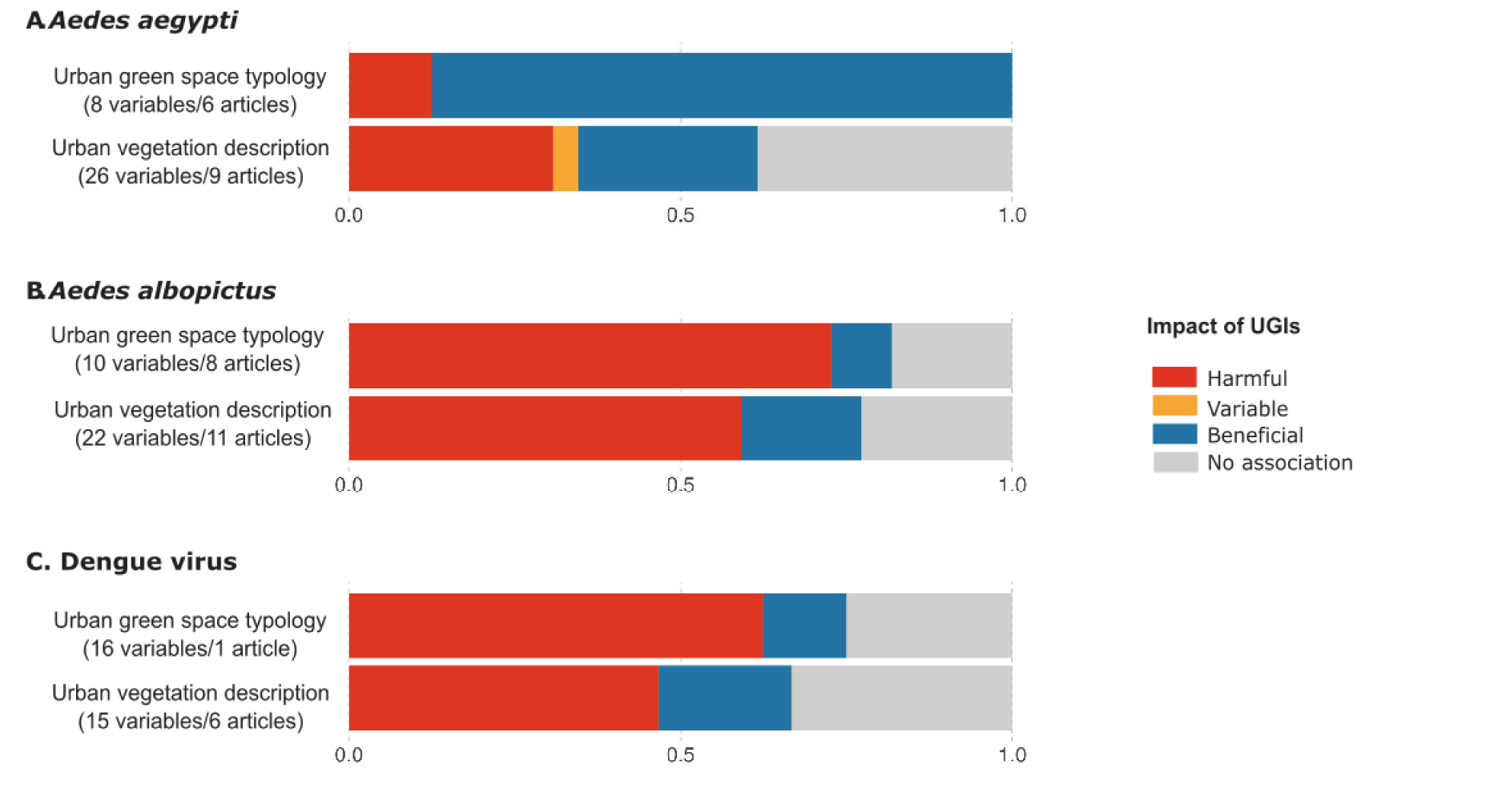
Impact of UGIs on *Aedes*-pathogen systems. (A) Impact of UGIs on *Aedes aegypti* populations. (B) Impact of UGIs on *Aedes albopictus* populations. (C) Impact of UGIs on Dengue virus dynamics (prevalence of infection in host or vector). Numbers in brackets indicate the total number of variables tested and the total number of articles from which these variables were extracted. Since some articles provide data across both categories, they may be counted more than once, which explains discrepancies with the total number of articles reported in Table 1. The same variable tested in multiple articles is also counted multiple times. Colors represent the proportion of variables in all studies that increased risk (harmful effect, red), decreased risk (beneficial effect, blue), varied (variable effect, yellow), or showed no association (grey) with risk. The x-axis represents the proportion of UGI indicators that, for a given vector-pathogen system, were classified as harmful, beneficial, variable, or showing no association.

**Figure 4.**
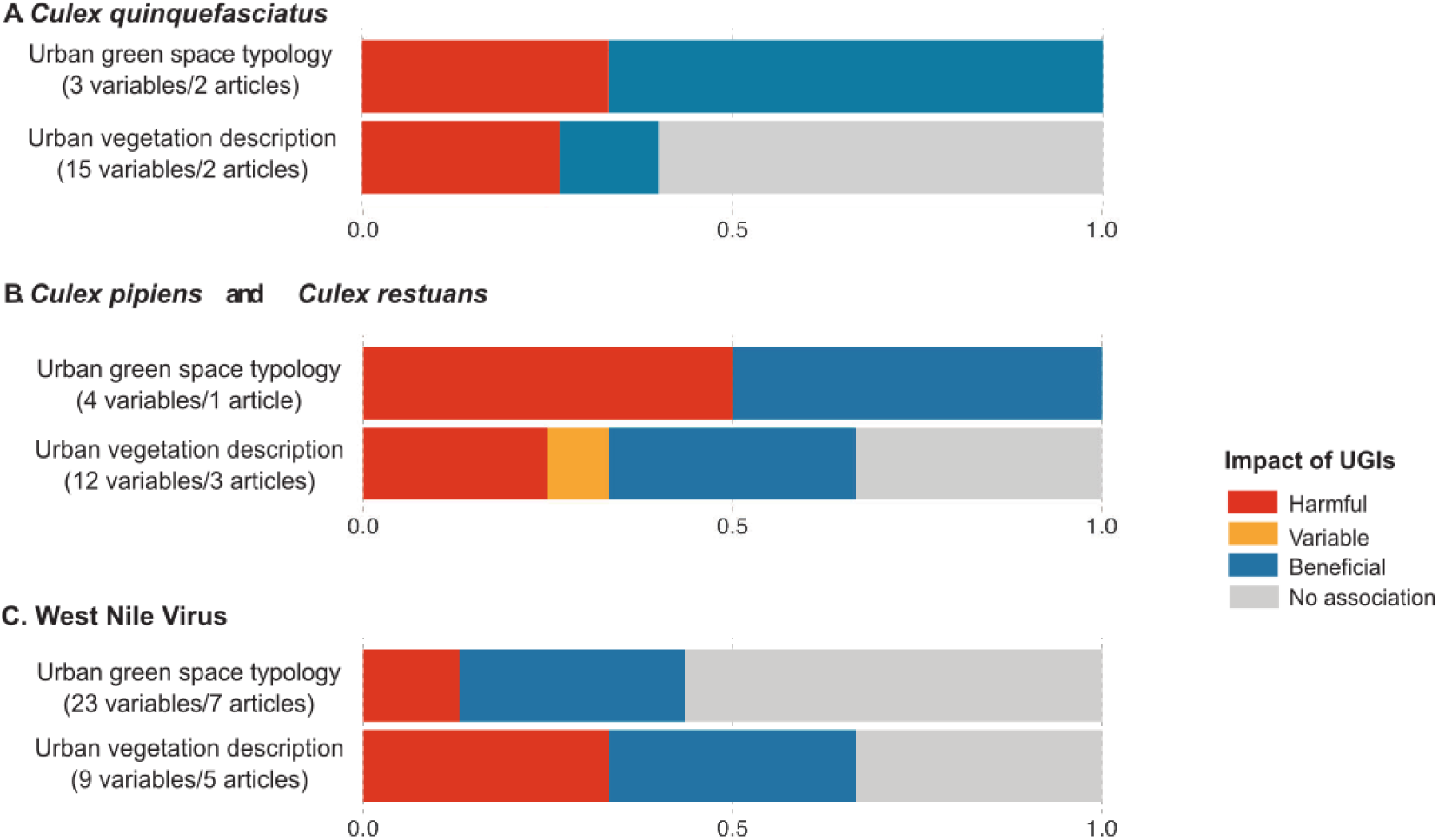
Impact of UGIs on *Culex*-pathogen system. (A) Impact of UGIs on *Culex quinquefasciatus* populations. (B) Impact of UGIs on *Culex pipiens* and *Culex restuans* populations. (C) Impact of UGIs on West Nile virus dynamics (prevalence of infection in host or vector). Numbers in brackets indicate the total number of variables tested and the total number of articles from which these variables were extracted. Since some articles provide data across both categories, they may be counted more than once, which explains discrepancies with the total number of articles reported in Table 1. The same variable tested in multiple articles is also counted multiple times. Colors represent the proportion of variables in all studies that increased risk (harmful effect, red), decreased risk (beneficial effect, blue), varied (variable effect, yellow), or showed no association (grey) with risk. The x-axis represents the proportion of UGI indicators that, for a given vector-pathogen system, were classified as harmful, beneficial, variable, or showing no association.

**Figure 5.**
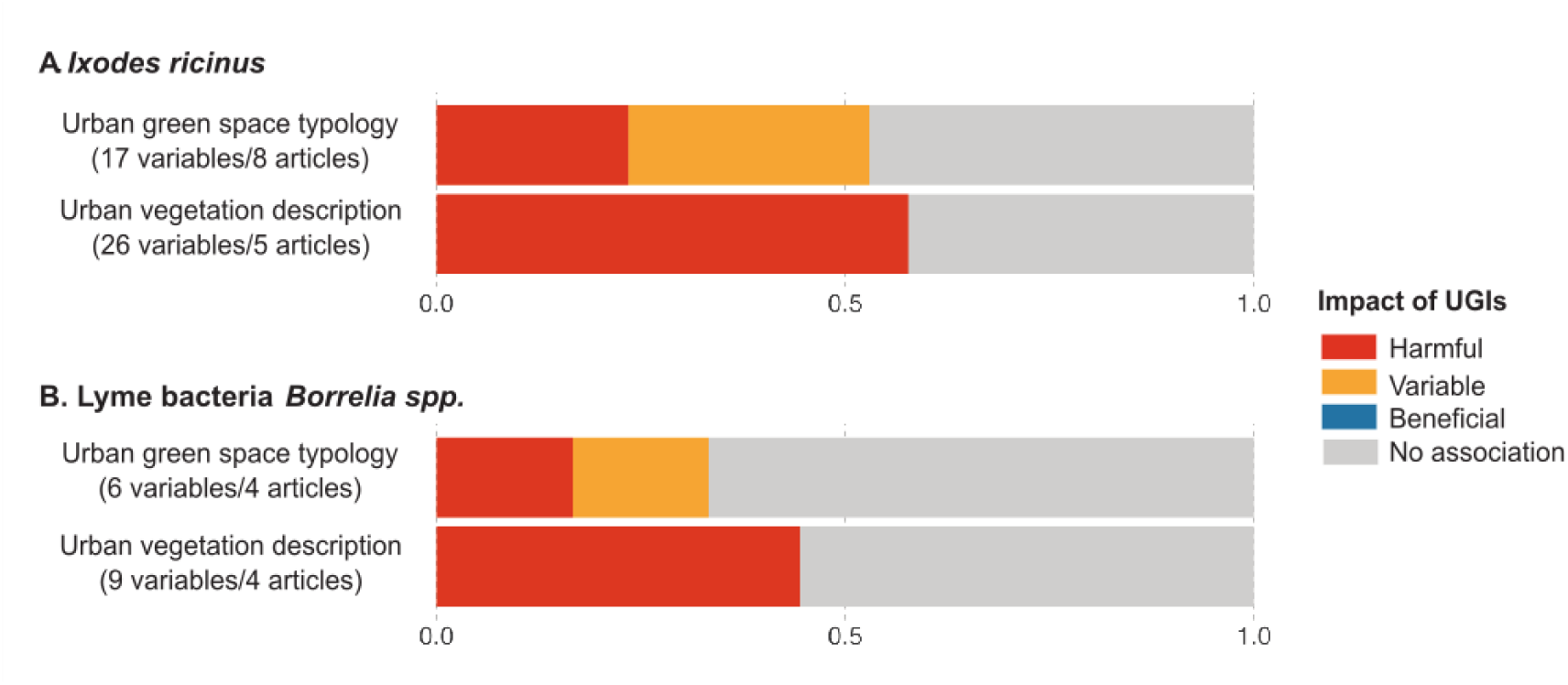
Impact of UGIs on *Ixodes*-pathogen systems. (A) Impact of UGIs on *Ixodes ricinus* populations. (B) Impact of UGIs on Lyme bacteria *Borrelia* spp. (prevalence of infection in host or vector). Numbers in brackets indicate the total number of variables tested and the total number of articles from which these variables were extracted. Since some articles provide data across both categories, they may be counted more than once, which explains discrepancies with the total number of articles reported in Table 1. The same variable tested in multiple articles is also counted multiple times. Colors represent the proportion of variables in all studies that increased risk (harmful effect, red), decreased risk (beneficial effect, blue), varied (variable effect, yellow), or showed no association (grey) with risk. The x-axis represents the proportion of UGI indicators that, for a given vector-pathogen system, were classified as harmful, beneficial, variable, or showing no association.

**Table 1:**
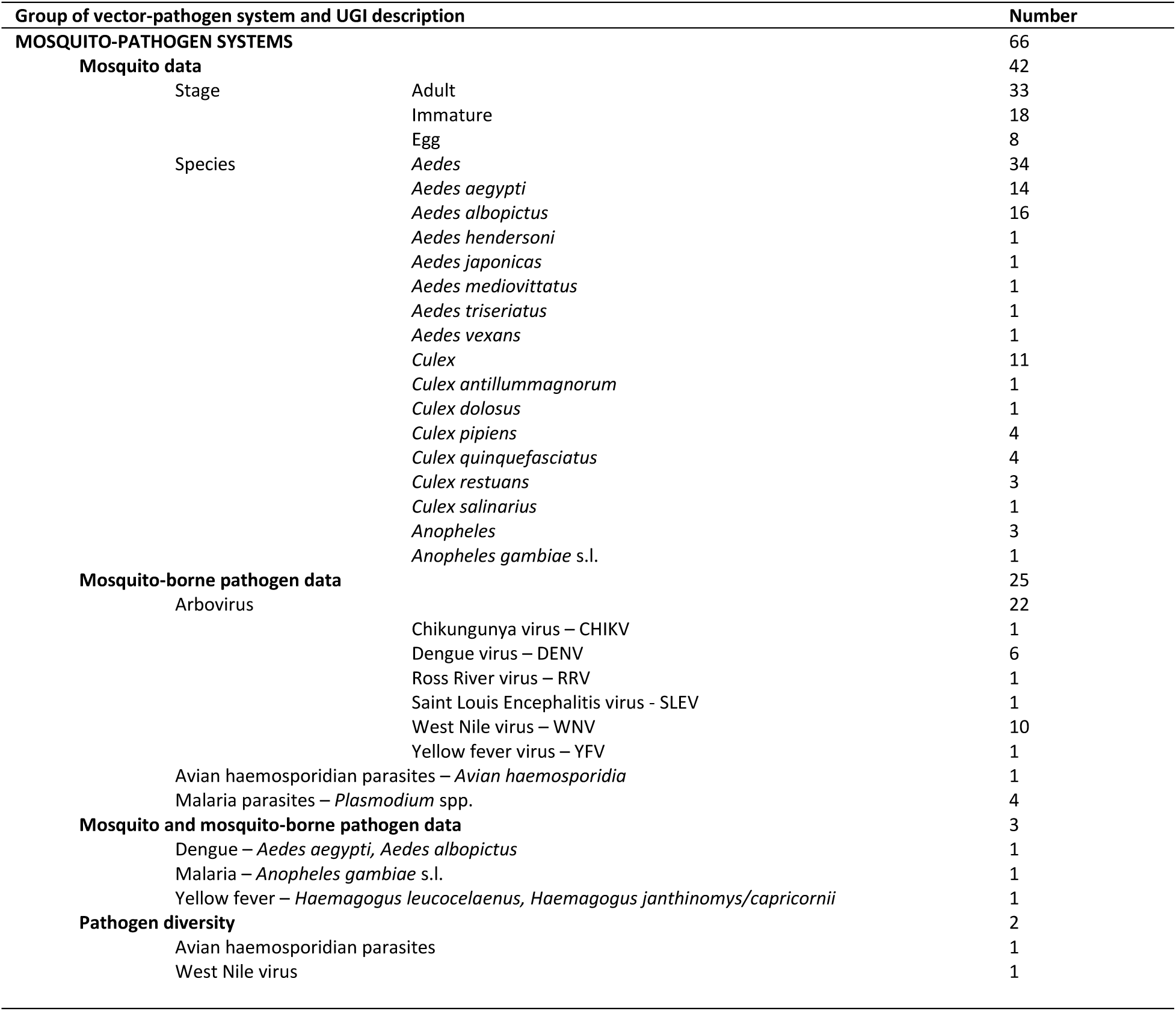
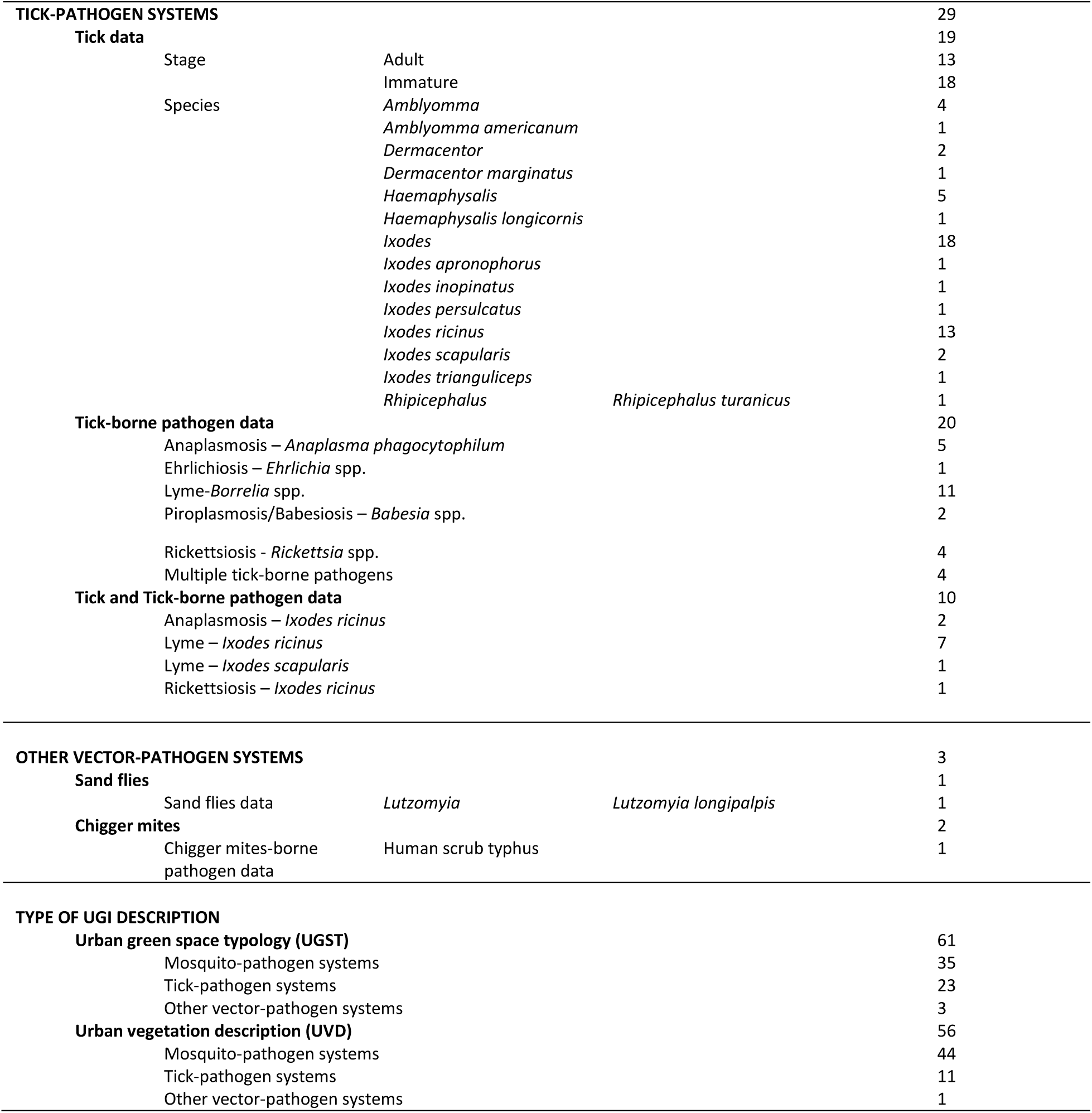
Classification of articles in accordance with the vector-pathogen system and the UGIs under investigation.

## Results

### Overview

The literature search yielded 1558 articles, of which 98 were included in the final selection after the different screening steps (Figure 1). Two-thirds of the articles focused on mosquitoes and mosquito-borne diseases or pathogens (67.3%, 66/98). One third addressed ticks and tick-borne diseases or pathogens (29.6%, 29/98). The remaining three articles examined chigger mites, sand flies, and associated pathogens. Additional file 4, Table S1, details the main results of each article, including factors associated with articles where a negative or non-association was observed with UGIs. Ninety percent of the articles featured studies conducted in a single city and country. However, studies in six articles investigated multiple cities within the same country (Levytska et al., 2021; Overzier et al., 2013; Pedrosa et al., 2020; Schorn, et al., 2011a; Schorn, et al., 2011b; Yadouléton et al., 2010), and three reported on studies carried out in cities of different countries (Carbó-Ramírez et al., 2017; Georganos et al., 2020; Talbot et al., 2021). The geographic distribution of studies varied (Figure 2A). Most mosquito-related articles originated from North America (36.3%, 24/66) and South America (28.7%, 19/66), whereas more than 70% of tick-related articles originated from Europe (72.4%, 21/29). The articles were published between 1999 and 2022, with a notable increase in mosquito-related articles after 2007 and tick-related articles after 2010 (Figure 2B).

The selected articles were categorized based on their focus: those examining vectors alone were classified under the’vector data’ category, while those studying pathogens within hosts or vectors were classified under the’vector-borne pathogen data’ category (Table 1 and Figure 2C). Among the 66 articles on mosquito-pathogen systems, 42 concentrated on mosquito data and 25 on mosquito-borne pathogens data. Only three articles addressed both mosquito and mosquito-borne pathogens data: yellow fever virus and *Haemagogus leucocelaenus* and *Haemagogus janthinomys/capricornii* (Abreu et al., 2022), dengue virus and *Aedes aegypti* (Francisco et al., 2021), and malaria and *Anopheles gambiae* s.l. (Yadouléton et al., 2010). The most studied mosquito genera were *Aedes* (34 articles), followed by *Culex* (11 articles) and *Anopheles* (three articles). Associated mosquito-borne diseases included West Nile fever (10 articles), dengue fever (six articles), and malaria (four articles). Of the 29 articles on tick pathogen systems, 19 investigated the effects of UGIs on tick vectors, 20 examined the effects of UGIs on tick-borne pathogens, and 10 addressed both categories. Ticks from the *Ixodes* genus were studied the most (18 articles), with a focus on *Ixodes ricinus* (14 articles). The most extensively studied pathogen was the *Borrelia* spp. bacterium (11 articles) and small rodents were the most frequently studied host group (seven articles).

UGIs were described using the urban green space typology (UGST) or the urban vegetation description (UVD) around the sampling points (Table 1, Box 2). Both approaches were commonly used for mosquito-pathogen systems (Table 1), appearing in 35 and 44 articles out of 66, respectively. In contrast, for tick-borne pathogen systems articles (Table 1), the urban green space typology was the dominant approach used in 23 out of 29 articles. Below, the impact of UGIs on mosquito-pathogen systems, then on tick-pathogen systems and finally on other vector-pathogen systems is examined.

### Mosquito-pathogen system

The impact of UGIs on mosquito-pathogen system is system-dependant (vector species and pathogen). To handle it, we describe results for each mosquito genus separately (i.e, *Aedes*, *Culex*, *Anopheles*). We first addressed the effects of UGIs on mosquito populations, followed by their influence on mosquito-borne pathogens, and we concluded with overarching insights into the impact of UGIs on the specific mosquito-pathogen system relationship. Only a limited number of articles examine the impact of UGIs on overall mosquito abundance without species-level differentiation. In Brazil, it has been shown that adult mosquito abundance is positively associated with urban parks in São Paulo (Wilke et al., 2017) and in Sinop (Vieira et al., 2020). The area of urban parks in São Paulo was also positively correlated with mosquito abundance (Medeiros-Sousa et al., 2017).In Malaysia, urban goat farms had more breeding sites than urban parks (Lee et al., 2020). Finally, in Tokyo (Japan), urban green roofs had less vector mosquito abundance than low elevation gardens (Wong & Jim, 2016, 2017, 2018).

#### Effects on Aedes-pathogen system

##### Effects on *Aedes* populations

Of the 34 articles studying the genus *Aede*s, 16 focused on *Aedes albopictus* and 14 on *Aedes aegypt*i (Table 1). *Aedes vexans*, *Aedes mediovittatus*, *Aedes japonicus*, *Aedes triseriatus* and *Aedes hendersoni* each appeared in a single article. In this section, we mainly describe the results for the two major disease vectors, *Ae. aegypti* and *Ae. albopictus*.

UGIs appeared to be unfavorable habitats for *Ae. aegypti* (Figure 3A). The impact of UGIs on *Ae. aegypti* was mainly assessed using UVD indicators rather than UGST indicators (Figure 3A), with most studies focusing on immature stages (78.6%, 11/14). In Brazil, the abundance of immature stages of *Ae. aegypti* was negatively correlated with vegetation cover, as observed in Rio de Janeiro (Honório et al., 2009) and São Paulo (Arduino et al., 2020). Adult *Ae. aegypti* densities were lower in urban green spaces than in highly urbanized residential areas (see Box 1), such as in Mariana and in Ouro Petro (Pedrosa et al., 2020), and lower than in built environment like asbestos-roofed structures in São Paulo (Lorenz et al., 2020).

Using UGST indicators, *Ae. aegypti* oviposition rates were observed to be higher (i) near houses compared to deeper forested areas in Rio de Janeiro (Lourenço-de-Oliveira et al., 2004), and (ii) in the peripheral part of the park with less vegetation near urban areas compared to the densely vegetated inner sections of the park (Heinisch et al., 2019). Also, other countries, including the USA (Puerto Rico) (Cox et al., 2007) and Vietnam (Huynh & Minakawa, 2022), have shown similar results. Breeding sites could be associated with residential areas, where artificial breeding containers like jars or planters were common (Huynh & Minakawa, 2022). Sun et al. (2021) observed that adult *Ae. aegypti* were more abundant in areas in Singapore with buildings and managed vegetation than in zones with dense forest cover.

Using UVD indicators, locations with a high percentage of vegetation coverage, like urban forests or urban grass in Singapore (Sun et al., 2021) or arboreal vegetation in Argentina (Benitez et al., 2020) exhibited low mosquito abundance. Moreover, in New Orleans (United States of America, USA) (de Jesús Crespo & Rogers, 2021) cemeteries with a higher heat index supported higher mosquito populations than cemeteries with lower heat index. Reduced temperature in vegetated areas may explain why *Ae. aegypti* was less commonly found in areas with more vegetation cover (de Jesús Crespo & Rogers, 2021).

Despite the general negative association between *Ae. aegypti* and UGIs, the mosquito still thrived in areas with vegetation near breeding sites, even in densely populated environments. For instance, using the UVD, Estallo et al. (2018) observed in Córdoba (Argentina) that vegetation adjacent to breeding sites was positively associated with the likelihood of these sites supporting mosquito larvae. In the same city, Andreo et al. (2021) found that the distance to vegetated areas could have a variable impact on the presence of breeding sites depending on the month, from decreasing their presence in November to increasing it in March. A study by Landau & van Leeuwen (2012) found that *Ae. aegypti* adult abundance in Tucson (USA) increased with the percentage of medium to tall trees within 10 to 50 meters of trap locations. No association was observed with herbaceous or shrub vegetation (Landau & van Leeuwen, 2012).

Contrary to *Ae. aegypti, Ae. albopictus* was generally favoured by UGIs, as indicated by both UGST and UVD indicators (Figure 3B), for both immature and adult stages. *Aedes albopictus* eggs and larvae were more abundant in urban green spaces near residential areas than in densely urbanized zones. This trend has been observed in Brazilian cities using UGST indicators, such as in Mariana and Ouro Preto (Pedrosa et al., 2020) and São Paulo (Heinisch et al., 2019; Wilke et al., 2017), as well as using UVD indicators, such as in Rio de Janeiro (Honório et al., 2009; Lourenço-de-Oliveira et al., 2004) and São Paulo (Arduino et al., 2020). Urban green spaces with higher vegetation cover favored the abundance of *Ae. albopictus* eggs, as observed in Greensboro (USA), where urban forests and urban parks yielded more *Ae. albopictus* eggs than on highly urbanized built environments, mainly composed of buildings and parking lots (Schwarz et al., 2020). Urban vegetation significantly supported the presence of breeding sites. Examples include the percentage of urban vegetation linked to *Ae. albopictus* egg abundance in Italy (Cianci et al., 2015) and the vegetation shade associated with breeding sites in Vietnam (Huynh & Minakawa, 2022). However, in the study by de Jesús Crespo & Rogers (2021) in New Orleans cemeteries, no significant link was found between the presence of vegetation next to breeding sites and the abundance of larvae. *Ae. albopictus* abundance showed only a slight non-significant increase in low-heat islands compared to high-heat islands and was less abundant than *Ae. aegypti.* The authors suggested that temperature might be the main factor influencing *Ae. albopictus* distribution there. Positive correlations between tree cover and mosquito breeding sites may be explained by specific factors. Tree holes, for example, serve as breeding sites for *Aedes* spp. mosquitoes, such as those found in *Delonix regia* and *Magnifera indica* trees in Bangladesh (Chowdhury et al., 2014), or, for *Ae. albopictus* eggs in tree canopies in Greensboro (Schwarz et al., 2020). Trees can also create a favorable microclimate for mosquito development by mitigating urban heat island effects, reducing temperatures, and providing shelter (de Jesús Crespo & Rogers, 2021). Some breeding sites, like water-filled flower vases, were associated with gardening practices (Che Dom et al., 2016). Francisco et al. (2021) also found a positive correlation between *Aedes* spp. egg abundance and the percentage of urban vegetation in the Philippines.

Regarding the abundance of *Ae. albopictus* adults, a similar trend emerged when using UGST or UVD indicators. Urban parks hosted higher adult abundances than densely residential areas, as shown in Vietnam (Huynh & Minakawa, 2022) and Brazil (Pedrosa et al., 2020). In Malaysia’s Klang Valley, urban farms had higher adult mosquito abundance than peri-urban farms (Lee et al., 2020). In Hong Kong (China), no differences in adult mosquito abundance were found between sites with green or bare roofs (Wong & Jim, 2018). Using the UVD, adult *Ae. albopictus* populations were generally more abundant in forested areas than in grassy areas, as observed in Singapore (Sun et al., 2021). Dense or poorly maintained vegetation can reduce the abundance of adults (Hendy et al., 2020; Sun et al., 2021). In Rome (Italy), Manica et al. (2016) found that adults preferred small urban green spaces, with a positive correlation between abundance and vegetation within 20 meters of the sampling trap and a negative correlation with vegetation at 300 meters. A potential explanation for the positive correlation between urban vegetation and *Ae. albopictus* adults is the role of certain plant species. Waxleaf Privet (*Ligustrum quihoui*) has been found to promote mosquito survival, while Glossy Abelia (*Abelia grandiflora*) enhanced female fecundity (Tian, 2019). However, in Manaus (Brazil), Hendy et al. (2020) reported no association between tree cover and adult abundance. The specific and context-dependent vegetation characteristics, such as the presence of certain plant species discussed here, played a critical role in mosquito abundance.

Studies examining both *Ae. albopictus* and *Ae. aegypti* in UGIs where the two species co-exist confirmed the contrasting effects on these environments on each vector (Table 2) (Arduino et al., 2020; de Jesús Crespo & Rogers, 2021; Heinisch et al., 2019; Honório et al., 2009; Huynh & Minakawa, 2022; Pedrosa et al., 2020; Sun et al., 2021). However, several articles identified similar results for both species. First, the abundance of immature stages was positively associated with vegetation adjacent to breeding sites, in Brazil (Honório et al., 2009), Bangladesh (Paul et al., 2018), Malaysia (Che Dom et al., 2016), and the Philippines (Francisco et al., 2021). A study from Rio de Janeiro reported a decline in immature stages abundance of both species with increasing distance to the peri-urban forest (Lourenço-de-Oliveira et al., 2004). Secondly, a multi-country study from South America reported that the adult abundance of both species was positively correlated with decorative vegetation and the proximity of urban vegetation to residential areas (Talbot et al., 2021).

**Table 2.** Summary of the main effects of Urban Green Infrastructures (UGIs) on *Aedes* vectors at immature and adult stages. Only variables reported in >= 2 studies are included. An “increase” in the number of immature or adult stages is classified as harmful to the risk. A “decrease” is classified as beneficial. “No association” indicates that there is no effect. “Variable” indicates that the effects vary across the same study. “No data” indicates an absence of information. UGI = Urban Green Infrastructures, UGST = Urban Green Spaces Typology, UVD = Urban Vegetation Description. Elements underlined correspond to studies addressing both vectors *Ae. aegypti* and *Ae. albopictus*.

| UGIs indicators | UGIs description | Effects on *Ae. aegypti* |  |  |  | Effects on *Ae. albopictus* |  |  |  |
| --- | --- | --- | --- | --- | --- | --- | --- | --- | --- |
|  |  | Effect on immature stages | Effects on adult stages | Countries | References | Effect on immature stages | Effects on adult stages | Countries | References |
| UGST indicators | Urban parks | Increase | Decrease | Vietnam, Brazil | (Huynh & Minakawa, 2022; Lorenz et al., 2020; Pedrosa et al., 2020) | Increase | Increase | Vietnam, Malaysia, Brazil, USA | (Huynh & Minakawa, 2022; Lee et al., 2020; Pedrosa et al., 2020; Schwarz et al., 2020) |
|  | Distance from edge within parks | Decrease | No data | Brazil | (Heinisch et al., 2019; Lourenço-de-Oliveira et al., 2004) | Increase<br>Decrease | No association | Brazil | (Heinisch et al., 2019; Hendy et al., 2020; Lourenço-de-Oliveira et al., 2004) |
| UGST & UVD indicators | Urban forest | Decrease | Decrease | USA (Puerto Rico), Singapore | (Cox et al., 2007; Sun et al., 2021) | Increase | Increase | USA, Singapore | (Schwarz et al., 2020; Sun et al., 2021) |
| UVD indicators | Vegetation cover | Increase | No association | Brazil, USA, Philippines | (Arduino et al., 2020; de Jesús Crespo & Rogers, 2021; Francisco et al., 2021; Honório et al., 2009; Landau & van Leeuwen, 2012) | Increase | Increase | Brazil, Italy, USA, Philippines | (Arduino et al., 2020; Cianci et al., 2015; de Jesús Crespo & Rogers, 2021; Francisco et al., 2021; Manica et al., 2016; Schwarz et al., 2020) |
|  |  | Decrease |  |  |  | No association | Decrease |  |  |
|  |  | No association |  |  |  |  |  |  |  |
|  | Grass | No data | Decrease<br>No association | USA, Singapore | (Landau & van Leeuwen, 2012; Sun et al., 2021) | Decrease | Decrease | Italy, Singapore | (Cianci et al., 2015; Sun et al., 2021) |
|  | Tree cover | Increase | Increase<br>No association | USA | (de Jesús Crespo & Rogers, 2021; Landau & van Leeuwen, 2012) | Increase<br>No association | No association | Italy, USA, Brazil | (Cianci et al., 2015; de Jesús Crespo & Rogers, 2021; Hendy et al., 2020) |
|  | Proximity to urban vegetation | Increase | Increase | Argentina, Brazil, Malaysia, Bangladesh | (Andreo et al., 2021; Che Dom et al., 2016; Estallo et al., 2018; Honório et al., 2009; Paul et al., 2018; Talbot et al., 2021) | Increase | Increase | Brazil, Malaysia, Bangladesh | (Che Dom et al., 2016; Honório et al., 2009; Paul et al., 2018; Talbot et al., 2021) |
|  |  | Decrease |  |  |  |  |  |  |  |
|  |  | No association |  |  |  |  |  |  |  |
|  |  | Variable |  |  |  |  |  |  |  |

Some studies have also explored the effects of UGIs on other *Aedes* species. For instance, *Aedes mediovittatus* larvae were more abundant in low-density residential areas, with over 20% vegetation cover, compared to forest patches and densely urbanized residential areas in Puerto Rico (Cox et al., 2007). In New Haven (USA), more *Aedes vexans* adults were found in moderately vegetated residential areas than in highly or un-vegetated areas (Brown et al., 2008). For *Aedes japonicus*, adult abundance showed no difference between green and bare roofs in Hong Kong (Wong & Jim, 2018). *Aedes triseriatu*s and *Aedes hendersoni* eggs were more abundant in urban forests than on urban campuses in Greensboro (Schwarz et al., 2020).

##### Effects on dengue virus dynamics

Dengue is an arboviral disease in which the virus (DENV) is transmitted primarily by *Aedes* mosquitoes, particularly *Ae. aegypti* and *Ae. albopictus*. Six articles examined the relationship between dengue incidence and UGIs (Table 1), revealing varying effects across locations and periods (Figure 3C). Only one of them used UGST indicators and observed that the impact of urban green spaces on dengue incidence in humans varied by city in China, with effects ranging from no association to increased risk (Chen et al., 2020). Using UVD indicators, the same authors found that dengue incidence was positively correlated with forest coverage in two Chinese cities (Chen et al., 2020). The other five studies reported a weak negative correlation between vegetation cover and human dengue incidence. This was observed in Nouméa (New Caledonia) (Zellweger et al., 2017), São Paulo (Araujo et al., 2015), and Manila (the Philippines) (Francisco et al., 2021). In other locations, such as Puntarenas (Costa Rica) (Troyo et al., 2009), and São Paulo (Ogashawara et al., 2019), there was no association found.

##### Effects on Chikungunya virus dynamics

Chikungunya virus (CHIKV) is primarily transmitted by *Aedes* mosquitoes, particularly *Ae. aegypti* and *Ae. albopictus*. To date, only one study explored the link between UGIs and CHIKV dynamics and showed that, in Rio de Janeiro, where *Ae. aegypti* serves as the vector, a higher proportion of green parks was associated with reduced CHIKV incidence in humans (Freitas et al., 2021).

##### Concluding remarks on *Aedes*–pathogen systems

In summary, the effects of UGIs on *Aedes*-pathogen systems vary between the two main arbovirus vectors (Table 2). *Ae. aegypti* generally shows a negative response to UGIs, while *Ae. albopictus* tends to be favored by them. When it comes to arboviruses, particularly DENV, the evidence is less consistent. This likely reflects the fact that DENV is transmitted by both species, which respond differently to urban green environments. Consequently, identifying the dominant vector in a given setting is crucial for accurately assessing dengue risk.

#### Effects on Culex-pathogen systems

##### Effects on *Culex* populations

Of the 11 articles reporting on the effect of UGIs on *Culex* mosquitoes, four focused on *Culex quinquefasciatus* (Cox et al., 2007; Landau & van Leeuwen, 2012; Sallam et al., 2017; Wong & Jim, 2018), four on *Culex pipiens* (Brown et al., 2008; DeMets et al., 2020; Gardner et al., 2013; Trawinski & Mackay, 2010) and three on *Culex restuans* (Brown et al., 2008; DeMets et al., 2020; Gardner et al., 2013) (Table 1, Figure 4A and Figure 4B). The main results are summarised in Table 3.

**Table 3.**
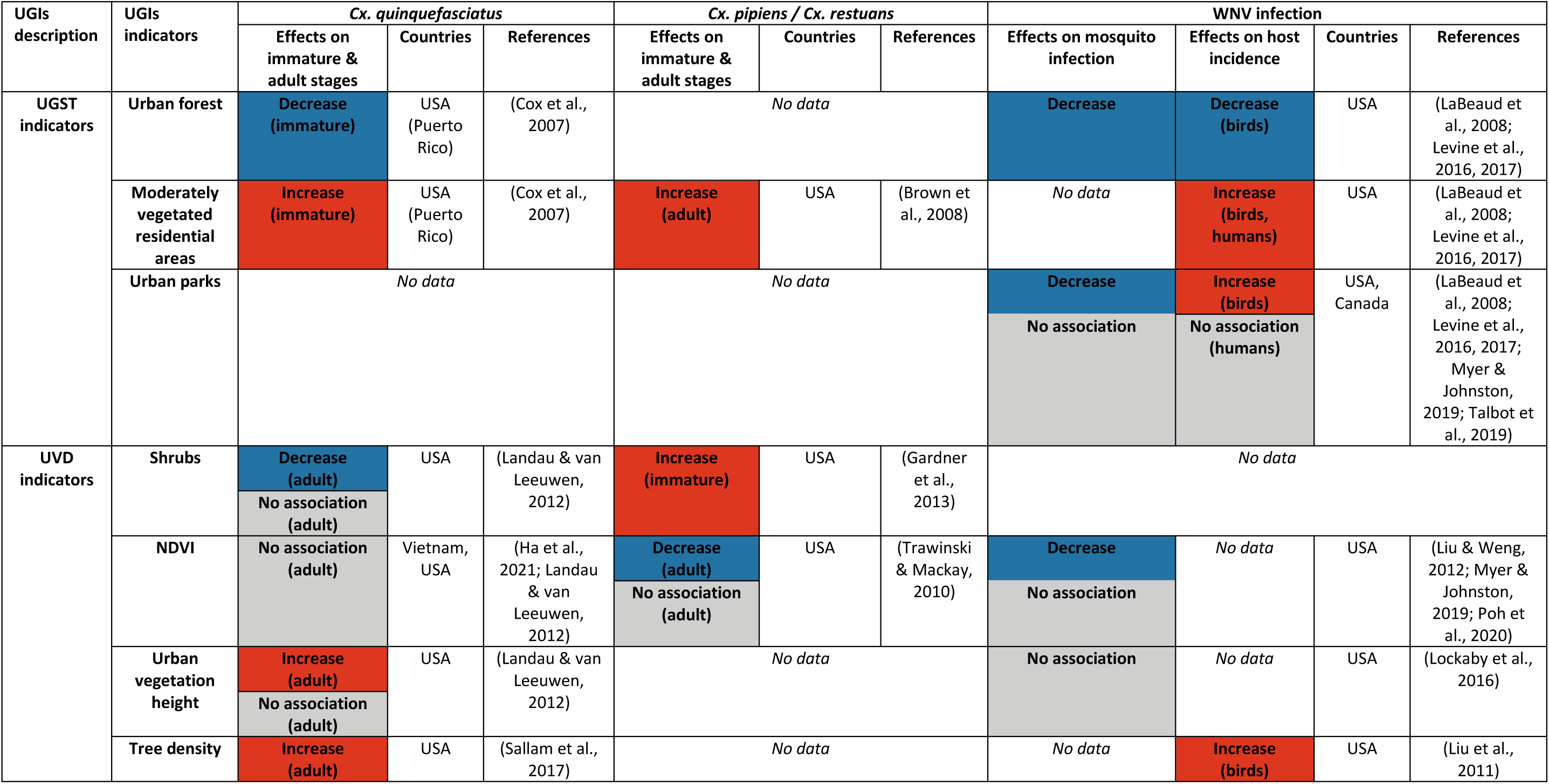
Summary of the main effects of UGIs on the *Culex*-pathogen system, including mosquito abundance and WNV incidence in mosquitoes and hosts. Only variables reported in ≥ 2 studies are included. An “increase” in the number of immature or adult stages is classified as harmful to the risk. A “decrease” is classified as beneficial. “No association” indicates that there is no effect. “Variable” indicates that the effects vary across the same study. “No data” indicates an absence of information. Abbreviations: UGI = Urban Green Infrastructures; UGST = Urban Green Space Typology; UVD = Urban Vegetation Description.

Regarding *Cx. quinquefasciatus,* UGST variables showed higher abundance of larvae in low-density residential areas with low vegetation cover compared to urban forests or densely urbanized residential areas in Puerto Rico (Cox et al., 2007) (Figure A). A study in Hong Kong reported fewer males on green roofs than on bare roofs (Wong & Jim, 2018). For UVD indicators (Figure 4A), adult abundance in Tucson (Landau & van Leeuwen, 2012) and in New Orleans (Sallam et al., 2017), was generally not influenced by urban vegetation. Some studies observed positive correlations with medium and high trees (Landau & van Leeuwen, 2012), and with urban tree density (Sallam et al., 2017), while negative associations were reported with herbaceous and shrub cover, depending on buffer distances around sampling points (Landau & van Leeuwen, 2012).

The impact of UGIs on *Cx. pipiens* and *Cx. restuans* did not reveal any clear trend as well (Figure 4B). Urban vegetation, measured indirectly by NDWI, was positively correlated with the abundance of *Cx. pipiens* and *Cx. restuans* in Toronto (Canada) (DeMets et al., 2020), but showed the opposite effect or no correlation in Amherst (USA) (Trawinski & Mackay, 2010). DeMets et al. (2020) noted that NDVI was not associated with the abundance of *Cx. pipiens* and *Cx. restuans*. In Chicago (USA), non-flowering shrubs and specific tree species, such as oak and pear, favored the abundance of immature stages of *Cx. pipiens* and *Cx. restuans*, while flowering shrubs and ash trees had the opposite effect (Gardner et al., 2013). Additionally, deciduous tree density showed no impact on abundance. When considering urban green spaces within residential areas, *Cx. pipiens* and *Cx. restuans* preferred moderately vegetated residential areas over residential areas with sparse or dense vegetation, as observed in New Haven (USA) (Brown et al., 2008).

Some studies have also explored the effects of UGIs on other *Culex* species. *Culex salinarius* females were more abundant in highly vegetated residential areas than in sparsely or moderately vegetated areas in New Haven (Brown et al., 2008). In Rio de Janeiro, *Cx. dolosus* was more commonly found deeper in urban forests than near residential areas (Lourenço-de-Oliveira et al., 2004). Similarly, *Cx. antillummagnorum* in Puerto Rico was more abundant in urban forests and low-density residential areas, with over 20% vegetation cover, than in densely urbanized residential areas (Cox et al., 2007). Additionally, in Hanoi (Vietnam), the abundance of female *Cx. vishnui* and *Cx. tritaeniorhynchus* was higher in urban rice fields than in urban forest areas (Ha et al., 2021). A study in Cleveland (USA) examined *Culex* mosquitoes without identifying them to species level (Yang et al., 2019). In this study, adult mosquito abundance was positively correlated with grass and shrub cover, while tree canopy coverage had no significant effect. The same study found that vegetation diversity, biomass, and the presence of vacant lots did not influence *Culex* abundance.

##### Effects on West Nile virus dynamics

West Nile virus (WNV) is primarily transmitted by *Culex* spp. mosquitoes and circulates within bird populations, particularly among corvids (crows, blue jays, etc.) (Chancey et al., 2015; Levine et al., 2016, 2017; Liu et al., 2011). Humans and horses serve as dead-end hosts. Ten articles examined the effects of UGIs on WNV dynamics, all of which were conducted in North America (Table 1). The impact of UGIs on WNV dynamics was mainly explored using UGST indicators compared to UVD indicators (Figure 4C). UGIs were often associated with negative or no effects on WNV dynamics (Table 3).

Eight of the ten studies examined the effect of UGIs on WNV infection in *Culex* spp. (Levine et al., 2016, 2017; Liu et al., 2008; Liu & Weng, 2012; Lockaby et al., 2016; Myer & Johnston, 2019; Poh et al., 2020; Talbot et al., 2019). Only urban agriculture showed a positive association with mosquito infection, as shown in Indianapolis (USA) (Liu et al., 2008), likely due to favorable microclimatic conditions. Four studies (three using UGST and one using UVD indicators) found no association between mosquito infection and urban green spaces, such as in Ottawa (Canada) (Talbot et al., 2019) and Atlanta (USA) (Levine et al., 2016, 2017), or with vegetation measured by NDVI, as in Los Angeles (USA) (Liu & Weng, 2012). The absence of an NDVI effect in Los Angeles may reflect a cross-correlation with higher elevation. Negative associations were reported in Atlanta (Lockaby et al., 2016), New York city (USA) (Myer & Johnston, 2019), and Dallas (USA) (Poh et al., 2020) where mosquito infection was inversely related to NDVI, pine forest cover (linked to dryness), and large forest patches, possibly reflecting a dilution effect through higher avian diversity.

Five studies investigated the association between WNV incidence or seroprevalence in hosts and UGIs, focusing on birds and humans (Figure 4C). WNV seroprevalence was higher in residential areas than in more vegetated areas for both birds (Levine et al., 2016, 2017) and humans (LaBeaud et al., 2008). In Atlanta, bird seroprevalence was also higher in urban wooded parks than in urban forests (Levine et al., 2016, 2017), as in Cleveland (USA) (LaBeaud et al., 2008). In Northern Virginia (USA), medium to high canopy density was positively associated with WNV incidence in corvids (Liu et al., 2011), whereas in Ottawa (Canada), no association was found between park proportion and human WN case incidence (Talbot et al., 2019). Two studies on WNV genetic diversity (Amore et al., 2010; Bertolotti et al., 2008) showed stability or decline in mosquitoes within urban green spaces, but higher diversity in birds compared to residential areas.

##### Effects on other *Culex*-borne pathogens

A study in Argentina found that prevalence in humans of Saint Louis encephalitis virus, transmitted by *Cx. quinquefasciatus*, was lower in low-density residential areas than in densely populated regions (Vergara Cid et al., 2013). In Brisbane (Australia), incidence of Ross River virus in humans, a virus transmitted by *Aedes vigilax* and *Culex annulirostris*, was positively associated with the percentage of riparian and bushland areas (Muhar et al., 2000).

##### Concluding remarks on *Culex*-pathogen systems

The diversity and specificity of UGI indicators made it challenging to identify consistent trends in their impacts on *Culex*-pathogen systems. Overall, our synthesis showed that *Cx. quinquefasciatus, Cx. pipiens,* and *Cx. restuans*, and WNV host seroprevalence were generally higher in moderately vegetated residential areas than in urban forest or areas with dense vegetation (Table 3). Such residential areas with moderate vegetation cover could potentially act as hotspots for mosquito abundance and WNV transmission. In contrast, urban forests appeared to reduce *Cx. quinquefasciatus* populations and WNV host incidence, while high tree density tended to increase mosquito abundance and bird infection rates. Other UGST and UVD indicators showed no influence or negative associations with mosquito infection. The dynamics of WNV transmission remain complex and are influenced by various factors, including plasticity in the feeding behavior of mosquitoes, host availability, and diversity in the ability of bird host species to amplify and further transmit the virus to the vector mosquitoes (Levine et al., 2016, 2017).

##### Effects on Anopheles-pathogen systems

The *Anopheles* spp. mosquitoes are the vectors of *Plasmodium* spp., the parasites that cause malaria. Only three of the selected articles focused on *Anopheles* spp. mosquitoes, all indicating a positive correlation between urban agriculture and the presence or abundance of *Anopheles* mosquitoes, and mainly using UGST indicators (Dongus et al., 2009; Matthys et al., 2010; Yadouléton et al., 2010). In Tanzania (Dongus et al., 2009) and Côte d’Ivoire (Matthys et al., 2010), the presence of wells and agricultural trenches increased the likelihood of *Anopheles* presence and its density. The proportion and proximity of urban agriculture, such as vegetable farms, were positively associated with *Anopheles* populations in Tanzania (Dongus et al., 2009) and, in Benin, the population of *Anopheles gambiae stricto senso* (Yadouléton et al., 2010). Similar patterns emerged in the four articles addressing malaria dynamics, using UGST (Matthys et al., 2006; Yadouléton et al., 2010) and UVD indicators (Georganos et al., 2020; Kabaria et al., 2016). Both mosquito infection rates and human malaria prevalence were positively linked to urban agricultural areas.

##### Effects on Haemagogus and on yellow fever dynamics

In Rio de Janeiro, Abreu et al. (2022) studied the relationships between UGIs and the yellow fever vector system, which involves the vectors *Haemagogus leucocelaenus* and *Haemagogus capricornii*, as well as human and non-human primates as hosts. The study found no difference in vector abundance between urban forests, urban pastures, and densely urbanized areas, but the abundance of vectors was positively associated with the NDVI. Furthermore, the prevalence of yellow fever in humans and non-human primates was higher in urban forest fragments than in heavily urbanized areas. Both *Hg. leucocelaenus* and *Hg. capricornii* showed higher infection rates for yellow fever virus (YFV) in urban forest settings compared to other urban areas.

##### Effects on other mosquito-pathogen systems

Only four studies evaluated the influence of UGIs on the abundance of all mosquitoes collected. In São Paulo, Medeiros-Sousa et al. (2017) found a positive association between mosquito abundance and urban parks. Another study in Sinop (Brazil) reported higher abundance of adult mosquitoes in urban forested parks and in urban forest fragments than in urbanized areas (Vieira et al., 2020). In Hong Kong, the abundance of all adult species was lower on green roofs compared to bare roofs (Wong & Jim, 2016) and low-elevation gardens (Wong & Jim, 2017). One multi-site study indicated that, in both Mexico and Germany, there was no association between seroprevalence of avian haemosporidia in birds and urban forests (Carbó-Ramírez et al., 2017).

##### Concluding remarks on the impact of UGIs on mosquito-pathogen systems

To sum up, the influence of UGIs on mosquito populations and mosquito-borne diseases is context-specific, depending on the vector-pathogen system studied and on the mosquito species involved. For the *Aedes*-pathogen system, *Ae. aegypti* abundance was mainly negatively correlated with UGIs, whereas *Ae. albopictus* was mainly positively associated, with no clear trend for DENV. For the *Culex*-pathogen system, *Culex* spp. abundance was generally unaffected or negatively influenced by UGIs, except in low-vegetation residential areas, where WNV host infection was higher than in more vegetated areas. Positive associations between urban agriculture and *Anopheles*-*Plasmodium* dynamics were also observed. Further research is urgently needed to clarify the effects of UGIs on other mosquito vector populations dynamics and associated mosquito-borne disease risk in cities (Additional file 5: Table S2, Additional file 6: Table S3).

### Effects of UGIs on tick-pathogen systems

Few studies have examined the effects of UGIs on all tick species. Considering all species, the UGST and UVD indicators mainly showed either an increase in tick populations or no effect. In the USA, urban parks and trails were positively correlated with tick abundance in New York (Hassett et al., 2022),as was the proportion of surrounding undeveloped parks in Oklahoma (Roselli et al., 2022). In contrast, in Cluj-Napoca, Romania, urban parks and vegetation cover were not associated with tick populations (Borşan et al., 2020). For each tick-pathogen system, we first addressed the effects of UGIs on ticks populations, followed by their influence on tick-borne pathogens, and we concluded with overarching insights into the impact of UGIs on the specific tick-pathogen system relationship.

#### Effects on Ixodes-pathogen system

##### Effects on *Ixodes* populations

Across 19 articles on UGIs and tick populations, thirteen articles centered on *Ixodes ricinus* in Europe and two on *Ixodes scapularis* in North America (Gregory et al., 2022; VanAcker et al., 2019). Six of the thirteen articles observed a positive correlation between UGIs and *I. ricinus* population density, with one using UGST indicators and five using UVD indicators (Figure 5A). Higher tick densities may be linked to greater connectivity between urban and peri-urban environments, allowing movement of the host, as observed in Antwerp (Belgium) (Heylen et al., 2019). Using UVD indicators, *I. ricinus* was more abundant at woodland edges and within forests than in grasslands, as demonstrated in Salisbury (United Kingdom, UK) (Hansford et al., 2017), London (UK) (Hansford et al., 2021), and Rome (Di Luca et al., 2013), as well as in urban parks in Bath (UK) (Hansford et al., 2022). Additionally, *I. ricinus* was more prevalent in deciduous or mixed forests than in coniferous forests, as noted in Poznań (Poland) (Michalik et al., 2003). This habitat preference can be explained by the presence and the abundance of small mammals (e.g., hedgehogs, *Apodemus* spp.) (Borşan et al., 2020; Michalik et al., 2003) and larger mammals such as white-tailed deer (Hansford et al., 2017) which serve as hosts for ticks.

In contrast, eight articles reported no consistent trend between UGIs and tick population density (Figure 5A), with seven articles using UGST indicators and one using UVD indicators. Studies conducted in Helsinki (Finland) (Junttila et al., 1999), Cluj-Napoca (Romania) (Borşan et al., 2020), and Warsaw (Poland) (Welc-Falęciak et al., 2014) revealed differences in *I. ricinus* abundance between urban green spaces, like urban parks, though these studies did not explain the observed variations. A study conducted in Lyon (France) (Mathews-Martin et al., 2020) showed higher tick abundance in the peri-urban park than in the urban parks. Other articles conducted in St Petersburg (Russia) (Tretyakov, 2009) and in Olstzyn (Poland) (Kubiak et al., 2019) found no significant differences in vector abundance across urban parks and urban forests. In Budapest (Hungary), Hornok et al. (2014) observed that tick abundance could vary by the month of sampling, sometimes being higher in cemeteries than in urban forests or parks or showing no differences. Finally, a study in Bath revealed no differences in tick vector abundance between woodland and grassland or between mixed and deciduous forests (Hansford et al., 2022).

In the two articles on *I. scapularis* in the city of New York (USA), urban green space connectivity and host accessibility were important in determining tick population density. Enhanced connectivity, measured by park-to-park links (VanAcker et al., 2019), was correlated with increased tick numbers. Permeable garden edges did not seem to raise the probability of tick detection (Gregory et al., 2022). High canopy cover around parks (VanAcker et al., 2019) or gardens (Gregory et al., 2022) was associated with increased tick counts, while lower tick numbers were found in grasslands or bare soil.

Other *Ixodes* species, such as *I. persulcatus, I. trianguliceps,* and *I. apronophorus*, have been observed in urban parks, such as in St. Petersburg (Tretyakov, 2009). In Hanover (Germany), Hauck et al. (2020) reported that *I. ricinus* and *I. inopinatus* densities were higher in urban mixed forests than in urban parks, with no difference between deciduous and mixed forests and between mixed forests and urban parks.

##### Effects on *Borrelia* spp. dynamics

Eleven articles focused on *Borrelia spp.* (Table 1), the bacterial pathogens causing Lyme disease, primarily transmitted by *Ixodes* ticks. The presence of infected ticks is closely linked to the movement and presence of small mammals, such as the striped field mouse (*Apodemus agrarius*), a rodent found in urban areas, which can be infected by *Borrelia miyamotoi* and may contribute to pathogen circulation (Gryczyńska et al., 2021). The effects of UGIs on tick infection rates were primarily studied using UVD indicators compared to UGST indicators (Figure 5B).

Five articles reported a positive correlation between UGIs and *Borrelia* dynamics, with two relying on UGST and three on UVD indicators (Figure 5B). In Budapest (Hungary), the *Ixodes* infection rates were higher in urban forests than in cemeteries (Hornok et al., 2014). In New York city, VanAcker et al. (2019) demonstrated that higher connectivity among urban green spaces— measured by a park’s contribution to network connectivity—increased tick infection rates. Patterns of the effects of UVD indicators on *Borrelia* infection closely aligned with the impact on *Ixodes* tick abundance. Infection rates were higher in urban woodlands compared to grasslands in London (Hansford et al., 2021) and at woodland edges compared to grasslands in Bath (Hansford et al., 2022). In New Castle county (USA), *Rosa multiflora* presence in urban forests was associated with higher infection rates in ticks (Adalsteinsson et al., 2018), as was woody debris compared to leaf litter.

Seven articles, including four using UGST and three employing UVD indicators, found no link between UGIs and *Borrelia* dynamics (Figure 5B). Urban parks were not significantly associated with *Borrelia* spp. infection rates in ticks in some European cities including Prague (Czech Republic), Helsinki, Budapest and Olsztyn (Hornok et al., 2014; Junttila et al., 1999; Kubiak et al., 2019; Kybicová et al., 2017). There was no significant difference in ticks’ infection rates between grasslands and woodlands in Salisbury (Hansford et al., 2017) and in Bath (Hansford et al., 2022), or between coniferous and deciduous forests in Poznań (Michalik et al., 2003).

##### Effects on other tick-borne pathogens dynamics

Four studies focused on the *Rickettsia/I. ricinus* vector-pathogen system in German cities (May & Strube, 2014; Overzier et al., 2013; Schorn, et al. 2011a) and Polish cities (Welc-Falęciak et al., 2014). They found no differences in ticks’ infection rates within urban green spaces, without offering specific explanations for these results. Five studies investigated the relationship between UGIs and the *Anaplasma phagocytophilum/I. ricinus* system. Infection rates for *An. phagocytophilum* varied within and between German cities’ green spaces (May & Strube, 2014; Schorn et al., 2011b). However, studies from Hungary and Poland (Hornok et al., 2014; Welc-Falęciak et al., 2014) found no significant differences between different urban green spaces. Infection rates were higher in ticks in urban parks than in peri-urban parks in Prague (Kybicová et al., 2017) and higher in urban parks than in cemeteries in Budapest (Hornok et al., 2014). One study explored the *Babesia/I. ricinus* system and reported no association between ticks’ infection rates and urban green spaces in Germany (Schorn, et al. 2011a).

Two studies conducted in Ukraine (Levytska et al., 2021) and in Belgium (Heylen et al., 2019) examined multiple pathogens (*Rickettsia spp.*, *Babesia spp.*, *Bartonella spp.*, and *Anaplasma phagocytophilum*) collectively. The findings indicated that infection rates with these tick-borne pathogens were not affected by urban green spaces or their connectivity.

##### Concluding remarks on *Ixodes*-pathogen systems

The abundance of *Ixodes* ticks was influenced by host movement, likely shaped by connectivity between urban green spaces and by local vegetation characteristics, with ticks exhibiting a preference for deciduous or mixed forests. The main results are summarised in Table 4. Variations in *Ixodes* tick abundance across different urban green spaces were location-specific and depended on the time of year. *Borrelia* infection rates in ticks were strongly linked to the presence and movement of competent hosts, particularly in woodlands or connected urban green spaces, reflecting the influence of UGIs on *Ixodes* populations. However, the effect of urban green spaces on ticks’ infection rate varied by location and specific characteristics, which may explain why some UGIs have minimal impact. For other tick-borne pathogens, infection rates in ticks did not exhibit any consistent pattern in relation to UGIs.

**Table 4.**
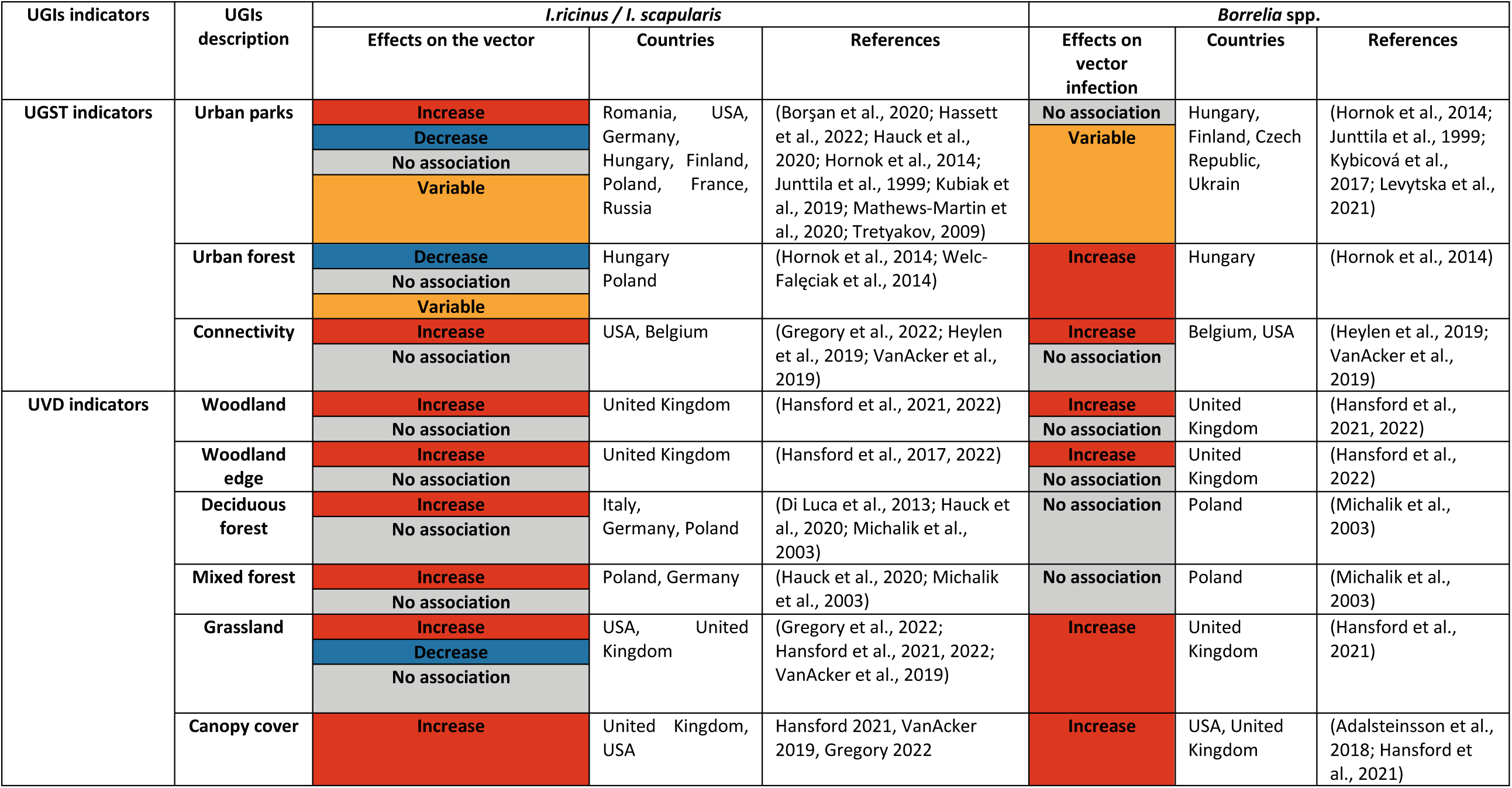
Summary of the main effects of UGIs on the *Ixodes*-pathogen system, including tick abundance and *Borrelia* tick infection. Only variables reported in ≥ 2 studies are included. An “increase” in the number of immature or adult stages is classified as harmful to the risk. A “decrease” is classified as beneficial. “No association” indicates that there is no effect. “Variable” indicates that the effects vary across the same study. “No data” indicates an absence of information. Abbreviations: UGI = Urban Green Infrastructures; UGST = Urban Green Space Typology, UVD: Urban Vegetation Description.

### Effects on Amblyomma-pathogen system

Only one study focused on *Amblyomma americanum*. This tick was positively associated with urban green spaces, particularly with permeable garden edges and increased canopy cover around yards in New York city (Gregory et al., 2022). The presence of log and brush piles in the yard also favored the probability of tick presence, which is not influenced by the presence of low canopy and grass around the yard.

Two studies reported results on other *Amblyomma*–pathogen systems. In Oklahoma (USA), infection rates of *A. americanum* with *Rickettsia* spp. or *Ehrlichia* spp. were comparable between residential and non-residential areas (Small & Brennan, 2021). Additionally, Noden et al. (2022) found that the percentage of developed land surrounding sampling points did not affect infection rates of *A. americanum* with *Borrelia* spp., *Rickettsia* spp., or *Ehrlichia* spp. Altogether, these results highlighted that *Amblyomma* tick populations were positively associated with the presence of urban green spaces, without evidence of differences in pathogen infection rates.

#### Effects on other tick species populations

Only one study focused on *Haemaphysalis longicornis* in New York city, showing that its presence was positively associated with logs and brush piles in yards, while it was unaffected by the permeable edge or the surrounding land cover of sampled yards (Gregory et al., 2022). In Italy, *Dermacentor marginatus* and *Rhipicephalus turanicus* were more prevalent in open landscapes, including pastures and wheat fields, than in deciduous woodlands (Di Luca et al., 2013).

Four studies explored the relationship between UGIs and various tick species from different genera. In Romania, *I. ricinus* and *Haemaphysalis punctata* were found to be unaffected by vegetation cover (Borşan et al., 2020). In Singapore, Kwak et al. (2021) observed higher infestation rates of small mammals by ticks such as *Amblyomma helvolum*, *Dermacentor auratus*, *Haemaphysalis semermis*, and *Ixodes granulatus* in old secondary forests compared to young secondary forests, with no influence from scrubland. In Oklahoma, Roselli et al. (2022) found that bird infestations by ticks, including *A. americanum*, *Amblyomma maculatum*, and *Haemaphysalis leporispalustris*, were not affected by the surrounding land cover of park sampling sites. In New York City, Hassett et al. (2022) reported that the densities of *I. scapularis*, *Amblyomma americanum*, and *Haemaphysalis longicornis* were negatively influenced by park management practices and were higher along trails than in park edges. Human-tick contact was also more frequent on trails than in natural open spaces. These results revealed no consistent trends across tick species, although park management practices were thought to reduce overall tick abundance.

#### Concluding remarks on other tick-pathogen systems

We found no record of negative effects of UGIs on tick populations. UGIs allowed the presence of ticks in cities and had either no or a positive effect on tick populations and the transmission of tick-borne diseases (Figure 3, Additional file 7: Table S4, Additional file 8: Table S5). Tick abundance and the prevalence of infected ticks were primarily determined by the movement of competent hosts between urban green spaces and peri-urban environments, as well as by local vegetation characteristics. Further research is needed to examine the effects of UGIs on the risk of tick-borne diseases other than Lyme disease in urban areas.

#### Effects of UGIs on others vector-pathogen systems

Only one study assessed the impact of UGIs on sand fly populations. In Correntes (Argentina), abundance of *Lutzomyia longipalpis* was higher in areas with low NDVI and in presence of farm animals compared to areas with high NDVI (Berrozpe et al., 2017).

Two studies in Asia investigated chigger mite vectors (Trombiculidae), known carriers of *Orientia tsutsugamushi*, the bacterium responsible for scrub typhus. Park et al. (2015) reported that human scrub typhus prevalence was higher in urban parks than in residential areas in South Korea. Additionally, in Thailand, rodent (*Rattus rattus*) infestation rates were higher in peri-urban and large parks than in small parks in urban centers and were negatively associated with distance from open fields (Wulandhari et al., 2021). This suggests that the environmental characteristics of urban parks may influence vector abundance and that proximity to open fields may increase vector populations (Wulandhari et al., 2021).

It is noteworthy that, due to the lack of study, no conclusive trend can be drawn concerning the relationships between UGIs and other vector-pathogen systems, including some of major public health concerns.

## Discussion

Since the early 2000’s, a growing body of research has emphasized the need to investigate the relationship between UGIs and the risk of VBD introduction and expansion. While urban green spaces may result in increased VBD emergence and transmission for city-dwellers, they also hold potential for VBD risk mitigation through careful UGI planning and management (Fournet et al., 2024). This review highlighted the diverse impacts of UGIs and urban vegetation features on arthropod vectors of human and animal diseases and associated VBD dynamics in cities. These impacts varied significantly depending on vector system and the local socio-ecological and epidemiological context.

The selection of databases and keywords may have resulted in the exclusion of certain articles. For instance, one study lacking the specified keywords was identified through cross-referencing, and was added to the 97 selected articles. However, the geographic and thematic heterogeneity of available studies underscores the need for more comprehensive and systematic sampling and monitoring of interactions between UGIs, disease vectors and VBDs affecting humans, but also animals. This should be achieved through an integrative One Health approach, as promoted by the UN in June 2021, which aims to sustainably balance and optimise the health of people, animals and ecosystems (OHHLEP, 2022). Such an approach should consider local sociological, political, and ecological contexts and use harmonized protocols and robust theoretical frameworks for data acquisition and analysis. This will help build scientific evidence and promote stakeholders’ engagement to mitigate VBD risk in green cities. Below, we discuss the main findings by geographical region, highlight key challenges, and propose research avenues to advance this field.

### Main results of the scoping review by geographical region

Direct comparisons across cities with contrasting landscapes and climates were challenging, but we present the main results stratified by vector species and geographical regions. Most *Aedes* studies were conducted in South and North America, predominantly in areas with hot and wet summers and mild winters in the USA, except for Tucson, which was really hot and dry, and Amherst, which had a cold winter. In Brazil and Argentina, the cities studied were located in areas with subtropical and equatorial climates. For *Ae. aegypti*, UGIs were negatively associated with vector presence, except in Tucson, where positive associations with low tree height were reported (Landau & van Leeuwen, 2012), and in New Orleans, where tree cover was positively associated with breeding sites (de Jesús Crespo & Rogers, 2021). For *Ae. albopictus*, UGIs were mostly positively associated regardless of the climate context (Table 2). The negative association reported by Manica et al. (2016) was linked to the use of a 300 m buffer encompassing suburban areas that were more vegetated and had fewer highly anthropised habitats which provide breeding sites and blood meals. In New Orleans, no association was found with vegetation cover, likely because temperature was the main driver of *Ae. albopictus* distribution (de Jesús Crespo & Rogers, 2021). *Culex*-pathogen systems were studied under temperate, continental, subtropical, and Mediterranean climates, with no clear differences across regions, suggesting that UGIs effects may not vary substantially with latitude. Studies on *Anopheles*-pathogen systems, carried out in West (Matthys et al., 2006, 2010; Yadouléton et al., 2010) and East Africa (Dongus et al., 2009; Georganos et al., 2020; Kabaria et al., 2016), showed similar results between urban agriculture and malaria risk. Finally, *Ixodes* were mainly studied in Eastern Europe and Eastern USA (similar latitudes), with a positive association with wooded habitats. A study in Malaysia reported results that were difficult to compare with those from Eastern Europe and East USA due to the differences in landscape (Kwak et al., 2021).

In countries where multiple vectors co-occur, associations with UGIs often diverge across species. In Argentina, *Ae. aegypti* was negatively associated with UGIs (Andreo et al., 2021; Estallo et al., 2018), while sandflies were linked to low NDVI zones (Berrozpe et al., 2017). In Brazil, *Ae. aegypti* was negatively associated with UGIs (Arduino et al., 2020; Heinisch et al., 2019; Honório et al., 2009; Lorenz et al., 2020; Lourenço-de-Oliveira et al., 2004; Pedrosa et al., 2020), whereas *Ae. albopictus* was positively associated (Arduino et al., 2020; Heinisch et al., 2019; Hendy et al., 2020; Honório et al., 2009; Pedrosa et al., 2020), except in deep forest (Lourenço-de-Oliveira et al., 2004). *Haemagogus* spp. was mainly associated with forested habitats (Abreu et al., 2022). In the USA, *Ae. albopictus* and *I. scapularis* were positively associated with tree cover and urban forests (Gregory et al., 2022; Hassett et al., 2022; Schwarz et al., 2020; VanAcker et al., 2019), while *Ae. aegypti* and *Culex* spp. were linked to residential areas with moderate vegetation (Brown et al., 2008; Cox et al., 2007). In Italy, both *Ae. albopictus* and *I. ricinus* were consistently associated with wooded areas (Cianci et al., 2015; Di Luca et al., 2013). In Singapore, positive correlations were observed between UGIs and *Ae. albopictus* as well as ticks, while *Ae. aegypti* showed a negative association with UGIs (Kwak et al., 2021; Sun et al., 2021). Taken together, these findings highlight distinct environments of concern, residential areas with low to moderate vegetation and forested UGIs, helping to define priority zones for integrated risk assessment, which can be further developed within a One Health framework. The impact of UGIs on pathogens can vary depending on the species of vector and host involved. For *Anopheles*-pathogen and *Ixodes*-pathogen systems, the effects of UGIs on pathogen transmission were mainly linked to the presence of the vector, but this can be more complex for other vector-pathogen systems. For DENV, the effects of UGIs depend on the system involved and differ depending on whether the vector was *Ae. aegypti* or *Ae. albopictus*. As shown in Table 3, the risk of WNV seems higher in residential areas with tree cover, as the abundance of vectors and hosts was higher in these areas. Future research in these priority areas should adopt standardized, locally adapted protocols to enhance vector monitoring, strengthen ecological understanding, and better assess how urban planning influences VBD risk.

### Data acquisition

Developing standard protocols for longitudinal monitoring of disease vector populations, pathogen dynamics, and urban vegetation features is essential. These protocols would enable local in-depth assessment of the risk of exposure to VBDs in green cities, and support comparative studies across different UGIs, study sites, and eco-socio-epidemiological contexts. On one hand, UGI indicators should be tailored to the vector studied, as relevant metrics differ. For instance, proximity to urban vegetation may be more relevant for mosquitoes, while connectivity between urban green spaces may be more appropriate for ticks. On the other hand, the effects of some UVD indicators, such as site size or connectivity, also deserve further investigation, particularly for mosquitoes. For example, Medeiros-Sousa et al. (2017) highlighted that the geographical isolation of urban green spaces can influence mosquito community composition. To evaluate VBD risk linked to UGIs, standardized metadata sets should be developed for studies and case reports, ensuring unbiased data assessment and interpretation, while also facilitating data sharing in line with FAIR (Findability, Accessibility, Interoperability and Reuse) principles (Wilkinson et al., 2016).

First, the studies included in this review encompassed considerable variability in sampling schemes and arthropod trapping methods, even among studies focusing on the same vector species. These inconsistencies can limit the representativeness of findings due to the limited number of spatial or temporal replicates (Amore et al., 2010; Carbó-Ramírez et al., 2017; de Jesús Crespo & Rogers, 2021; Dongus et al., 2009; Francisco et al., 2021; Georganos et al., 2020; Ha et al., 2021; Huynh & Minakawa, 2022; Matthys et al., 2006; Sallam et al., 2017; Wilke et al., 2017). The diversity of indicators used to assess VBD risk further complicates comparisons between studies on diseases and pathogens. For example, some studies relied on indicators based on vector abundance. Yet these metrics are known to vary significantly depending on the physiological stage of the vector sampled (eggs, larvae, adults) and are often uncorrelated with one another or with disease transmission risk (Corriveau et al., 2003). For example, a recent study in Leiden (the Netherlands) demonstrated that distribution of *Cx. pipiens* life stages differed across urban green and grey spaces where ovipositioning rates (number of egg rafts) and larval life stages were far more abundant in residential areas compared to adults being more abundant in parks (Krol et al., 2024). Other studies relied on indicators such as disease incidence and prevalence in humans, or vector infection rates, each representing distinct temporal outcomes of pathogen transmission, therefore challenging their comparison. Human health data are often aggregated spatially (Freitas et al., 2021) or tied to residential locations of patients (Muhar et al., 2000; Troyo et al., 2009), which may not correspond to sites where the patient acquired infection through the bite of a vector (Lilienfeld, 1983). This mismatch can distort within-city fine-scale risk assessments. Moreover, the approach can be limited due to a low number of reported cases (Talbot et al., 2019) or underestimation of cases (Zellweger et al., 2017). In addition to human and vector-based indicators, the detection and monitoring of pathogens in animal remain scarce despite their importance for comprehensive VBD risk assessment (ECDC, 2023). Technological advances, including smart mosquito traps (ECDC, 2023), non-invasive eDNA surveys (L’Ambert et al., 2023) and metabarcoding (Bohmann et al., 2022.), offer new opportunities for monitoring of vectors, pathogens, and host diversity. When combined with ecological network analysis, these approaches can enhance our understanding of vector-pathogen-host interactions and support integrated, harmonized assessments of VBD risks in UGIs.

Second, there is a need for precise, consistent characterization of urban green spaces, and associated vegetation. The diversity of definitions and descriptions used to characterize urban landscapes, particularly UGIs, made it difficult to generalize findings across studies (Cianci et al., 2015; Talbot et al., 2019). To address this, controlled vocabularies and ontologies are needed to standardize both field and remote data acquisition. The typology proposed by L. Jones et al. (2022) provided a clear and comprehensive framework for classifying UGIs. Further development of this typology could significantly enhance the comparability of UGI studies. The choice of spatial scale can substantially affect results, as the same variables may yield different outcomes depending on the buffer size, with even small differences of a few meters potentially influencing the findings (Landau & van Leeuwen, 2012; Manica et al., 2016). Longitudinal studies (Mackenstedt et al., 2015; Tarsitano, 2006), for instance, are complicated by the continuous evolution of urban landscape design and use over time. Several studies (Brown et al., 2008; Chen et al., 2020; Cianci et al., 2015; Gardner et al., 2013; Landau & van Leeuwen, 2012; Sun et al., 2021; Yang et al., 2019) have shown that the effects of landscape factors on VBD risks depend on the UGIs indicators used, the year, and the city investigated. First, regarding UGIs indicators, in the *Culex*-pathogen system, NDVI alone may show no association or even a negative association with VBD risk. However, NDVI cannot distinguish between deciduous and coniferous vegetation, which can have opposite effects depending on the vector system (Lockaby et al., 2016). Second, the same UGI indicator, such as percentage of vegetation cover, can produce different effects across years. For example, in New Caledonia, vegetation cover had a harmful effect on VBD risk in 2008 but showed no effect in 2012 (Zellweger et al., 2017). Finally, the city itself also plays an important role: in China, Chen et al. (2020) demonstrated that the effect of UGIs on VBD risk can differ between two cities.

The characterization of UGIs, like some landscape indices and summary statistics used in the studies involved in this review, may not adequately capture the complexity of data needed for a thorough investigation. Notably, biotic interactions significantly influence vector arthropod fitness and the dynamics of pathogen transmission within ecosystems. For example, vector trophic behavior can shift with the seasons (Hedeen et al., 2016; Levine et al., 2017), and host-switching or niche-switching throughout the seasons can lead to complex epidemiological outcomes (Lefèvre et al., 2013; Lejal et al., 2021). Indeed, vegetation may impact mosquito population dynamics in multiple ways. It may directly serve as resting site or sugar source for adults (Peach et al., 2019). Indirectly, vegetation can alter abiotic and biotic factors by modifying microclimates and by providing habitats for various animal species, including pathogen hosts, mosquito competitors, and predators (Benitez et al., 2020; Cianci et al., 2015; Myer & Johnston, 2019). As detailed below, integrating the full range of interactions mediated by arthropod disease vectors into an ecological network (food web) approach can help identify relevant biotic and abiotic factors to help design composite indicators for VBD risk assessment in UGIs.

Finally, the multifaceted nature of VBD transmission dynamics requires consideration of additional drivers beyond biological and physical parameters associated with vector and pathogen prevalence. Assessing VBD risk also requires evaluating exposure and vulnerability of populations to infected vectors. Exposure depends on multiple factors, including time spent outdoors and the use of individual protective measures such as repellents or clothing. These factors are often linked to socio-economic conditions. Accurately quantifying human and animal exposure to arthropod bites, and thus actual host-vector contact in UGIs, remains challenging due to limited data on human activities and vector biting behavior. Targeted Knowledge-Attitude-Practices (KAP) surveys are therefore encouraged to identify gaps in knowledge and prioritize protective behaviors.

Additionally, citizen science approaches can be leveraged to acquire data on human–vector contact, monitor vector occurrence, and raise public awareness about VBDs. Mobile applications such as Mosquito Alert (Južnič-Zonta et al., 2022), Muggenradar (Kampen et al., 2015), iMoustique mobile application (Gautreau, 2024) or ‘Signalement TIQUE‘ (Hassler et al., 2024) are promising tools to improve completeness and reliability of data acquisition. Earlier work has shown that such data are extremely valuable for modelling, vector management, and communication with the general public (Abourashed et al., 2021; Južnič-Zonta et al., 2022; Kampen et al., 2015).

### Theoretical frameworks and models

Most studies included in this review assessed the effects of UGIs on VBD dynamics (i.e. on vector and/or pathogen presence or abundance), often in combination with weather variables and other land cover features. However, these analyses rarely accounted for key socio-demographic confounding factors, such as human density (Bertolotti et al., 2008; Sallam et al., 2017; Vergara Cid et al., 2013), income levels (Chen et al., 2020; LaBeaud et al., 2008; Matthys et al., 2010; Zellweger et al., 2017), or urbanization features like building age (Sun et al., 2021). To address these limitations, various theoretical, statistical, and/or process-driven frameworks and models are being developed to explore the relationships between landscape features (mainly abiotic parameters) and population dynamics of arthropod vectors or VBD epidemiology. These models range from global-scale analyses (Eberhard et al., 2020) to regional-scale studies (Ducheyne et al., 2018). At a more local scale, deterministic weather-driven population dynamic models based on generic frameworks (Cailly et al., 2012) are particularly valuable. These models can map the environmental carrying capacity and identify hotspots for arthropod vector abundance, such as *Ae. albopictus,* in tropical and temperate environments (Marti et al., 2022; Tran et al., 2020). When tailored to urban green spaces, these process-based models can be generalized further by incorporating additional variables, such as local weather parameters of importance, landscape features and vector data. Mechanistic models have also been developed for other arthropod vectors, including sand flies (Erguler et al., 2019) and *Ixodes* ticks, to capture their seasonality and the transmission dynamics of *Borrelia burgdorferi* (Li et al., 2016).

Furthermore, hematophagous arthropod vectors are part of complex ecological interaction networks. These organisms contribute to ecosystem functioning and stability in multiple ways, with food webs top-down and bottom-up effects (Bellekom et al., 2021; Macfadyen et al., 2009; Ostfeld et al., 2018). Arthropod vectors may serve as prey for insectivore species at higher trophic levels, act as competitors with other species sharing similar habitats and resources, and have a major top-down influence as vectors of pathogens to a wide range of hosts. In this context, developing multilayer ecological networks (Pocock et al., 2012) appears promising for shedding light on complex and/or unexpected ecological interactions with direct relevance to vector populations and disease dynamics (Bellekom et al., 2021; Ostfeld et al., 2018; Renault et al., 2024).

To forecast the impact of urban green planning on VBD risks (Wilke et al., 2023), predictive modelling and machine-learning tools could be applied, integrating interactions between organisms and their environment. This approach requires robust and well-validated field data, including entomological information, pathogen circulation in vectors, animals, or humans, as well as data on microclimates, landscapes, and weather. In addition, complementary information on human knowledge and practices related to VBD risk reduction, socio-demographic characteristics, and the specific ecology of each vector species is essential to enhance both the accuracy and the relevance of these predictive approaches. Using these data as inputs, models can be combined with future urban (re)naturation scenarios and relevant Shared Socio-economic Pathways (SSPs), which project socio-economic, demographic, and climatic changes over the 21st century (Van Vuuren et al., 2017). By integrating these projections, models can simulate entomological and epidemiological outcomes under different plausible futures, illustrating how various urban greening strategies could influence vector populations and VBD risks over time. This approach explicitly links the predictive power of machine-learning methods with scenario-based urban planning for vegetation.

### Stakeholder engagement and implications for urban planning

Stakeholder engagement is vital for the development of effective public policies. City planners should adopt integrated, participatory approaches (Gulachenski et al., 2016; Landau & van Leeuwen, 2012; Millins et al., 2017; Sprong et al., 2018; Syal, 2021) and embrace a One Health strategy, which considers the interconnected health of humans, animals, and the environment. Implementing this approach ensures that urban planning decisions are guided by evidence on human, animal, and ecosystem health, ultimately supporting more effective and sustainable public policies. In practice, it can be operationalized across social, ecological, economic, and technical dimensions (Ellwanger et al., 2022; Koren & Butler, 2006). For example, evaluating a new urban green space involves monitoring vector populations (e.g., ticks, mosquitoes) and pathogen circulation in vectors and local wildlife, assessing environmental factors such as microclimate, and implementing management measures like targeted vegetation maintenance or mosquito breeding site control. Engaging citizens through awareness and participation is also crucial for reducing exposure and supporting effective risk mitigation.

Such an integrated approach requires robust communication and education campaigns targeting many stakeholders, including citizens, health professionals, urban planners, technicians, entomologists, biologists, architects, botanists, urban ecologists, and sociologists (Ellwanger et al., 2022). These efforts are important for fostering collaboration and ensuring comprehensive VBD risk assessments (Wilke et al., 2023). Knowledge gaps and public health priorities vary worldwide, highlighting the need for tailored strategies. Our scoping review found that 70 % of studies on ticks, particularly *Ixodes*-pathogen systems, were conducted in Northern and Eastern Europe. No studies addressed *Culex*-virus dynamics, despite increasing concern about UGIs and *Culex*-pathogen systems interactions (Krol et al., 2024). In Southern Europe, only two studies focused on *Ae. albopictus* and the arboviruses it transmits, despite rising indigenous cases (Cattaneo et al., 2025). Additionally, no research was identified on sandflies in Spain, even though recent leishmaniasis outbreaks in Madrid underscore the need to understand sandfly ecology in UGIs (Arce et al., 2013). Across all regions, urban park managers play a frontline role in VBD risk assessment and mitigation. In the USA, the primary concerns include West Nile fever and Lyme disease. In Africa and in South America, basic research is being conducted on mosquito-pathogen systems, but there is a critical lack of studies on ticks and tick-borne disease systems in urban ecosystems despite emerging concerns such as the spread of *Amblyomma cajennense* and transmission of *Rickettsia rickettsii* in urban green spaces in Rio de Janeiro (Szabó et al., 2013).

International knowledge exchange is essential, particularly between countries with long-term experience of vector presence and countries that have been recently affected. For example, *Ae. albopictus* is well-established in Italy but is now colonizing Belgium (Deblauwe et al., 2022). Similarly, countries actively advancing urban greening efforts may share insights with those that are just beginning. Most studies on urban greening have been conducted in Europe or America, with relatively few in Africa and Asia. However, a recent study in Gabon investigated the influence of urban green spaces on VBDs, focusing specifically on urban forest ecosystems (Obame-Nkoghe et al., 2023). This highlights the growing interest in these issues in different regions and the need for international cooperation.

In 2019, the Food and Agricultural Organization (FAO) published a report advocating for the global expansion of urban agriculture to address future food shortages (Taguchi & Santini, 2019). The report promoted the development of urban farms and agriculture, often associated with ecopastoralism (urban grazing) involving livestocks. Such developments may contribute to the introduction of competent hosts such as cattle for pathogens like the Crimean-Congo Hemorrhagic Fever Virus (CCHFV), transmitted by the invasive tick *Hyalomma marginatum* (Vial et al., 2016). While urban agriculture improves food access and supports ecological urban green space management, it also presents city planners with the challenge of controlling vector populations in these rapidly changing environments. Urban and peri-urban agriculture presents a significant issue in these rapidly changing environments. Urban and peri-urban agriculture represents a significant issue in Africa, South America, and Asia. Follmann et al. (2021) conducted a systematic review on the adaptation of agriculture to urbanization and proposed a conceptual workflow highlighting how urban agriculture may influence socio-economic dynamics. However, their workflow does not integrate the need to address VBDs, such as malaria, linked to urban agriculture.

Although the impact of UGIs on plant health was out of the scope of this review, the risk of introducing agricultural pests, such as the tree moth (Ciceoi et al., 2017), into cities and contaminating peri-urban forests and/or surrounding crops cannot be ignored and should be carefully assessed when introducing greenery into urban area. This is particularly important when introducing exotic plant specimens that can serve as hosts and reservoirs for devastating pathogens or their vectors (Atkins et al., 2021).

Lastly, city planners should consider the impact of global warming in their risk assessments. We did not identify articles that explicitly consider global warming as a contributing factor to VBD risks associated with urban green spaces. However, climate warming could significantly influence the spatial and temporal distribution of vectors. For instance, the tick *Hyalomma marginatum* is expanding its range into southern France (Bah et al., 2022). Similarly, for *Ae. albopictus*, future projections identified potential hotspots in the southern United Kingdom, Germany, and Benelux, while southern Spain and Portugal are predicted to become less suitable due to drought and hot summers (Caminade et al., 2012).

## Conclusion

In the context of climate change and rapid urbanization, re-naturing cities is essential for sustainability and human well-being. However, this process can also heighten the risk of VBDs that affect both humans and animals, underscoring the need to integrate health considerations into urban policy. This scoping review summarizes evidence on how UGIs influence VBD risk across various geographical areas. The effects vary by vector-pathogen system. *Aedes aegypti* was generally negatively associated with UGIs, while *Ae. albopictus* showed positive associations. For *Culex* mosquitoes and WNV transmission, both mosquito abundance and host seroprevalence tended to be higher in residential areas with moderate vegetation cover. *Anopheles* mosquitoes, linked to malaria, were positively associated with urban agriculture. Finally ticks, particularly *Ixodes ricinus*, were mainly positively associated with UGIs that include wooded areas. Overall, our review highlights the complex role of UGIs in shaping VBD risks and identifies specific environments of concern. Addressing these challenges requires collaboration among scientists, policymakers, urban planners, and citizens to ensure that urban development enhances health and sustainability.

## Acknowledgements

We would like to thank members of the VECT-OH consortium including Eline Ampt, Roser Fisa Saladrigas, Rosina Girones Llop, Gregory Lanzaro, Sonja Marie Neumeister and Xavier Roca Geronès for useful discussions during the review process.

## Authors’ contributions

FS was the principal investigator of the VECTOH Project and coordinated the manuscript draft with FF. MM and CB conducted the literature review, analyzed the articles and drafted the original version of the manuscript. NM conceptualized the data extraction and description, including the interactive map with references. All co-authors contributed to revision of the manuscript, and approved the final version of the manuscript for publication. All authors read and approved the final manuscript

## Data, scripts, code, and supplementary information availability

Supplementary information and data are available online: https://doi.org/10.5281/zenodo.17136148, (Bartholomée & Mercat, 2025).

The interactive map of studies is available online: https://doi.org/10.5281/zenodo.17136279, (Bartholomée & Mercat, 2025b).

