## Supplementary material for "Green cities and the risk for vector-borne disease transmission for humans and animals: a scoping review": Caption Additional file

This file contains all the supplementary **information** related to the article
*"Green cities and the risk of vector-borne disease transmission for humans and animals: a scoping review."*

Below is a description of each additional file and its significance:

- **Additional File 1:** Text S1 – PRISMA Extension for Scoping Review Checklist (*Additional_file_1_Text_S1_PRISMA-ScR-Checklist.doc*)
- **Additional File 2:** Text S2 – Search Strategy (*Additional_file_2_Text_S2_Search_Strategy.doc*)
- **Additional File 3:** Dataset S1 – Raw data extracted from the included articles. Each row represents a single relationship: one risk variable × one UGI variable. The dataset consists of two files: a "README" file and a data file. (*Additional_file_3_Dataset_S1_Summary_raw_data.xls*)
- **Additional File 4:** Table S1 – Overview of included articles, including key metadata, methods, and results (Additional_file_4_Table_S1_Overview.doc)
- **Additional File 5:** Table S2 – Impact of Urban Green Infrastructure on mosquito populations (*Additional_file_5_Table_S2_UGIs_Mosquito_Pop.doc*)
- **Additional File 6:** Table S3 – Impact of Urban Green Infrastructure on mosquito-borne pathogens (*Additional_file_6_Table_S3_UGIs_Mosquito_Patho.doc*)
- **Additional File 7:** Table S4 – Impact of Urban Green Infrastructure on tick populations (*Additional_file_7_Table_S4_UGIs_Tick_Pop.doc*)
- **Additional File 8:** Table S5 – Impact of Urban Green Infrastructure on tick-borne pathogens (*Additional_file_8_Table_S5_UGIs_Tick_Patho.doc*)
