## Additional 1 Text S1 for "Green cities and the risk for vector-borne disease transmission for humans and animals: a scoping review"

**Preferred Reporting Items for Systematic reviews and Meta-Analyses extension for Scoping Reviews (PRISMA-ScR) Checklist**

| **SECTION** | **ITEM** | **PRISMA-ScR CHECKLIST ITEM** | **REPORTED ON PAGE #** |
| --- | --- | --- | --- |
| **TITLE** | | | |
| Title | 1 | Identify the report as a scoping review. | #1, L1-3 |
| **ABSTRACT** | | | |
| Structured summary | 2 | Provide a structured summary that includes (as applicable): background, objectives, eligibility criteria, sources of evidence, charting methods, results, and conclusions that relate to the review questions and objectives. | #1-2, L35-62 |
| **INTRODUCTION** | | | |
| Rationale | 3 | Describe the rationale for the review in the context of what is already known. Explain why the review questions/objectives lend themselves to a scoping review approach. | #3-4, L68-125 |
| Objectives | 4 | Provide an explicit statement of the questions and objectives being addressed with reference to their key elements (e.g., population or participants, concepts, and context) or other relevant key elements used to conceptualize the review questions and/or objectives. | #4, L126-139 |
| **METHODS** | | | |
| Protocol and registration | 5 | Indicate whether a review protocol exists; state if and where it can be accessed (e.g., a Web address); and if available, provide registration information, including the registration number. | No protocol registration |
| Eligibility criteria | 6 | Specify characteristics of the sources of evidence used as eligibility criteria (e.g., years considered, language, and publication status), and provide a rationale. | #6, L245-257 |
| Information sources* | 7 | Describe all information sources in the search (e.g., databases with dates of coverage and contact with authors to identify additional sources), as well as the date the most recent search was executed. | #6, L245-257 |
| Search | 8 | Present the full electronic search strategy for at least 1 database, including any limits used, such that it could be repeated. | Additional_file_2_Text_S2 |
| Selection of sources of evidence† | 9 | State the process for selecting sources of evidence (i.e., screening and eligibility) included in the scoping review. | #6-7, L259-278 |
| Data charting process‡ | 10 | Describe the methods of charting data from the included sources of evidence (e.g., calibrated forms or forms that have been tested by the team before their use, and whether data charting was done independently or in duplicate) and any processes for obtaining and confirming data from investigators. | #7, L295-309 |
| Data items | 11 | List and define all variables for which data were sought and any assumptions and simplifications made. | #7, 280-294 |
| Critical appraisal of individual sources of evidence§ | 12 | If done, provide a rationale for conducting a critical appraisal of included sources of evidence; describe the methods used and how this information was used in any data synthesis (if appropriate). | NA |
| Synthesis of results | 13 | Describe the methods of handling and summarizing the data that were charted. | #7, L295-309 |
| **RESULTS** | | | |
| Selection of sources of evidence | 14 | Give numbers of sources of evidence screened, assessed for eligibility, and included in the review, with reasons for exclusions at each stage, ideally using a flow diagram. | #8, L320-335  Figure 1 (#9) |
| Characteristics of sources of evidence | 15 | For each source of evidence, present characteristics for which data were charted and provide the citations. | #11-13, L350-377  Figures 2 (#10)  Table 1 (#11-12)  Additional_file_4_Table_S1 (#8, L324-326) |
| Critical appraisal within sources of evidence | 16 | If done, present data on critical appraisal of included sources of evidence (see item 12). | NA |
| Results of individual sources of evidence | 17 | For each included source of evidence, present the relevant data that were charted that relate to the review questions and objectives. | #12-28, L378-890  Additional_file_4_Table_S1 (#8, L324-326)  Figures 3(#14)– 4(#18)– 5(#24)  Table 2(#17)-3(#21)-4(#26) |
| Synthesis of results | 18 | Summarize and/or present the charting results as they relate to the review questions and objectives. | #16, L534-540  #20, L645-656  #22, L697-707  #25, L820-829  #27, L869-875  Figures 3(#14)– 4(#18)– 5(#24)  Table 2(#17)-3(#21)-4(#26) |
| **DISCUSSION** | | | |
| Summary of evidence | 19 | Summarize the main results (including an overview of concepts, themes, and types of evidence available), link to the review questions and objectives, and consider the relevance to key groups. | #29, L914-963 |
| Limitations | 20 | Discuss the limitations of the scoping review process. | #28, L901-906  #29, L979-986  #30, L993-1002  #30, L1008-1030  #30-31, L1031-1044  #31, 1062-1080 |
| Conclusions | 21 | Provide a general interpretation of the results with respect to the review questions and objectives, as well as potential implications and/or next steps. | #33, L1174-1187 |
| **FUNDING** | | | |
| Funding | 22 | Describe sources of funding for the included sources of evidence, as well as sources of funding for the scoping review. Describe the role of the funders of the scoping review. | #34, L1193-1195 |

JBI = Joanna Briggs Institute; PRISMA-ScR = Preferred Reporting Items for Systematic reviews and Meta-Analyses extension for Scoping Reviews.

* Where *sources of evidence* (see second footnote) are compiled from, such as bibliographic databases, social media platforms, and Web sites.

† A more inclusive/heterogeneous term used to account for the different types of evidence or data sources (e.g., quantitative and/or qualitative research, expert opinion, and policy documents) that may be eligible in a scoping review as opposed to only studies. This is not to be confused with *information sources* (see first footnote).

‡ The frameworks by Arksey and O’Malley (6) and Levac and colleagues (7) and the JBI guidance (4, 5) refer to the process of data extraction in a scoping review as data charting*.*

§ The process of systematically examining research evidence to assess its validity, results, and relevance before using it to inform a decision. This term is used for items 12 and 19 instead of "risk of bias" (which is more applicable to systematic reviews of interventions) to include and acknowledge the various sources of evidence that may be used in a scoping review (e.g., quantitative and/or qualitative research, expert opinion, and policy document).

*From:* Tricco AC, Lillie E, Zarin W, O'Brien KK, Colquhoun H, Levac D, et al. PRISMA Extension for Scoping Reviews (PRISMAScR): Checklist and Explanation. Ann Intern Med. 2018;169:467–473. [doi: 10.7326/M18-0850](http://annals.org/aim/fullarticle/2700389/prisma-extension-scoping-reviews-prisma-scr-checklist-explanation).
