## Additional 2 Text S2 for "Green cities and the risk for vector-borne disease transmission for humans and animals: a scoping review"

**S1 Text. Search Strategy**

Key words used for the search in the three databases: Scopus, PubMed and Web of Science.

When available, we use proximity operator to find papers where the terms joined by the operator are within a specified of words of each other (w/ for Scopus, and NEAR in Web of Science):

- **Urban:** urban* OR cities OR city OR built-up OR roof
- **Greening:** green OR biodiversity OR forest OR rewilding OR park OR nature OR veget* OR corridor OR farm OR agriculture OR garden OR plant OR tree
- **Vectors:** vector OR arthropod OR insect OR mosquito OR tick
- **Vector-borne disease:** vector-borne disease OR tick-borne disease OR virus OR bacteria OR parasite OR pathogen

PubMed: last access on October 6, 2022

Search equation:

*(urban*[Title/Abstract] OR cities[Title/Abstract] OR city[Title/Abstract] OR built-up[Title/Abstract] OR roof*[Title/Abstract]) AND (green*[Title/Abstract] OR biodiversity[Title/Abstract] OR forest*[Title/Abstract] OR rewilding[Title/Abstract] OR park*[Title/Abstract] OR nature*[Title/Abstract] OR veget*[Title/Abstract] OR corridor*[Title/Abstract] OR farm*[Title/Abstract] OR agriculture[Title/Abstract] OR garden*[Title/Abstract] OR plant*[Title/Abstract] OR tree*[Title/Abstract]) AND (vector*[Title/Abstract] OR arthropod*[Title/Abstract] OR insect*[Title/Abstract] OR mosquito*[Title/Abstract] OR tick*[Title/Abstract]) AND (vector borne disease*[Title/Abstract] OR tick borne disease*[Title/Abstract] OR virus[Title/Abstract] OR bacteria[Title/Abstract] OR parasite*[Title/Abstract] OR pathogen*[Title/Abstract])*

Scopus: last access on October 6, 2022

Search equation:

*( TITLE-ABS-KEY ( urban* W/15 ( green* OR biodiversity OR forest* OR rewilding OR park* OR nature* OR veget* OR corridor* OR farm* OR agriculture OR garden* OR plant* OR tree* ) OR city W/15 ( green* OR biodiversity OR forest* OR rewilding OR park* OR nature* OR veget* OR corridor* OR farm* OR agriculture OR garden* OR plant* OR tree* ) OR cities W/15 ( green* OR biodiversity OR forest* OR rewilding OR park* OR nature* OR veget* OR corridor* OR farm* OR agriculture OR garden* OR plant* OR tree* ) OR built-up W/15 ( green* OR biodiversity OR forest* OR rewilding OR park* OR nature* OR veget* OR corridor* OR farm* OR agriculture OR garden* OR plant* OR tree* ) ) AND TITLE-ABS-KEY ( vector* OR arthropod* OR insect* OR mosquito* OR tick* ) AND TITLE-ABS-KEY ( ( vector AND borne AND disease ) OR ( tick AND borne AND disease ) OR virus OR bacteria OR parasite OR pathogen ) )*

Web of Science: last access on October 6, 2022

Search equation:

*urban* NEAR/15 (green* OR biodiversity OR forest* OR rewilding OR park* OR nature* OR veget* OR corridor* OR farm* OR agriculture OR garden* OR plant* OR tree*) OR cities NEAR/15 (green* OR biodiversity OR forest* OR rewilding OR park* OR nature* OR veget* OR corridor* OR farm* OR agriculture OR garden* OR plant* OR tree*) OR city NEAR/15 (green* OR biodiversity OR forest* OR rewilding OR park* OR nature* OR veget* OR corridor* OR farm* OR agriculture OR garden* OR plant* OR tree*) OR built-up NEAR/15 (green* OR biodiversity OR forest* OR rewilding OR park* OR nature* OR veget* OR corridor* OR farm* OR agriculture OR garden* OR plant* OR tree*) OR roof* NEAR/15 (green* OR biodiversity OR forest* OR rewilding OR park* OR nature* OR veget* OR corridor* OR farm* OR agriculture OR garden* OR plant* OR tree*) (Topic) AND arthropod* OR mosquito* OR tick* OR vector* OR insect (Topic) AND vector borne disease OR tick borne disease OR virus OR bacteria OR parasite OR pathogen (Topic)*
