## Additional 4 Table S1 for "Green cities and the risk for vector-borne disease transmission for humans and animals: a scoping review"

| **References** | **Country (city)** | **Vector** | **Species/Genus** | **Pathogen** | **Host** | **Studied stage** | **Risk indicator** | **Vegetation indicator** | **Main results** |
| --- | --- | --- | --- | --- | --- | --- | --- | --- | --- |
| **MOSQUITOES** | | | | | | | | | |
| (Cox et al. 2007) | USA, Puerto Rico (San Juan) | Mosquitoes | *Aedes aegypti, Aedes mediovittatus, Culex quinquefasciatus* | NA | NA | Immature stages (larvae) | Abundance of larvae | Landcover classes: high-density housing (HDH) (urban region with few forests patches), low-density housing (LDH) (more forests patches) and forests (F). | The abundance of *Aedes aegypti* is higher in low and high-density housing than in urban forests. The abundance of *Aedes mediovittatus* is higher in low-density than in high-density housing. *Culex quinquefasciatus* is less abundant in urban forests than in low-density housing. *Culex antillummagnorum* is more abundant in urban forests and in low-density residential areas than in high-residential areas. |
| (Wong et al. 2018) | China (Hong-Kong) | Mosquitoes | *Aedes aegypti, Aedes albopictusa, Aedes japonicus, Anopheles spp, Culex quinquefasciatus, Culex tritaeniorhynchus* | NA | NA | Adults | Mosquito abundance | 3 categories:   - Green Roofs  - Bare roofs   - Low-elevation gardens | Male abundance was consistently lower on the rooftop sites compared to female abundance. The abundance of males was lower on green and bare roofs compared to low-elevation gardens. For *Aedes albopictu*s and *Aedes japonicus*, there was no difference between the green roof and the bare roof. For *Culex quinquefasciatus* males, the abundance was greater on bare roofs than on green roofs. |
| (Vergara Cid et al. 2013) | Argentina (Cordoba) | Mosquitoes | *Culex quinquefasciatus complex ** | Saint-Louis Encephalitis Virus (NA) | Human | NA | Saint-Louis Encephalitis (SLE) Human prevalence | Landcover: high and low density urban construction + Distance to areas with NDVI>0.3 | The main landscape elements contributing to the outbreak of SLE in 2010 were the Distance to places with NDVI > 0.3 and the presence of low-density urban constructions, like residential areas. |
| (Troyo et al. 2009) | Costa Rica (Puntarenas) | Mosquitoes | *Aedes aegypti** | DENV* | Human | NA | Dengue Human cases | Weekly enhanced vegetation index (EVI) + mean NDVI + land use: tree (tree cover), grass | Mean NDVI was negatively correlated with dengue incidence in the dry season. The percentage of tree cover didn't show association with dengue incidence. The weekly EVI seemed to increase dengue incidence. |
| (Brown et al. 2008) | USA (New Haven) | Mosquitoes | *Culex pipiens, Culex restuans, Culex salinarius, Aedes vexans* | NA | NA | Adults | Mosquito abundance | NDVI, NDWI, moderate and highly vegetated residential areas | The abundance of mosquitoes is positively associated with the NDVI and the NDWI. The abundance of *Culex pipiens, Culex restuans*, and *Aedes vexans* is higher in moderately vegetated residential areas than in highly vegetated residential areas. The abundance of *Culex salinarius* is higher in highly vegetated areas than in residential areas with less vegetation. |
| (Bertolotti et al. 2008) | USA (Chicago) | Mosquitoes | *Culex pipiens, Culex restuans* | WNV | Birds | Adults | WNV diversity in vector and in host | Housing density, Residential land cover, Urban land cover, Distance from natural area | The diversity of West Nile Virus (WNV) among mosquitoes captured in residential areas surpasses the viral diversity observed in mosquitoes captured in natural "urban green space" sites. Conversely, the WNV diversity among birds collected in residential areas exhibits a reverse relationship. This phenomenon in birds might be attributed to variations in avian species diversity between natural and residential sites. |
| (Liu et al. 2008) | USA (Indiana polis) | Mosquitoes | *Culex sp.* | WNV | NA | Adults | WNV mosquito infections | % urban agriculture,  size of urban wetland | The major environmental factors contributing to WNV outbreaks are the percentage of agriculture and the size of wetlands. |
| (Honório et al. 2009) | Brazil (Rio de Janeiro) | Mosquitoes | *Aedes aegypti, Aedes albopictus* | NA | NA | Immature stages (larvae) | Abundance of mosquitoes | 4 land cover classes: densely built area. Low vegetation coverage. Medium vegetation coverage. High vegetation coverage | Aedes albopictus was more abundant in areas with high to medium-dense vegetation coverage, whereas Aedes aegypti dominated the densely populated areas. However, the Distance to green urban vegetation was positively associated with the abundance of both. |
| (de Abreu et al. 2022) | Brazil (Rio de Janeiro) | Mosquitoes | *Haemagogus leucocelaenus, Haemagogus janthinomys/capricornii* | Yellow Fewer Virus (YFV) | Human and Non Human Primate | Adults | Mosquito abundance, Infection in vector (MIR) and Yellow Fewer Host cases | Observation of main landcover classes: urban fragment and urban intra-domicile. Remote Sensing: NDVI, forest fragment side. | The abundance of *Haemagogus* mosquitoes was positively associated with the NDVI, but there was no difference between urban forest, urban pasture, and densely urban areas. The *Haemagogus* pathogen infection and the yellow fever human prevalence were higher in the urban forest fragment than in the other land cover. |
| (Matthys et al. 2010) | Cote d'Ivoire (Abidjan) | Mosquitoes | *Anopheles spp., Culex spp.* | NA | NA | Immature stages (larvae and pupae) | Density of mosquitoes | Agricultural zones: habitat type, physical characteristics, presence of vegetation, occurrence of predators. | The density of *Anopheles* varied between the urban agricultural zones. The density (larvae and pupae) was higher in zones with many agricultural trenches, which often became shallow water bodies and became the most productive habitat type for pupae and larvae. It was negatively correlated with the presence of floating vegetation. |
| (Dongus et al. 2009) | Tanzania (Dar es Salaam) | Mosquitoes | *Anopheles* | NA | NA | Immature stages (larvae) | Presence of *Anopheles* larvae | Variable on agricultural characteristics (land use, ward, farming site, production type, type of crops) | Cultivated areas are more likely than others (including urban settlements) to contain breeding habitats with *Anopheles larvae*. Urban agriculture and the presence of urban irrigation wells positively influenced the breeding sites of *Anopheles*. Contrary to leguminous areas, the sugar cane and leafy vegetable areas increased the positivity of breeding sites. Crop diversity is negatively associated with the positivity of breeding sites during the wet season and negatively during the wet season. |
| (Heinisch et al. 2019) | Brazil (Sao Paulo) | Mosquitoes | *Aedes aegypti, Aedes albopictus* | NA | NA | Immature stages (eggs) | Oviposition Rate Ratio (ORR) ( ratio of the egg count to the corresponding count of a reference value) | 3 areas within 1 park: internal (more vegetation, intermediate and peripheral (little vegetation close to inhabited) | There is a spatial segregation of *Aedes aegypti*, with greater oviposition in the park peripheral area, which has lower vegetation coverage and is closer to the urban area. *Aedes albopictus* is more associated with an internal and intermediate park zone. |
| (Lorenz et al. 2020) | Brazil (Sao Paulo) | Mosquitoes | *Aedes aegypti* | NA | NA | Adults (only female) | Abundance | Seven land cover classes: pavement, tile roof, asbestos roof, roof slab, green area, exposed soil, water + shadow areas | *Aedes aegypti* abundance is negatively associated with urban green areas and positively with asbestos roofs and exposed soil. |
| (Matthys et al. 2006) | Cote d'Ivoire (Man) | Mosquitoes | *Anopheles gambia*, Anopheles funestus** | *Plasmodium falciparum, Plasmodium malariae* | Human | NA | Human malaria prevalence | Agricultural zone (type of crop and practice) | There were significant malaria infection differences between agricultural zones. The lowest prevalence was in traditional small-holder rice plots and large rice perimeters, whereas the highest was in mixed crop systems. |
| (Pedrosa et al. 2020) | Brazil (Mariana, Ouro Preto) | Mosquitoes | *Aedes aegypti, Aedes albopictus* | NA | NA | Adults and immature stages (eggs and larave) | Abundance | 3 biotopes: green areas, residential area close to green area, very urbanized and Distance to urban green areas | There were more *Aedes albopictus* in residential areas close to green areas than in green areas and very urbanized regions. There was a positive association between the abundance of *Aedes albopictus* and the distance to urban green spaces. There were more *Aedes aegypt*i in residential areas, which are more urbanized than urban green areas. |
| (Paul et al. 2018) | Bangladesh (Dhaka) | Mosquitoes | *Aedes aegypti, Aedes albopictus* | NA | NA | Immature stages (larvae and pupae) | Presence and abundance of larvae and pupae | Classification of egg-laying sites + presence of vegetation near the containers | The risk factors for *Aedes* are the presence of containers outside and vegetation close to the containers. |
| (Schwarz et al. 2020) | USA (Greensboro) | Mosquitoes | *Aedes albopictus, Aedes triseriatus, Aedes hendersoni* | NA | NA | Immature stages (eggs) | Abundance of eggs | 3 biotopes: campus, park and forest. 3 tree height levels: ground, mid-tree and canopy level. | The three species were most abundant in urban forests, followed by urban parks and campuses (densely urbanized areas). The abundance of *Aedes albopictus* followed the same pattern. There was no difference in the abundance of *Aedes triseriatus* between the urban forest and the urban park, but it is more abundant in the urban forest and the urban park than on campus. *Aedes hendersoni* is only present in urban forests. *Aedes albopictus* is more present at the canopy than at the ground level, contrary to the other species. |
| (Talbot et al. 2019) | Canada (Ottawa) | Mosquitoes | *Culex pipiens/Culex restuans, Culex salinarius, Aedes vexans,  Anopheles punctipennis, Ochlerotatus japonicus, Ochler  otatus  trivittatus, Ochlerotatus triseriatus/Ochlerotatus  hendersoni* | WNV | Human | Adults | WN human incidence and WNV mosquito infection | Land cover | The results show that natural areas may affect WNV risk in humans, but not urban parks. The proportion of urban parks was not associated with epidemiological WNV risk. WNV human incidence was positively correlated with population density and the percentage of aged housing |
| (Trawinski et al. 2010) | USA (Amherst) | Mosquitoes | *Culex pipiens-restuans, Aedes vexans* | NA | NA | Adults | Abundance | Land use, Land cover, NDVI | In all the tested buffers, the abundance of *Aedes vexans* was not associated with NDVI. The influence of NDVI on the abundance of *Culex pipiens* depends on the buffer and the sampling weeks. The correlation can be positive for the 400 m and 1000m buffer or inexistent for the 200m, 400m, and 1000m buffer. *Culex pipiens – restuans* is positively correlated with human density and housing unit density. |
| (Muhar et al. 2000) | Australia (Brisbane) | Mosquitoes | *Aedes vigilax*, Culex annulirostris** | Ross River Virus (RRV) | Human | NA | RRV Human case | 7 landcover class : littoral wetland, ephemeral wetland, open freshwater, riparian vegetation, Melaleuca open forest, wet eucalyptus open forest, other bushlands. | The higher human cases were located in the suburbs, positively correlated with bushlands and riparian forests, and wetlands. It is linked to the presence of many breeding sites and the presence of dense vegetation. |
| (Amore et al. 2010) | USA (Chicago) | Mosquitoes | *Culex pipiens* | WNV | NA | Adults | WNV diversity in vector | Residential areas and green urban spaces | The WNV genetic diversity is lower from mosquitoes captured in green urban spaces than in residential areas. This variation may be explained by fine-scale variations in avian hosts or microclimate. |
| (Ha et al. 2021) | Vietnam (Hanoi) | Mosquitoes | *Culex tritaeniorhynchus, Culex quinquefasciatus, Culex vishnui* | NA | NA | Adults (females) | Abundance of mosquitoes females | NDVI and land cover | There is a negative correlation between the forest cover ratio at the 250 m buffer and mosquito abundance and a positive correlation between rice cover at 250 m and mosquito abundance. There is no association between NDVI at the 250 m buffer and the abundance. The abundance was associated significantly with the bi-monthly mean rainfall and negatively with the human population density.hedy |
| (Che Dom et al. 2016) | Malaysia (Shah Alam) | Mosquitoes | *Aedes aegypti, Aedes albopictus* | NA | NA | Immature stages (larvae) | Positivity and productivity of breeding sites | Type of breeding sites and description of the vegetation on the water (algae, emergent aquatic grass and floatic aquatic vegetation) | Larvae prefer breeding sites with clear water and the presence of floating debris. Flower vases are positive breeding sites of *Aedes albopictus*. The proximity of the breeding sites to vegetation increases the probability of positivity. |
| (Estallo et al. 2018) | Argentina (Cordoba) | Mosquitoes | *Aedes aegypti* | NA | NA | Immature stages (larvae) | Presence of larvae | NDVI,   Distance to vegetation | The abundance of breeding sites is positively associated with the Distance to vegetation. |
| (Levine et al. 2017) | Usa (Atlanta) | Mosquitoes | *Culex spp.* | WNV | passerine birds | Adults | Sero prevalence in birds and MIR of *Culex* | 5 urban micro habitats: - mixed-use parks (divided into wooded and water sections) - residential areas - old- growth forest patches - Zoo Atlanta | There is no association between the WNV *Culex* infection and the land cover (residential areas, urban parks with water and wood, and urban forest). However, the WNV bird's seroprevalence is higher in residential areas than in urban parks with water and urban forests.  The lack of any such avian-to-mammalian feeding shift during the critical, highly infectious months of August and September in the Atlanta area contributes to diminished human transmission levels observed. The shift of avian feeding from American robins (highly competent for WNV) to northern cardinals (moderately competent for WNV) occurred between July and August, precisely before the critical infectious months of August and September in either the host or vector populations, and helps account lower rates of avian WNV infection in the urban forest patch. |
| (Zellweger et al. 2017) | New Caledonia (Numea) | Mosquitoes | *Aedes aegypti** | DENV* | Human | NA | Dengue Human cases | Vegetation cover: % of surface covered by vegetation / House and people density/ Built environment/Socio-economic satut/demographics | Dengue incidence rates were higher in neighborhoods with higher vegetation coverage in 2008-2009. To reduce garden sites may help reduce breeding sites. There was no association between dengue incidence and the vegetation coverage in 2012-2013. |
| (Talbot et al. 2021) | Colombia (Ibagué), Ecuador (Manta) and Argentina (Posadas) | Mosquitoes | *Aedes aegypti, Aedes albopictus* | NA | NA | Adults | Density of mosquitoes | Green areas and decorative vegetation | The presence of decorative vegetation and green areas next to households are factors which increase the abundance of *Aedes*. |
| (Yadouléton et al. 2010) | Benin (Cotonou, Porto-Novo, Parakou) | Mosquitoes | *Anopheles gambiae s.l.* | *Plasmodium falciparum* | Human | Adults | Human-biting-rate and Infection in mosquitoes | Households closed to vegetable farms and househomds far from vegetable farms | The human biting rates and sporozoite infective *Anopheles gambiae* were higher in households close to vegetable farms than in those on the far side of the farms because of the presence of breeding sites in farms. |
| (Freitas et al. 2021) | Brazil (Rio de Janeiro) | Mosquitoes | *Aedes aegypti** | CHIKV | Human | NA | Chikungunya human incidence | % green areas per neighborhood | When no spatial effect is considered, the proportion of green areas decreases the human incidence. When spatial effect is taken into account, there is no effect on green areas. Human incidence was mainly influenced by the inverse of the socioeconomic development index and by the temperature. |
| (LaBeaud et al. 2008) | USA (Cleveland) | Mosquitoes | *Culex sp.* | WNV | Human | Adult | WN human incidence | Landcover class: open, forest and wetland versus residential and other urban | WNV human cases were negatively associated with urban green spaces, such as urban parkland and urban agriculture. WNV human cases were positively associated with high income, population over 65 years old and road densitythe |
| (Francisco et al. 2021) | Philipinnes (Manila) | Mosquitoes | *Aedes aegypti, Aedes albopictus* | DENV* | Human | Immature stages (eggs) | Abundance of eggs (ovitrap index) and Dengue Human incidence | NDVI, Land surface temperature land use (30 categories) | There is a negative correlation between human dengue incidence and vegetations. Human dengue incidence was positively correlated with the ovitrap index and temperature at a three-month lag qand with the precipitation at a one-month lag and *Aedes* mosquito abundance is positively associated with vegetation. |
| (Sun et al. 2021) | Singapore (Singapore) | Mosquitoes | *Aedes aegypti, Aedes albopictus* | NA | NA | Adults | Abundance of mosquitoes | 4 vegetation land cover classes: water, grass, forest and managed vegetation + urban variables: drain line density, building age | The proportion of urban forests decreases the abundance of *Aedes aegypti* and has the opposite effect on the abundance of *Aedes albopictus.* The proportion of urban grass decreases the abundance of both species. Their abundance is positively associated with the percentage of urban-managed vegetation. |
| (Lockaby et al. 2016) | Usa (Atlanta) | Mosquitoes | *Culex quinquefasciatus* | WNV* | NA | Adults | Vector index: product of the abundance of Cx. quinquefasciatus and the maximum likelihood of the infection rate. | Landcover of 4 classes: water, forest, impervious and open space.  Types of forests are also described, according to tree species, height and percent of shrubs. | The Vector Index varies according to the type of forest. The vector index decreased significantly in urban pine forests compared to deciduous forests, either because of a habitat unsuitable for corvid species or an unfavorable microclimate. Larger forest patches are also negatively correlated to the vector index, which may be because of bird species diversity (Dilution effect). |
| (Gardner et al. 2013) | USA (Chicago) | Mosquitoes | *Culex restuans, Culex pipiens* | NA | NA | Immature stages (larvae) | Abundance of larvae/basin/week | Field observation: description of the vegetation in a buffer of 25 m around the catch basin (tree genus and height, shrubs type and height, presence of flowers) | Larval rates varied spatially and temporally. Some trees, like elms or trees of less than one meter in height, influenced positively the larval rates. Other, like ash, have negative impacts. Shrubs play a role as a resting place and source of nectar as well as bird blood. In addition, the waste products from the trees modify the composition of the pond, which can attract more mosquitoes and allow their survival. The presence of flowers is negatively correlated with the larval rates. |
| (Cianci et al. 2015) | Italy (Rome) | Mosquitoes | *Aedes albopictus* | NA | NA | Immature stages (eggs) | Abundance of eggs | 7 landcover classes : buildings, roads, hedge, botanical garden, grass, tree and car parks + buffer zone around of 15 m | Open areas (grass) don't attract oviposition females. A positive correlation exists between the abundance of eggs and trees and solar radiation. Favorable habitats are covered by trees, with sun. |
| (Liu et al. 2011) | USA (Whashington DC) | Mosquitoes | *Culex pipiens*, Culex restuans** | WNV | Corvids birds and humans | NA | WNV Birds and human cases | Classification of impervious surface in 5 classes and of canopy cover in 4 classes | Bird cases correlate positively with a medium canopy density (30%) and human cases with a lower canopy density. Tree canopy allows birds to forage and nest, and mosquitoes feed on these birds. |
| (Liu et al. 2012) | USA (Los Angeles) | Mosquitoes | *Culex ttigmatosoma, Culex quinquefasciatus , Culex tarsalis, Culex erythrothorax, Culex incidens, Culex thriambus* | WNV | NA | Adults | WNV mosquito infections | NDVI | The NDVI was not associated with infection in mosquitoes. There were some confounding facts that limit the NDVI interpretation: some elevated lands in LA were well covered by vegetation but with relatively lower temperatures and, thus, likely smaller mosquito populations. |
| (Landau et al. 2012) | USA (Tucson) | Mosquitoes | *Aedes aegypti, Culex quinquefasciatus* | NA | NA | Adult | Abundance of mosquitoes | NDVI + 4 land use categories (Managed Green Spaces, Mixed Use, residential and washes) + 11 land cover categories (bare earth, pavement, structure, pool, water, shadiw, herbaceous, shrub and 3 tree height classes) | *Culex quinquefasciatus* abundance has positive associations with medium-height trees and negative associations with shrub presence.  *Aedes aegypti* abundance has positive associations with medium-height trees. |
| (Araujo et al. 2015) | Brazil (Sao Paulo) | Mosquitoes | *Aedes aegypti** | DENV* | Human | NA | Dengue Human cases | Vegetation covercategories ( low, moderate and high) | The majority of dengue cases occurred in areas with low vegetation cover, and a land surface temperature > 32°C. Areas with high vegetation cover were cooler (26 ± 2 ◦C) compared to areas with moderate or low vegetation cover (both, 29 ± 2 ◦C). The microclimate of less vegetated areas is more favorable to mosquito breeding. Less dengue cases occurred in areas with low socioeconomic values and high population density. |
| (Kabaria et al. 2016) | Tanzania (Dar es Salaam) | Mosquitoes | *Anopheles gambiae*, Anopheles funestus** | *Plasmodium falciparum* | Human | NA | Plasmodium falciparum human prevalence | 13 land cover classes: urban classes (depending on the type of buildings), vegetation classes (light, dense or riverine), water classes and bare soil classes (sand, bare soils and bare soils with vegetation | Malaria positivity is higher in dense and riverine vegetation. The risk of malaria infection increases with the increase in lush vegetation. Malaria risk in the city is not homogenous. |
| (Levine et al. 2016) | Usa (Atlanta) | Mosquitoes | *Culex spp.* | WNV | passerine birds | Adults | Sero prevalence in birds and MIR of *Culex* | 4 urban microhabitats - mixed-use parks, divided into wooded and water sections; - residential areas - and old-growth forest patches | The highest total bird seroprevalence was from the residential microhabitat types (the less vegetated habitat), while the lowest was from the Forest Patch microhabitat types. There was a slightly positive relationship between diversity and avian seroprevalence, perhaps because of the amplification effect. The residential microhabitat type also had the highest total WNV MIR observed. There was a slightly negative relationship between bird diversity and mosquito infection. There was no association between mosquito infection rates and avian seroprevalence rates. |
| (Wong et al. 2016) | China (Hong-Kong) | Mosquitoes | *Aedes aegypti, Aedes albopictus, Aedes japonicus, Anopheles spp, Culex quinquefasciatus, Culex tritaeniorhynchus* | NA | NA | Adults | Mosquito diversity and abundance | 3 categories:   - Green Roofs  - Bare roofs   - Low-elevation gardens | The vector abundance was higher in bare roofs than green roofs, attributed to the occasional presence of water pools in depressions of roofing membranes after rainfall. The richness and evenness of mosquito species in vegetated urban habitats (green roofs and low-elevation gardens) were more significant than in bare urban habitats. |
| (Carbo-Ramirez et al. 2017) | Mexico (Pachuca and Xalapa), Germany (Freiburg) | Mosquitoes | NA | Avian haemosporidians: P*lasmodium, Haemoproteus* and *Leucocytozoon* | Birds | NA | Bird prevalence and pathogen diversity | 3 urban greenspaces : nearctic urban greenspace, neotropical urban green- space, palearctic urban greenspace | The three green urban spaces show ecological parallels: haemosporidian infection prevalence is similar, like the parasite abundance structure. The bird's species diversity is higher in neotropical urban green spaces and similar in the neartic and paleartic urban green spaces. The pathogen diversity is higher in the paleartic urban green spaces and similar in the two other categories. |
| (Wilke et al. 2017) | Brazil (Sao Paulo) | Mosquitoes | *Aedes albopictus, Aedes fluviatilis, Aedes scapularis, Wyeomyia, Toxorhynchites, Culex quinquefasciatus, Culex nigripalpus, Limatus, Trichoprosopon* | NA | NA | Adults and immature stages (larvae) | Mosquito abundance | 2 urban parks | *Aedes albopictus* was the most abundant mosquito, with five other vector species: *Aedes aegypti,Aedes fluviatilis, Aedes scapularis, Culex nigripalpus* and *Culex quinquefasciatus*. The correlation between the abundance of mosquitoes and rainfall depended on the parks. This relation may be related to the availability of breeding sites in parks. There is no difference of diversity between parks. |
| (Sallam et al. 2017) | USA (New Orleans) | Mosquitoes | *Culex quinquefasciatus* | WNV* | Human | Adults | Vector-host contact | Land use and land cover :  - 3 urban classes (residential, industrial and commercial, other urban built-tup) - 1 class of waterbodies and  - 4 classes of vegetation : (1) non-forested wetland, (2) forest wetland, (3) deciduous forestland, and (4) tree density | Non-forested wetland land cover type (1 type on vegetation type) alongside tree density (TD) positively correlated with the vector-host contact ratios. However, the increase in residential and urban settings shared a reduced negative influence on the spatial distribution of the WNV mosquito vector. |
| (Medeiros-Sousa, et al. 2017) | Brazil (Sao Paulo) | Mosquitoes | *Culex nigripalpus, Aedes albopictus, Culex quinquefasciatus, Aedes fluviatilis, Aedes scapularis, Culex declarator, Aedes aegypti, Culex chidesteri, Limatus durhami and Culex lygrus* | NA | NA | Adults | Mosquito diversity and richness, Mosquito abundance | 9 Urban parks : measure of green areas (hectare)   + 1km buffer landscape metrics (vegetation, water bodies, urban) + the degree of isolation of each park (proximity index) | Larger green areas tend to be more species-rich than smaller green ones. Similarity increases between assemblages in species-poor sites (nested pattern). The vector mosquitoes, *Culex nigripalpus, Aedes scapularis, Aedes fluviatilis, Culex declarator, Aedes aegypti, Aedes albopictus* and *Cules quinquefasciatus* were the most common and abundant species in the urban green spaces in species-rich and species-poor locations. |
| (Myer et al. 2019) | USA (New York) | Mosquitoes | *Culex pipiens, Culex restuans* | WNV | NA | Adults | WNV positive infection rate in mosquitoes | Landcover class: open, forest and wetland versus residential and other urban | WVN mosquito infection is negatively correlated with emergent herbaceous wetlands and green spaces, maybe because of the ecology of *Culex pipiens. Culex pipiens* can use artificial breeding sites. Mosquito infection is also negatively correlated with mean precipitation lagged at 1 week, the high intensity development and open water. It was positively correlated with the number of mosquitoes trapped and the mean temperature lagged at 2 weeks. |
| (Tian et al. 2019) | China (Zhejiang (in lab)) | Mosquitoes | *Aedes albopictus* | NA | NA | Adults (only females) | Survival and fecundity | In lab: different ornamental plants (11 species), with or without flowers | Mosquitoes' survival fed on *Ligustrum quihoui* was more significant than those provided on the other plant species or water alone. The lifespan of mosquitoes given access to N. domestica was significantly shorter than that of mosquitoes given access to other plant cuttings, compared with sucrose or only water. Mosquitoes' fecundity varies depending on plant species (females produce more eggs when exposed to *Abelia grandiflora* than other plants). The presence of decorative vegetation and green areas for households and the absence of water bodies increase the abundance of *Aedes.* |
| (Ogashawara et al. 2019) | Brazil (Sao Paulo) | Mosquitoes | *Aedes aegypti** | DENV* | Human | NA | Dengue Human cases | NDVI, NDBI, NDWI, LST | Dengue human cases were significantly correlated with sea surface temperatures and mimum surface temperature. There was no relationship with other environmental variables because of the failure to account for socioeconomic factors and the powerful influence of temperament on the vector |
| (Demets et al. 2020) | Canada (Toronto) | Mosquitoes | *Culex pipiens, Culex restuans* | NA | NA | Adults | Mosquito abundance | NDVI (Normalized Difference Vegetation Index)  NDBI (Normalized Difference Built-up Index),   gNDWI (green-based Normalized Difference Water Index,   sNDWI (SWIR-based Normalized Difference Water Index) and   NBR (Normalized Burn Ratio) | The best predictor of *Culex* abundance is sNDWI, which is a marker of the moisture content of the canopy and necessary in *Culex* ecology. |
| (Arduino et al. 2020) | Brazil (Sao Paulo) | Mosquitoes | *Aedes aegypti, Aedes albopictus* | NA | NA | Immature stages (larvae) | Mosquito presence | Landcover (%) of: urbanized, woods, grass-shrub | *Aedes albopictus* was more concentrated in environments with more vegetation, even those in urban areas. *Aedes aegypti* was more concentrated in the urban environment. They coexist in some areas. Microclimate can explain a part of the different distribution. |
| (Vieira et al. 2020) | Brazil (Sinop) | Mosquitoes | *Anophelinae, Culicinae, Aedeomyiini, Aedini* | NA | NA | Adults | Mosquito diversity and abundance | 4 categories : Highly urbanized neighborhoods (HUN), Neighborhoods located next to PPAs (NPP), Neighborhoods with forest fragments (NFF), Urban forest parks located in completely urbanized neighborhoods (PPAs) | The mosquito's abundance, diversity and richness were higher in urban forests, in neighborhoods with forest fragments, in highly urbanized neighborhoods and neighborhoods next to the forest. |
| (Lee et al. 2020) | Malaysia (Klang Valley) | Mosquitoes | *Aedes albopictus, Armigeres spp., Culex spp., Culex quinquefcasciatus, Mansonia spp., Tripteroides spp., Uranotaenia spp.* | NA | NA | Adults and immature stages (larvae) | Mosquitoes diversity, abundance of *Aedes albopictus* and of breeding sites | 3 classes: urban park, goat farm in a peri-urban area and natural forest hiking | The mosquito diversity is higher peri-urban farms than in urban parks. *Aedes albopictus* is more abundant in urban environments. |
| (Hendy et al. 2020) | Brazil (Manaus) | Mosquitoes | *Aedes aegypti, Aedes albopictus, Culex pipiens* | NA | NA | Adults | Mosquito diversity and vector abundance | Urban parks, NDVI and distance from edge | No relation is found between total mosquito abundance, the mean number of species, the NDVI, and the distance edge. *Aedes aegypti* and *Aedes albopictus* were both located within the park (to 100m inside). The tiger mosquito abundance was highest at the forest edge and was positively correlated with medium NDVI and negatively with low and high NDVI. There was a cohabitation of *Aedes albopictus* and *Aedes aegypti* within the park. |
| (Poh et al. 2020) | USA (Dallas) | Mosquitoes | *Culex quinquefasciatus* | WNV | NA | Adults (only female) | WNV mosquito infections | NDVI, % impervious surface, % urbanisation | The most significant predictors of the presence of WNV in *Culex quinquefasciatus* pools were increased urbanization (index based on greater population density, lower NDVI, higher coverage of urban land types, and more impervious surfaces), older human populations, and lower elevation. There were more *Aedes albopictus* in residential areas close to green areas than in green areas and very urbanized regions. There were more *Aedes aegypti* in residential areas and is more urbanized than in green areas. There was a positive correlation between *Aedes aegypti* abundance and the increasing distance to vegetation |
| (Benitez et al. 2020) | Argentina (Cordoba) | Mosquitoes | *Aedes aegypti* | NA | NA | Immature stages (larvae) | Mosquito abundance | Landcover: buildings, arboreal vegetation, herbaceous vegetation, soil with scanty vegetation and water | *Aedes aegypti* abundance was more abundant in areas with less arboreal vegetation, which corresponded to more urbanized areas. |
| (Georganos et al. 2020) | Tanzania (Dar es Salaam) and Uganda (Kampala) | Mosquitoes | *Anopheles gambiae*, Anopheles arabiensis*, Anopheles funestus** | *Plasmodium falciparum* | Human | NA | *Plasmodium falciparum* human prevalence | Land cover, land use and NDVI+C66B66:K66L66D66A66:K66 | In both cities, malaria prevalence is positively associated with dense informal settlements, wetlands, marshes, riparian vegetation, and urban agriculture. *Anopheles* vectors can use swamps, marshes, and riparian vegetation as breeding sites. |
| (Huynh et al. 2022) | Vietnam (Ho Chi Minh) | Mosquitoes | *Aedes aegypti, Aedes albopictus* | *NA* | NA | Adults and immature stages (larvae) | Abundance of mosquitoes | Parks and residential areas Shade categories : vegetation shade, under eaves, open outdoor sites | *Aedes albopictus* larvae abundance was positively associated with the parks and vegetation shade.The abundance of *Aedes aegypti* larvae was higher in urban parks than in residential areas, ans was positively related to indoor habitats and adjacent residential areas. Most larval breeding sites in the parks were not found on plants but in vegetation shade. The density of *Aedes aegypti* adults was more significant at all distances in the residential areas compared with the parks. The thickness of *Aedes albopictus* adults was higher in the parks than in residential areas. |
| (de Jesús Crespo et al. 2022) | USA (New Orleans) | Mosquitoes | *Aedes aegypti, Aedes albopictus* | NA | NA | Immature stages (larvae) | Abundance | Remote Sensing: 7 cemeteries with a buffer of 200m around for Urban Heat Island and vegetation cover. Observation: for breeding site, measure of the tree canopy cover. | *Ae. aegypti* is a more abundant species than *Ae. albopictus* in the cemeteries. *Ae. aegypti* was most closely associated with locations with a high cemetery heat index and low vegetation cover. It prefers to be under a shaded canopy in the high-heat cemeteries to the oviposition. The distribution of *Ae. albopictus* shows, not significantly, the opposite trend: the species was present in low-heat cemeteries. According to the authors, the temperature was the main factor explaining the distribution of *Ae. albopictus.* |
| (Yang et al. 2019) | USA (Cleveland) | Mosquitoes | *Culex spp. Aedes japonicus, Aedes vexans, Aedes triseriatus, Aedes trivittatus, Aedes albopictus, Anopheles punctipennis, Orthopodomia signifera, Uranotaenia sapphirina, Coquillettidia perturbans* | WNV | NA | Adults | Mosquito abundance and WNV mosquito infection | For vacant lots:  Two different treatments: mowing/control  measure of the vegetation biomass, dominant plant species diversity, total bloom area     Land cover classes (buffer of 60 m and 1000m): Grass and shrubs, Buildings, Impervious Surface, Tree Canopy over Vegetation, and Tree Canopy over Impervious Surface | A greater vegetation diversity within a vacant lot was positively associated with an increased abundance of *Aedes* and of *Culex.*  *Aedes* and *Culex* mosquito abundances did not differ between mowed treatments.  Vegetation biomass also positively influenced *Aedes* abundances within light traps. There was no significant relationship between bloom area and mosquito abundance.  Partial support that mosquito abundance is higher in greener landscapes (around a vacant lot) depends on scale and species.  WNV was found in *Culex spp.* |
| (Chowdhury et al. 2014) | Bangladesh (Dhaka) | Mosquitoes | *Aedes albopictus, Aedes spp.* | NA | NA | Immature stages (eggs) | Presence and abundance of *Aedes* in tree hole | Tree species (sample of tree holes) | *Delonix regia* and *Mangifera indica* are the trees that have most of the tree holes with the presence of *Aedes albopictus* eggs. There is a positive correlation between the presence of *Aedes albopictus* eggs and the presence of other Aedes eggs. The size of the tree holes and the type of debris can influence the mosquito's position, with an advantage for the older trees. |
| (Lourenço-de-Oliveira et al. 2004) | Brazil (Rio de Janeiro) | Mosquitoes | *Aedes aegypti, Aedes albopictus, Limatus durhami, Culex dolosus* | NA | NA | Immature stages (eggs) / Immature stages (eggs and larvae) | Abundance | Gradient residential to forest | *Aedes aegypti* and *Aedes albopictus* mosquitoes were more abundant near houses than deeper in the forest. However, *Aedes albopictus* was found in abundance even up to 1,000m into the woods. Conversely, *Culex dolosus* was more abundant deeper in the forest than near houses. *Limatus durhami* was more abundant near houses than in the forest. |
| (Andreo et al. 2021) | Argentina (Cordoba) | Mosquitoes | *Aedes aegypti* | NA | NA | Immature stages (eggs and larvae) | Mosquito abundance | NDVI, NDWI and land classification | There is a positive association between mosquito abundance and built neighborhoods but a negative one with vegetated urban and commercial areas. Important variables included distance to built surfaces and to vegetated areas. |
| (Wong et al. 2017) | China (Hong-Kong) | Mosquitoes | *Aedes aegypti, Aedes albopictus, Aedes japonicus, Anopheles spp, Culex quinquefasciatus, Culex tritaeniorhynchus* | NA | NA | Adults | Mosquito abundance | 3 categories:   - Green Roofs  - Bare roofs   - Low-elevation gardens | The abundance was lower in rooftops compared to low-elevation gardens. There was a positive correlation between vector abundance and the mean air temperature and a negative correlation between the mean wind speed and the mean diurnal temperature variation. |
| (Chen et al. 2020) | China (Guangzhou and Foshan) | Mosquitoes | *Aedes aegypti*, Aedes albopictus** | DENV* | Human | NA | Dengue Human cases | green area, % of forest coverage + environmental socioecological variables | A positive correlation exists between green area/forest coverage and dengue human cases. Green areas attract many people, allowing contact between humans and mosquitoes. Temperature, humidity, forest coverage, and green areas are natural environmental factors that contribute to the incidence of dengue fever. |
| (Manica et al. 2016) | Italy (Rome) | Mosquitoes | *Aedes albopictus* | NA | NA | Adults | Mosquito abundance | Land cover : ‘artificial surfaces’ (including ‘roads/concrete’ and ‘buildings’) and ‘vegetation cover’ (including ‘woods’, ‘shrubs’ and ‘grasslands’) | Positive correlation of the abundance with the vegetation cover in a 20 m buffer in an urban environment, contrary to the peri-urban and rural environment. All environments have a negative correlation between vegetation cover in a 300 m buffer and the vector abundance. |
| **TICKS** | | | | | | | | | |
| (Heylen et al. 2019) | Belgium (Antwerp) | Ticks | *Ixodes frontalis, Ixodes hexagonus, Ixodes ricinus* | *Borrelia, Anaplasma phagocytophilum, Babesia spp., Candidatus Neoehrlichia mikurensis* | Roe deer (to calculate connectivity) | larvae, nymph and adult | Tick density and pathogen tick infection | Landcover, vegetation height and green space connectivity | The density of *Ixodidae* increases with the degree of connectivity between green spaces. When ticks are present, the *Borrelia* infection has no specific association. |
| (Kybicová et al. 2017) | Czech Republic (Prague) | Ticks | *Ixodes ricinus* | *Borrelia burgdorferi, Anaplasma phagocytophilum* | NA | nymph and adult | Pathogen tick infection | Types of green areas : one in urban, one in suburban and one in rural area | There is no association between the urban park and the *Borrelia* infection. However, the infection of *Anaplasma phagocytophilum* is higher in urban parks than in peri-urban parks. |
| (Hansford et al. 2017) | England (Salisbury) | Ticks | *Ixodes ricinus* | *Borrelia* | Woodland birds (e.g. blackbird, robin), wood mice, bank voles, foxes, grey squirrels and hedgehogs, roe deer (to compute the variable : the pres- ence/absence of any signs of host activity) | nymph | Tick abundance and pathogen tick infection | Vegetation landcover: grassland, hedge, park, woodland and woodland edge. | *Ixodes ricinus* are more abundant in woodland edge habitats than in grassland, hedges, parks, and woodland sites, even if ticks are present in each of the five habitats. *Borrelia i*nfection does not vary depending on habitat. |
| (Hansford et al. 2021) | England (London) | Ticks | *Ixodes ricinus* | *Borrelia* | NA | larvae, nymph and adult | Tick density and pathogen tick infection | Vegetation land cover: 5 categories in 9 parks: bracken, tall-sward grassland, short-sward grassland, under and woodland. | The tick density is the highest in urban woodland, followed by urban canopy and long and short grass. The density of ticks infected by *Borrelia* is the highest in the urban canopy, followed by the urban woodland, the long and short grass. |
| (Hansford et al. 2022) | England (Bath) | Ticks | *Ixodes ricinus* | *Borrelia* | NA | larvae, nymph and adult | Tick density and pathogen tick infection | Vegetation landcover (field observation): 4 categories: grassland park woodland woodland edge | The presence of ticks is more probable on the woodland edge, in woodland, and in grass than in urban parks. The abundance followed the same pattern. The density of ticks infected is higher on the woodland edge than in the urban grass. The density of *Borrelia*-infected nymphs is highest in woodland and woodland edge compared to grassland and park (borderline significant). |
| (Juntila et al. 1999) | Finland (Helsinki) | Ticks | *Ixodes ricinus* | *Borrelia* | small rodents (to compute null hypothesis) | larvae, nymph and adult | Tick abundance and pathogen tick infection | Types of parks | The tick density and *Borrelia* tick infection are variable in the different urban green spaces studied. The risk of contracting a *Borrelia* infection can be present in an urban environment. The low number of concurrent infections suggests that *Borrelia afzelii* and *Borrelia garinii* favor two distinct reservoir animal populations. |
| (Mathews-Martin et al. 2020) | France (Lyon) | Ticks | *Ixodes ricinus* | *NA* | NA | nymph and adult | Tick abundance | Types of parks (natural, suburban and urban) and vegetation landcover: Forest and footpath, closed environment, forest edge as a transitional environment, and footpath/track in an open area and meadow as open environments. | Tick abundance was higher in the peri-urban parks than in urban parks. In peri-urban parks, ticks were mainly collected on forest or on forest edges. |
| (Schorn et al. 2011a) | Germany (Munich, Regensburg, Ingolstadt, Augsburg and Berg) | Ticks | *Ixodes ricinus* | *Babesia, Rickettsia, Anasplasma phagocytophilum, Bartonella* | NA | larvae, nymph and adult | Pathogen tick infection | Type of parks: intra-urban city parks and suburban recreational areas | The *Babesia* tick infection does not vary in green urban spaces, while the *Rickettsia* tick infection does. Because of the presence of hosts, tick-borne pathogens can occur in recreational urban areas. |
| (Hauck et al. 2020) | Germany (Hanover) | Ticks | *Ixodes ricinus, Ixodes frontalis, Ixodes hexagonus, Ixodes inopinatus,* | *NA* | NA | larvae, nymph and adult | Tick abundance | Type of green areas (field observation): urban parks, mixed forests and broad-leaved forests within city | There is no difference in the tick abundance between the urban mixed and deciduous forests, the deciduous forest, and the park. However, tick density is significantly lower in urban parks than in urban mixed forests. |
| (Overzier et al. 2013) | Germany (Munich, Regensburg) | Ticks | *Ixodes ricinus* | *Rickettsia spp., Babesia* | NA | larvae, nymph and adult | Pathogen tick infection | Type of green areas: urban parks | *Babesia* tick infection is higher in natural areas compared to urban parks. The *Rickettsia* tick infection varies between urban parks, but is higher in natural areas compared ti urban parks. It most be linked to the habitat structure and availability of mammals. |
| (May et al. 2014) | Germany (Hamburg) | Ticks | *Ixodes ricinus* | *Anaplasma phagocytophilum, Rickettsia* | NA | nymph and adult | Pathogen tick infection | Type of parks: different recreational areas | The percentage of infected ticks by *Anaplasma phagocytophilum* and by *Rickettsia* per sampling site varied significantly, but with no explanation given by the authors |
| (Schorn et al. 2011b) | Germany (Munich, Regensburg, Ingolstadt, Augsburg and Berg) | Ticks | *Ixodes ricinus* | *Anaplasma phagocytophilum* | *NA* | nymph and adult | Pathogen tick infection | Type of parks: urban parks in different cities | The infection of *Anaplasma phagocytophilum* can vary between urban parks. That depends of geographical, climatic and biological factors. |
| (Hornok et al. 2014) | Hungary (Budapest) | Ticks | *Ixodes ricinus, Haemaphysalis concinna Dermacentor reticulatus Dermacentor marginatus* | *Anaplasma phagocytophilum, Borrelia.* | NA | larvae, nymph and adult | Tick abundance and pathogen tick infection | Type of green areas: parks, cemeteries and forests | There is no association between the tick density and the different urban green spaces, except for the occasional higher density in the cemetery than in the park and the forest. *Ixodes ricinus* and *Haemaphysalis concinna* were collected in the three biotopes, while *Dermacentor reticulatus* was only collected in parks and forests. There is a positive association between the urban park and the *Anaplasma phagocytophilum* infection but no association with the forest and the cemetery. A positive association exists between the urban forest and the *Borrelia* infection, but no association exists between the *Borrelia* infection and the cemeteries and the urban parks. |
| (Di Luca et al. 2013) | Italy (Rome) | Ticks | *Ixodes ricinus, Dermacentor marginatus, Haemaphysalis punctata, Rhipicephalus bursa and Rhipicephalus turanicus* | *NA* | NA | nymph and adult | Presence and diversity of ticks | Type of green areas: pasture, deciduous forests and ecotonal areas | There is a significant link between the composition of the tick population and the biotope. *Rhipicephalus turanicus* and *Dermacentor marginatus* are more prevalent in pastures than in deciduous mixed wood and ecotonal areas, and *Ixodes ricinus* is more prevalent in deciduous mixed wood than in ecotonal areas and pastures. |
| (Welc-Falęciak et al. 2014) | Poland (Warsaw) | Ticks | *Ixodes ricinus* | *Anasplasma phagocytophilum, Rickettsia, Ehrlichia, Candidatus Neoehrlichia mikurensis* | NA | nymph and adult | The density of ticks and density of infected nymph | Type of green areas: urban and natural areas | Tick abundance varies throughout the urban forest. The same pattern exists for the *Rickettsia* infection. There is no difference for the *Anaplasma phagocytophilum* infection. Tick abundance is lower in urban forests than in natural habitats, while pathogen infection is higher in urban forests compared to natural habitats. |
| (Kubiak et al. 2019) | Poland (Olsztyn) | Ticks | *Ixodes ricinus* | *Borrelia* | NA | nymph and adult | Tick abundance and pathogen tick infection | Type of green areas: 3 urban green areas with 3 different Anthropopressure degree | The abundance of ticks did not vary between the different urban green spaces. *Ixodes ricinus* infected with *Borrelia* were detected in from all study sites with no difference between collection sites |
| (Gryczynska et al. 2021) | Poland (Warsaw) | Ticks | *NA* | *Borrelia* | striped field mouse  yellow-necked mouse  wood mouse | NA | Pathogen host prevalence | Type of green areas: green spaces situated strictly in the city centre and surrounding suburban areas | This study confirms the infection of city-inhabiting rodents, which is lower in green areas located in the city center than in gren areas in suburbs. |
| (Michalik et al. 2003) | Poland (Poznan ́) | Ticks | *Ixodes ricinus* | *Borrelia* | *A. flavicollis, A. sylvaticus, C. glareolus, and Microtus arvalis* | larvae, nymph and adult | Pathogen host prevalence and density of infected nymph | Type of vegetation: deciduous, mixed and coniferous forests | The most optimal environmental conditions for ticks occur in deciduous and mixed forests. |
| (Borsan et al. 2020) | Romania (Cluj-Napoca) | Ticks | *Ixodes ricinus, Haemaphysalis punctata* | *NA* | hedgehogs, birds, micromammals, domestic mammals | larvae, nymph and adult | Presence of hosts and tick abundance | Type of green areas: park, woodland, woodland edge, grassland and hedge + estimation vegetation cover : shrub, arboreal and perennial grass cover (%) | The abundance of *Ixones ricinus* and *Haemaphysalis punctata* in urban parks depends of the sampling site. The density of *Ixodes ricinus* was higher in urban gardens and parks compared to peri-urban forests. There was a significant correlation between the daily mean temperature of campaign and the tick abundance. There was no association between tick abundance and the vegetation cover. The difference can be explained by the presence of hedhehogs which can increase ticks abundance. |
| (Tretyakov et al. 2009) | Russia (St. Petersburg) | Ticks | *Ixodes persulcatus, Ixodes ricinus, Ixodes trianguliceps, and Ixodes apronophorus* | *NA* | Small mammals (Common shrew, Pygmy shrew, Laxmannís shrew, Ural field mouse, Black-striped field mouse, Yellow-necked mouse, House mouse, Bank vole, Common vole) | larvae and nymph | Presence of hosts and tick diversity and tick abundance | Type of parks: small suppressed parks, landscape parks and forest parks | *Ixodes persulcatus*, *Ixodes trianguliceps* and *Ixodes apronophorus* were found in small mammals in the different parks investigated. *Ixodes ricinus* ware rare. The difference between the parks can be explained by the adjacent territories. Large forestlands allow the presence of medium-sized and large mammals, feeding imagos while parks bordering with built-up areas do not allow the presence of large wild mammals. |
| (Kwak et al. 2021) | Singapore (Singapore) | Ticks | *Amblyomma helvolum,Dermacentor auratus, Haemaphysalis semermis, Ixodes granulatus* | *NA* | Small mamals (Muridae, Sciuridae, Soricidae, Tupaiidae) | adult | Host presence and tick abundance | Vegetation landcover: Old secondary forests,Young secondary forests, Scrublands Urban habitat | There is a statistically significant difference in infestation rate between old secondary and young secondary forest but not between old secondary forest and scrubland, scrubland and young secondary forest. |
| (Levytska et al. 2021) | Ukraine (Chernivtski, Khmelnytski, Kyiv, Ternopil and Vinnytsia) | Ticks | *Ixodes ricinus, Ixodes hexagonus, Dermacentor reticulatus* | *Anaplasma phagocytophilum, Rickettsia, Babesia, Bartonella, Borellia* | wild and domestic animals | nymph and adult | Pathogen tick infection | Type of parks | *Dermacentor reticulatus* is the most abundant tick collected, *Ixodes ricinus* the second, and *Ixodes hexagonus* the last. All pathogens are found in all parks. *Borrelia* is present only in *Ixodes ricinus*. There is no difference on pathogen tick infection between parks and cities. |
| (Adalsteinsson et al. 2012) | USA (New Castle County) | Ticks | *Ixodes dentatus, Ixodes scapularis, Haemaphysalis leporispalustris* | *Borrelia, Anaplasma phagocytophilum, Babesia* | Mice | nymph | Density of infected nymph | Vegetation landcover: % of ground covered by R. multiflora, by coarse woody debris, and density of understory vegetation + land use variable | Infection in ticks from forests invaded by *Rosa multiflora* is almost double that in uninvaded woods. The density of ticks infected is higher in woody debris than in leaf litter. The total understory cover is also positively associated with tick infection. *Rosa multiflora* invasion and the additional factors positively influencing pathogen infection point to suitable habitat characteristics for small mammal and bird hosts that are competent pathogen reservoirs |
| (VanAcker et al. 2019) | USA (New York) | Ticks | *Ixodes scapularis* | *Borrelia* | white-tailed deer* (connectivity factors) | nymph | Density of tick and pathogen tick infection | Land cover (Remote Sensing) : tree canopy, grass/shrub, bare soil, water, buildings, roads, and other paved surfaces | The distance between parks best explains the presence and abundance of *Ixodes scapularis*. Connectivity of parks for deer and other hosts, had the largest positive effect on the density and infection of *Ixodes scapulari*s nymphs. Covariates that describe the landscape composition surrounding each park also have a significant positive (percentage tree canopy) or negative (percentage of grass and shrubs and and bare soil) effect on the densities of nymphs. Sometimes, the percentage of grass and shrub does not influence the tick density. |
| (Small et al. 2021) | USA (Oklahoma) | Ticks | *Amblyomma americanum* | *Rickettsia amblyommatis, Ehrlichia chaffeensis* | NA | nymph and adult | Pathogen tick infection | Type of parks: residential zoning and agricultural zoning | There are no significant differences in bacteria infection between the two parks, located in a residential zoning and in an agricultural zoning. |
| (Roselli et al. 2021) | USA (Oklahoma) | Ticks | *Amblyomma americanum, Amblyomma maculatum, Haemaphysalis leporispalustris* | *NA* | Birds | larvae and nymph | Tick abundance on bird | Land cover: Urbanization gradient | Non-migratory bird species and migratory bird species during their breeding season are important carriers of ticks on urban parks and urban green spaces. The tick infestation for all birds (Northern Cardinal, Carolina Wren) combined declined with increasing urbanization intensity. |
| (Noden et al. 2022) | USA (Oklahoma) | Ticks | *Amblyomma americanum, Amblyomma maculatum, Dermacentor variabilis* | *Rickettsia spp., Babesia, Hepatozoon, Theileria, Ehrlichia and Anaplasma spp., Borrelia* | *NA* | nymph and adult | Density of infected nymph | Land cover: Urbanization gradient | The surrounding landscape does not seem to influence the *Rickettsia* and *Borrelia* tick infection. For *Ehrlichia* and *Babesia*, the percentage of urbanization of the surroundinf landscape can be negatively associated with tick infection or can have no effect. That depends if the percentage of urbanization is used as a categorical or a continuous variable. |
| (Hasset et al. 2022) | USA (New York) | Ticks | *Ixodes scapularis, Amblyomma americanum, Haemaphysalis longicornis* | *NA* | Human behaviour | larvae, nymph and adult | Host and tick presence | Field land cover: maintained grass, unmain tained herbaceous, leaf litter, impervious, and edge | People’s KAPs do not change across parks, even if parks represent different exposure risks. Tick density is positively associated with urban parks. There is less ticks in maintained grass and in edge than in unmaintained herbaceous areas and in trails. The contact between ticks and humans is higher in trail than in open spaces. |
| (Gregory et al. 2022) | USA (New York) | Ticks | *Ixodes scapularis, Amblyomma americanum, Haemaphysalis longicornis* | *NA* | *NA* | *nymph* | Tick presence | Vegetation land cover: rass, shrub, low canopy, high canopy, bare soil, water and impervious surface + landscape metrics | The risk of tick exposure in residential yards is considerable. The dynamics of three tick vector species in a highly fragmented urban environment are determined by yard- and landscape-level features (yard permeability, land cover). The effect depends on tick species. *Amblyomma americanum* is positively associated with log and brush piles in yards, permeable edge, a high canopy cover and no associated with vegetable of flower gardens and a low canopy cover and grass around the yard. *Ixodes scapularis* is positively associated with log and brush, high canopy cover and no associated with a permeable edge, a low canopy cover and grass around yard. *Haemaphysalis longicornis* is only positively associated with the presence of log and brush piles. |
| **SANDFLIES** | | | | | | | | | |
| (Berrozpe et al. 2017) | Argentina (Corientes) | Sandflies | *Lutzomyia longipalpis* | *Leishmania infantum* | NA | Adult | Sanflies presence, Sandflies abundance and Sadnflies diversity | 2 categories (5 classes in each : urban, bare soil, water, high vegetation and low vegetation):  - urban  - peri-urban   - rural islands | Vector abundance is associated with farmyard animals and low NDVI. |
| **CHIGGER MITES** | | | | | | | | | |
| Park et al. 2015 | South Korea (Seoul) | Chigger mites | *Leptotrombidium species: Helenicula miyagawai, L. scutellare, L. zetum and L. palpale* | *Orientia tsutsugamushi* | Human | Adults | Scrub typhus human disease, abundance of chigger mites and presence of O. tsutusgamushi in vectors | 3 biotopes level in city: nayural area, partially-exploited and agricultural farms/city parks | The Human scrub typhus prevalence in the city park was lower than in the city in mountainous areas and higher than in residential areas. Some chigger mites infected with *Orientia tsutsugamushi* were present in Seoul. |
| Wulandhari et al. 2021 | Thailand (Bangkok) | Chigger mites | *Leptotrimbidium deliense, Ascoschoengastia indica* | Orientia tsutsugamushi | Small rodents (*Rattus rattus, Rattus exulans, Rattus norvegicus, Tupaia belangeri*) | Adults | Infestation of vectors on host, Pathogen vector infection | Land use types: (1) post-flooding/rain-fed land, (2) mosaic cropland/vegetation, (3) shrub/grassland, (4) close to a mixed forest and (5) artificial surface/urban within a certain park  + 4 categories of surrounding parks: (1) water body); (2) open field, (3) shaded tree and (4) building. | Urban public parks provide a suitable habitat for chigger species to survive since they exploit the available hosts to maintain their population in a big city. There was a higher chigger abundance in open field habitat. |
