## Additional 5 Table S2 for "Green cities and the risk for vector-borne disease transmission for humans and animals: a scoping review"

| **TYPE OF DESCRIPTION** | **GROUP OF DESCRIPTION** | **CATEGORY** | **SUBCATEGORY** | **VECTOR** | **EFFECT OF URBAN VEGETATION (and REFERENCES)** |
| --- | --- | --- | --- | --- | --- |
| ***Urban green space typology*** | Urban green space | Park | | *Aedes spp.* | Harmful (Schwarz et al. 2020) |
|  |  |  |  | *Aedes aegypti* | Harmful (Huynh et al. 2022), Beneficial (Huynh et al. 2022) |
|  |  |  |  | *Aedes albopictus* | Harmful (Huynh et al. 2022, Lee et al. 2020, Schwarz et al. 2020) |
|  |  |  |  | *Aedes hendersoni* | No association (Schwarz et al. 2020) |
|  |  |  |  | *Aedes triseriatus* | Harmful (Schwarz et al. 2020) |
|  |  |  |  | Various genus combined | Harmful (Medeiros-Sousa et al. 2017, Wilke et al. 2017) |
|  |  | Forest | | *Aedes spp.* | Harmful (Schwarz et al. 2020) |
|  |  |  |  | *Aedes aegypti* | Beneficial (Cox et al. 2007) |
|  |  |  |  | *Aedes albopictus* | Harmful (Schwarz et al. 2020) |
|  |  |  |  | *Aedes hendersoni* | Harmful (Schwarz et al. 2020) |
|  |  |  |  | *Aedes triseriatus* | Harmful (Schwarz et al. 2020), No association (Schwarz et al. 2020) |
|  |  |  |  | *Culex antillummagnorum* | Harmful (Cox et al. 2007) |
|  |  |  |  | *Culex quinquefasciatus* | Beneficial (Cox et al. 2007) |
|  |  |  |  | *Haemagogus leucocelaenus, Haemagogus janthinomys/capricornii* | No association (de Abreu et al. 2022) |
|  |  |  |  | Various genus combined | Harmful (Vieira et al. 2020) |
|  |  | Residential areas | Residential areas | *Aedes aegypti* | Harmful (Cox et al. 2007) |
|  |  |  |  | *Aedes mediovittatus* | Harmful (Cox et al. 2007) |
|  |  |  | Low residential areas density | *Culex antillummagnorum* | Harmful (Cox et al. 2007) |
|  |  |  |  | *Culex quinquefasciatus* | Harmful (Cox et al. 2007) |
|  |  |  | Moderate vegetated residential areas | *Aedes vexans* | Harmful (Brown et al. 2008) |
|  |  |  |  | *Culex pipiens* | Harmful (Brown et al. 2008) |
|  |  |  |  | *Culex restuans* | Harmful (Brown et al. 2008) |
|  |  |  | Highly vegetated residential areas | *Culex salinarius* | Harmful (Brown et al. 2008) |
|  |  | Green roofs | | *Aedes albopictus* | No association (Wong et al. 2018) |
|  |  |  |  | *Aedes japonicus* | No association (Wong et al. 2018) |
|  |  |  |  | *Culex quinquefasciatus* | Beneficial (Wong et al. 2018) |
|  |  |  |  | Various genus combined | Beneficial (Wong et al. 2016, Wong et al. 2017, Wong et al. 2018) |
|  |  | Urban agriculture | sugar cane | *Anopheles spp.* | Harmful (Dongus et al. 2009) |
|  |  |  | leafy vegetables | *Anopheles spp.* | Harmful (Dongus et al. 2009) |
|  |  |  | rice cover | *Culex spp.* | Harmful (Ha et al. 2021) |
|  |  |  | crop diversity | *Anopheles spp.* | Harmful (Dongus et al. 2009), Beneficial (Dongus et al. 2009) |
|  |  |  | urban agriculture | *Anopheles spp.* | Harmful (Dongus et al. 2009) |
|  |  | Urban farms | | *~~Aedes albopictus~~* | ~~Harmful (Wilke et al. 2020)~~ |
|  |  |  |  | Various genus combined | Harmful (Lee et al. 2020, ~~Wilke et al. 2020~~) |
|  |  | Generic urban green spaces | | *Aedes aegypti* | Beneficial (Lorenz et al. 2020, Pedrosa et al. 2020) |
|  | Characteristics of urban green space | Specific area within urban green space | Distance from edge within the park | *Aedes aegypti* | Beneficial (Heinisch et al. 2019, Lourenço-de-Oliveira et al. 2004) |
|  |  |  |  | *Aedes albopictus* | Harmful (Heinisch et al. 2019), No association (Hendy et al. 2020), Beneficial (Lourenço-de-Oliveira et al. 2004) |
|  |  |  |  | *Culex dolosus* | Harmful (Lourenço-de-Oliveira et al. 2004) |
|  |  |  |  | *Limatus durhami* | Beneficial (Lourenço-de-Oliveira et al. 2004) |
|  |  |  |  | *Phosphora amazonica* | No association (Hendy et al. 2020) |
|  |  |  |  | *Trichoprosopon digitatum* | No association (Hendy et al. 2020) |
|  |  |  |  | Various genus combined | No association (Hendy et al. 2020) |
|  |  | Management | Mown treatments | *Aedes spp.* | No association (Yang et al. 2019) |
|  |  |  |  | *Culex spp.* | No association (Yang et al. 2019) |
|  |  | Urban agricultural trenches | | *Anopheles spp.* | Harmful (Matthys et al. 2010) |
|  |  | Urban irrigation well presence | | *Anopheles spp.* | Harmful (Dongus et al. 2009) |
|  |  | ~~Distance to an highly urbanized area~~ | | *~~Culex quinquefasciatus~~* | ~~Harmful (Wilke et al. 2020)~~ |
|  | ~~Distance~~ Proximity to urban green spaces | | | *Aedes spp.* | Harmful (Talbot et al. 2021) |
|  |  |  |  | *Aedes albopictus* | Harmful (Pedrosa et al. 2020) |
|  |  |  |  | *Anopheles gambiae s.l.* | Harmful (Yadouléton et al. 2010) |
| ***Urban vegetation description*** | Type of vegetation | Low vegetation | Ground vegetation | *Aedes albopictus* | No association (Hendy et al. 2020) |
|  |  |  | Herbaceous | *Aedes aegypti* | No association (Landau et al. 2012) |
|  |  |  |  | *Culex quinquefasciatus* | Beneficial (Landau et al. 2012), No association (Landau et al. 2012) |
|  |  |  | Grass | *Aedes aegypti* | Beneficial (Sun et al. 2021) |
|  |  |  |  | *Aedes albopictus* | Beneficial (Cianci et al. 2015, Sun et al. 2021) |
|  |  |  | Grass and shrub | *Aedes spp.* | Harmful (Yang et al. 2019), No association (Yang et al. 2019) |
|  |  |  |  | *Culex spp.* | Harmful (Yang et al. 2019), No association (Yang et al. 2019) |
|  |  |  | Shrub | *Aedes aegypti* | No association (Landau et al. 2012) |
|  |  |  |  | *Culex pipiens, Culex restuans* | Harmful (Gardner et al. 2013) |
|  |  |  |  | *Culex quinquefasciatus* | Beneficial (Landau et al. 2012), No association (Landau et al. 2012) |
|  |  |  | Flowering shrub | *Culex pipiens, Culex restuans* | Beneficial (Gardner et al. 2013) |
|  |  |  | non-forested wetlands | *Culex quinquefasciatus* | Harmful (Sallam et al. 2017) |
|  |  |  | Decorative vegetation | *Aedes spp.* | Harmful (Talbot et al. 2021) |
|  |  | High vegetation | High vegetation | *Aedes aegypti* | No association (Landau et al. 2012) |
|  |  |  |  | *Culex quinquefasciatus* | No association (Landau et al. 2012) |
|  |  |  | Arboreal vegetation | *Aedes aegypti* | Beneficial (Benitez et al. 2020) |
|  |  |  | Forest | *Culex spp.* | Beneficial (Ha et al. 2021) |
|  |  |  |  | *Aedes aegypti* | Beneficial (Sun et al. 2021) |
|  |  |  |  | *Aedes albopictus* | Harmful (Sun et al. 2021) |
|  |  |  |  | *Haemagogus spp.* | No association (de Abreu et al. 2022) |
|  |  | Vegetation management | % urban managed vegetation | *Aedes aegypti* | Harmful (Sun et al. 2021) |
|  |  |  |  | *Aedes albopictus* | Harmful (Sun et al. 2021) |
|  | Vegetation index | Percentage of vegetation | | *Aedes spp.* | Harmful (Francisco et al. 2021) |
|  |  |  |  | *Aedes aegypti* | No association (Landau et al. 2012), Beneficial (Arduino et al. 2020) |
|  |  |  |  | *Aedes albopictus* | Harmful (Arduino et al. 2020, Cianci et al. 2015) |
|  |  |  |  | *Culex quinquefasciatus* | No association (Landau et al. 2012) |
|  |  | NDVI | | *Aedes aegypti* | Harmful (Landau et al. 2012), No association (Landau et al. 2012) |
|  |  |  |  | *Aedes albopictus* | Harmful (Hendy et al. 2020) |
|  |  |  |  | *Aedes vexans* | No association (Trawinski et al. 2010) |
|  |  |  |  | *Culex spp.* | No association (Ha et al. 2021) |
|  |  |  |  | *Culex pipiens* | Beneficial (Trawinski et al. 2010), No association (Trawinski et al. 2010) |
|  |  |  |  | *Culex quinquefasciatus* | No association (Landau et al. 2012, Ha et al. 2021) |
|  |  |  |  | *Haemagogus spp.* | Harmful (de Abreu et al. 2022) |
|  |  |  |  | *Phosphora amazonica* | No association (Hendy et al. 2020) |
|  |  |  |  | *Trichoprosopon digitatum* | No association (Hendy et al. 2020) |
|  |  |  |  | Various genus combined | Harmful (Brown et al. 2008) |
|  |  | NDWI | | *Culex spp.* | Harmful (Demets et al. 2020) |
|  |  |  |  | Various genus combined | Harmful (Brown et al. 2008) |
|  |  | Vegetation cover | | *Aedes aegypti* | Beneficial (Honório et al. 2009), No association (de Jesús Crespo et al. 2022), ~~Harmful (de Jesús Crespo et al. 2022)~~ |
|  |  |  |  | *Aedes albopictus* | Harmful (Honório et al. 2009, Cianci et al. 2015, Manica et al. 2016), No association (de Jesús Crespo et al. 2022), Beneficial (Manica et al. 2016) |
|  |  |  |  | *~~Culex quinquefasciatus~~* | ~~Beneficial (de Jesús Crespo et al. 2022)~~ |
|  |  | Tree cover | | *Aedes aegypti* | Harmful (Landau et al. 2012,de Jesús Crespo et al. 2022 ), No association (Landau et al. 2012) |
|  |  |  |  | *Aedes albopictus* | Harmful (Cianci et al. 2015), No association (Hendy et al. 2020) |
|  |  |  |  | *Culex quinquefasciatus* | No association (Landau et al. 2012) |
|  | Vegetation characteristics | Canopy level | | *Aedes albopictus* | Harmful (Schwarz et al. 2020)~~,~~ ~~No association (Hendy et al. 2020)~~ |
|  |  |  |  | *Aedes triseriatus, Aedes hendersoni* | Beneficial (Schwarz et al. 2020) |
|  |  | Ground vegetation | | *Aedes albopictus* | No association (Hendy et al. 2020) |
|  |  | Presence of vegetation in breeding site | | *Anopheles spp.* | Beneficial (Matthys et al. 2010) |
|  |  | Tree height | Low tree height | *Aedes aegypti* | Beneficial (Landau et al. 2012), No association (Landau et al. 2012) |
|  |  |  |  | *Culex quinquefasciatus* | No association (Landau et al. 2012) |
|  |  |  | Medium tree height | *Aedes aegypti* | Harmful (Landau et al. 2012) |
|  |  |  |  | *Culex quinquefasciatus* | Harmful (Landau et al. 2012), No association (Landau et al. 2012) |
|  |  |  | High tree height | *Aedes aegypti* | Harmful (Landau et al. 2012), No association (Landau et al. 2012) |
|  |  |  |  | *Culex quinquefasciatus* | Harmful (Landau et al. 2012), No association (Landau et al. 2012) |
|  |  | Vegetation shade | | *Aedes aegypti* | Beneficial (Huynh et al. 2022) |
|  |  |  |  | *Aedes albopictus* | Harmful (Huynh et al. 2022) |
|  |  | Vegetation diversity | | *Aedes spp.* | Harmful (Yang et al. 2019) |
|  |  |  |  | *Culex spp.* | Harmful (Yang et al. 2019), No association (Yang et al. 2019) |
|  |  | Vegetation biomass | | *Aedes spp.* | Harmful (Yang et al. 2019) |
|  |  |  |  | *Culex spp.* | No association (Yang et al. 2019) |
|  |  | Bloom area | | *Aedes spp.* | No association (Yang et al. 2019) |
|  |  |  |  | *Culex spp.* | No association (Yang et al. 2019) |
|  |  | Land use below the canopy | Tree canopy over vegetation | *Aedes spp.* | Harmful (Yang et al. 2019), No association (Yang et al. 2019) |
|  |  |  |  | *Culex spp.* | No association (Yang et al. 2019) |
|  |  |  | Tree canopy over impervious surface | *Aedes spp.* | Harmful (Yang et al. 2019), No association (Yang et al. 2019) |
|  |  |  |  | *Culex spp.* | No association (Yang et al. 2019) |
|  |  | Tree density | Tree density | *Culex quinquefasciatus* | Harmful (Sallam et al. 2017) |
|  |  |  | Decideous tree density | *Culex pipiens, Culex restuans* | No association (Gardner et al. 2013) |
|  |  | Flower vases | | *Aedes albopictus* | Harmful (Che Dom et al. 2016) |
|  |  | Tree holes size | | *Aedes spp.* | Variable (Chowdhury et al. 2014) |
|  | Plant species | maple | | *Culex pipiens, Culex restuans* | Variable (Gardner et al. 2013) |
|  |  | oak, arborvitae, spruce, elm, pear, birch density | | *Culex pipiens, Culex restuans* | Harmful (Gardner et al. 2013) |
|  |  | ash | | *Culex pipiens, Culex restuans* | Beneficial (Gardner et al. 2013) |
|  |  | palm tree | | *Aedes albopictus* | No association (Hendy et al. 2020) |
|  |  | *Abelia grandiflora* | | *Aedes albopictus* | Harmful (Tian et al. 2019) |
|  |  | *Delonix regia* | | *Aedes spp.* | Harmful (Chowdhury et al. 2014) |
|  |  | *Ligustrum quihoui* | | *Aedes albopictus* | Harmful (Tian et al. 2019) |
|  |  | *Mangifera indica* | | *Aedes spp.* | Harmful (Chowdhury et al. 2014) |
|  |  | *Nandina domestica* | | *Aedes albopictus* | Beneficial (Tian et al. 2019) |
|  | Proximity to urban vegetation | | | *Aedes spp.* | Harmful (Honório et al. 2009, Paul et al. 2018, Che Dom et al. 2016) |
|  |  |  |  | *Aedes aegypti* | Harmful (Estallo et al. 2018), Beneficial (Andreo et al. 2021), Variable (Andreo et al. 2021), No association ( Andreo et al. 2021) |
