## Additional 6 Table S3 for "Green cities and the risk for vector-borne disease transmission for humans and animals: a scoping review"

| **TYPE OF DESCRIPTION** | **GROUP OF DESCRIPTION** | **CATEGORY** | **SUBCATEGORY** | **VECTOR BORNE DISEASE (Pathogen)** | **VECTOR** | **EFFECT OF URBAN VEGETATION (and REFERENCES)** |
| --- | --- | --- | --- | --- | --- | --- |
| ***Urban green space typology*** | Type of urban green space | Park | | West Nile Fever (WNV) | *Culex spp* | No association (Levine et al. 2016, Levine et al. 2017) |
|  |  |  |  |  | *Various genus combined* | No association (Talbot et al. 2019) |
|  |  | Forest | | Yellow fever (YFV) | *Haemagogus spp.* | Harmful (de Abreu et al. 2022) |
|  |  |  |  | West Nile Fever (WNV) | *Culex spp* | Beneficial (Levine et al. 2017, Levine et al. 2016), No association (Levine et al. 2017, Levine et al. 2016) |
|  |  | Residential areas | Residential areas | West Nile Fever (WNV) | *Culex spp* | Harmful (Levine et al. 2017), No association (Levine et al. 2016, Levine et al. 2017) |
|  |  |  | Low residential areas density | Saint-Louis Encephalitis (SLE) | *Culex quinquefasciatus* | Harmful (Vergara Cid et al. 2013) |
|  |  | Zoo | | West Nile Fever (WNV) | *Culex spp* | No association (Levine et al. 2016), Beneficial (Levine et al. 2016) |
|  |  | Urban agriculture | Urban mixed crop systems | Malaria (*Plasmodium falciparum*) | *Anopheles spp.* | Harmful (Matthys et al. 2006) |
|  |  |  | Arable land | Dengue (DENV) | *Aedes aegypti, Aedes albopictus* | Beneficial (Chen et al. 2020) |
|  |  |  | % of urban agriculture | West Nile Fever (WNV) | *Culex spp* | Harmful (Liu et al. 2008) |
|  |  |  |  | Malaria (*Plasmodium falciparum)* | *Anopheles gambiae s.l.* | Harmful (Yadouléton et al. 2010) |
|  |  | Generic urban green spaces | | West Nile Fever (WNV) | *Culex spp.* | Beneficial (LaBeaud et al. 2008) |
|  |  |  |  |  | *Culex pipiens restuans* | Beneficial (Myer et al. 2019) |
|  | Characteristics of urban green space | Surface area | Urban forest | West Nile Fever (WNV) | *Culex quinquefasciatus* | Beneficial (Lockaby et al. 2016) |
|  |  |  |  | Dengue (DENV) | *Aedes aegypti, Aedes albopictus* | No association (Chen et al. 2020) |
|  |  |  | Wetland | West Nile Fever (WNV) | *Culex spp.* | Harmful (Liu et al. 2008) |
|  |  |  | Green urban area | Dengue (DENV) | *Aedes aegypti, Aedes albopictus* | Harmful (Chen et al. 2020), No association (Chen et al. 2020) |
|  |  | Green spaces in built up areas | | Dengue (DENV) | *Aedes aegypti, Aedes albopictus* | Harmful (Chen et al. 2020) |
|  |  | Proportion of natural areas parks | | West Nile Fever (WNV) | *Culex spp.* | No association (Talbot et al. 2019), Beneficial (Talbot et al. 2019) |
|  |  | Proportion of urban parks | | West Nile Fever (WNV) | *Culex spp.* | No association (Talbot et al. 2019) |
|  |  | Number of park | | Dengue (DENV) | *Aedes aegypti, Aedes albopictus* | Harmful (Chen et al. 2020) |
|  | Distance to urban green spaces | | | Malaria (*Plasmodium falciparum)* | *Anopheles gambiae s.l.* | Harmful (Yadouléton et al. 2010) |
| ***Urban vegetation description*** | Type of vegetation | Forest | Forest | Dengue (DENV) | *Aedes aegypti, Aedes albopictus* | Harmful (Chen et al. 2020) |
|  |  |  | Pine forest | West Nile Fever (WNV) | *Culex quinquefasciatus* | Beneficial (Lockaby et al. 2016) |
|  |  | Riparian | | Malaria (*Plasmodium falciparum*) | *Anopheles gambiae, Anopheles funestus* | Harmful (Kabria et al. 2016) |
|  |  |  |  | Ross River Fever (RRV) | *Aedes vigilax, Culex annulirostris* | Harmful (Muhar et al. 2000) |
|  |  | bushlands | | Ross River Fever (RRV) | *Aedes vigilax, Culex annulirostris* | Harmful (Muhar et al. 2000) |
|  | Vegetation index | percentage of vegetation | | Chikungunya (CHIKV) | *Aedes aegypti* | No association (Freitas et al. 2021), Beneficial (Freitas et al. 2021**)** |
|  |  |  |  | Dengue (DENV) | *Aedes aegypti, Aedes albopictus* | Harmful (Chen et al. 2020), Beneficial (Francisco et al. 2021) |
|  |  | NDVI | | Dengue (DENV) | *Aedes aegypti* | No association (Ogashawara et al. 2019), Beneficial (Troyo et al. 2009) |
|  |  |  |  | West Nile Fever (WNV) | *Culex spp.* | No association (Liu et al. 2012) |
|  |  |  |  |  | *Culex quinquefasciatus* | Beneficial (Poh et al. 2020) |
|  |  |  |  |  | *Culex pipiens restuans* | Beneficial (Myer et al. 2019) |
|  |  |  |  | Malaria (*Plasmodium falciparum)* | *Anopheles gambiae, Anopheles funestus* | Harmful (Kabaria et al. 2016) |
|  |  | NDWI | | Dengue (DENV) | *Aedes aegypti* | No association (Ogashawara et al. 2019) |
|  |  | Weekly Enhanced Vegetation Index | | Dengue (DENV) | *Aedes aegypti* | Harmful (Troyo et al. 2009) |
|  |  | Vegetation cover | Vegetation cover | Dengue (DENV) | *Aedes aegypti* | Harmful (Zellweger et al. 2017), No association (Zellweger et al. 2017) |
|  |  |  | Low vegetation cover | Dengue (DENV) | *Aedes aegypti* | Harmful (Araujo et al. 2015) |
|  |  |  | Medium vegetation cover | Dengue (DENV) | *Aedes aegypti* | Harmful (Araujo et al. 2015) |
|  |  |  | High vegetation cover | Dengue (DENV) | *Aedes aegypti* | Beneficial (Araujo et al. 2015) |
|  |  | Canopy cover | Canopy cover | Dengue (DENV) | *Aedes aegypti* | No association (Troyo et al. 2009) |
|  |  |  |  | Dengue (DENV) | *Aedes aegypti, Aedes albopictus* | Harmful (Chen et al. 2020) |
|  |  |  | High urban canopy density | West Nile Fever (WNV) | *Culex pipiens, Culex restuans* | Harmful (Liu et al. 2011) |
|  | Vegetation characteristics | tall vegetation | | Malaria (Plasmodium falciparum) | *Anopheles gambiae, Anopheles arabiensis, Anopheles funestus* | Beneficial (Georganos et al. 2020) |
|  |  | Tree diameter | | West Nile Fever (WNV) | *Culex quinquefasciatus* | No association (Lockaby et al. 2016) |
|  |  | Vegetation heigh | | West Nile Fever (WNV) | *Culex quinquefasciatus* | No association (Lockaby et al. 2016) |
|  | Distance to urban vegetation | | | Saint-Louis Encephalitis (SLEV) | *Culex quinquefasciatus* | Harmful (Vergara Cid et al. 2013) |
