## Additional 7 Table S4 for "Green cities and the risk for vector-borne disease transmission for humans and animals: a scoping review"

| **TYPE OF DESCRIPTION** | **GROUP OF DESCRIPTION** | **CATEGORY** | **SUBCATEGORY** | **VECTOR** | **EFFECT OF URBAN VEGETATION (and REFERENCES)** |
| --- | --- | --- | --- | --- | --- |
| ***Urban green space typology*** | Type of urban green space | Urban park | | *Ixodes scapularis* | Harmful (Hasset et al. 2022) |
|  |  |  |  | *Ixodes ricinus* | Variable (Borsan et al. 2020, Mathews-Martin et al. 2020), Beneficial (Hauck et al. 2020, Hornok et al. 2014), No association (Borsan et al. 2020, Hauck et al. 2020, Hornok et al. 2014, Tretyakov et al. 2009) |
|  |  |  |  | *Ixodes persulcatus* | Harmful (Tretyakov et al. 2009) |
|  |  |  |  | *Ixodes trianguliceps* | Harmful (Tretyakov et al. 2009) |
|  |  |  |  | *Ixodes apronophorus* | Harmful (Tretyakov et al. 2009) |
|  |  |  |  | *Amblyomma americanum* | Harmful (Hasset et al. 2022), Variable (Hasset et al. 2022) |
|  |  |  |  | *Haemaphysalis longicornis* | Harmful (Hasset et al. 2022), Variable (Hasset et al. 2022) |
|  |  |  |  | *Haemaphysalis punctata* | No association (Borsan et al. 2020), Variable (Borsan et al. 2020) |
|  |  |  |  | *Ixodes inopinatus* | Beneficial (Hauck et al. 2020), No association (Hauck et al. 2020) |
|  |  | Urban forest | | *Ixodes ricinus* | No association (Hornok et al. 2014), Beneficial (Hornok et al. 2014), Variable (Welc-Falęciak et al. 2014) |
|  |  | Urban cemetery | | *Ixodes ricinus* | Harmful (Hornok et al. 2014), No association (Hornok et al. 2014) |
|  |  | Urban agriculture | Urban cropland | *Ripicephalus turanicus* | Harmful (Di Luca et al. 2013) |
|  |  |  |  | *Dermacentor marginatus* | Harmful (Di Luca et al. 2013) |
|  |  | Generic urban green spaces | | *Ixodes ricinus* | Variable (Juntila et al. 1999), No association (Kubiak et al. 2019) |
|  | Characteristics of urban green space | Specific area within green urban spaces | Edge | *Amblyomma americanum* | Beneficial (Hasset et al. 2022) |
|  |  |  |  | *Haemaphysalis longicornis* | Beneficial (Hasset et al. 2022) |
|  |  |  | Trail | *Amblyomma americanum* | Harmful (Hasset et al. 2022) |
|  |  |  |  | *Haemaphysalis longicornis* | Harmful (Hasset et al. 2022) |
|  |  | Managed vegetation | | *Ixodes scapularis* | Beneficial (Hasset et al. 2022) |
|  |  |  |  | *Amblyomma americanum* | Beneficial (Hasset et al. 2022) |
|  |  |  |  | *Haemaphysalis longicornis* | Beneficial (Hasset et al. 2022) |
|  |  | Connectivity | | *Amblyomma americanum* | Harmful (Gregory et al. 2022) |
|  |  |  |  | *Haemaphysalis longicornis* | No association (Gregory et al. 2022) |
|  |  |  |  | *Ixodes scapularis* | No association (Gregory et al. 2022), Harmful (VanAcker et al. 2019) |
|  |  |  |  | *Ixodes ricinus* | Harmful (Heylen et al. 2019) |
|  |  | Permeable edge | | *Ixodes scapularis* | No association (Gregory et al. 2022) |
|  |  |  |  | *Haemaphysalis longicornis* | No association (Gregory et al. 2022) |
|  |  |  |  | *Amblyomma americanum* | Harmful (Gregory et al. 2022) |
|  |  | surrounding land cover | Urban tree canopy | *Ixodes scapularis* | Harmful (VanAcker et al. 2019) |
|  |  |  | Low urban tree canopy cover | *Amblyomma americanum* | No association (Gregory et al. 2022) |
|  |  |  |  | *Haemaphysalis longicornis* | No association (Gregory et al. 2022) |
|  |  |  |  | *Ixodes scapularis* | No association (Gregory et al. 2022) |
|  |  |  | High urban tree canopy | *Amblyomma americanum* | Harmful (Gregory et al. 2022) |
|  |  |  |  | *Haemaphysalis longicornis* | No association (Gregory et al. 2022) |
|  |  |  |  | *Ixodes scapularis* | Harmful (Gregory et al. 2022) |
|  |  |  | Urban grassland and shrubs | *Ixodes scapularis* | Beneficial (VanAcker et al. 2019), No association (VanAcker et al. 2019, Gregory et al. 2022) |
|  |  |  |  | *Amblyomma americanum* | No association (Gregory et al. 2022) |
|  |  |  |  | *Haemaphysalis longicornis* | No association (Gregory et al. 2022) |
|  |  |  | Urban bare soil | *Ixodes scapularis* | Beneficial (VanAcker et al. 2019) |
|  |  |  | % developed lands | *Amblyomma americanum* | Beneficial (Roselli et al. 2021), No association (Roselli et al. 2021) |
|  |  |  |  | *Amblyomma maculatum* | Beneficial (Roselli et al. 2021), No association (Roselli et al. 2021) |
|  |  |  |  | *Haemaphysalis leporispalustris* | Beneficial (Roselli et al. 2021), No association (Roselli et al. 2021) |
| ***Description of urban vegetation*** | Type of vegetation | Forest | Woodland | *Ixodes ricinus* | Harmful (Hansford et al. 2021, Hansford et al. 2022), No association (Hansford et al. 2022) |
|  |  |  |  | *Amblyomma helvolum* | Harmful (Kwak et al. 2021) |
|  |  |  |  | *Dermacentor auratus* | Harmful (Kwak et al. 2021) |
|  |  |  |  | *Haemaphysalis semermis* | Harmful (Kwak et al. 2021 |
|  |  |  |  | *Ixodes granulatus* | Harmful (Kwak et al. 2021) |
|  |  |  | Woodland edge | *Ixodes ricinus* | Harmful (Hansford et al. 2017, Hansford et al. 2022) , No association (Hansford et al. 2022) |
|  |  |  | Deciduous forest | *Ixodes ricinus* | Harmful (Michalik et al. 2003, Di Luca et al. 2013), No association (Hauck et al. 2020) |
|  |  |  |  | *Ixodes inopinatus* | No association (Hauck et al. 2020) |
|  |  |  | Coniferous forest | *Ixodes ricinus* | Beneficial (Michalik et al. 2003) |
|  |  |  | Mixed forest | *Ixodes ricinus* | Harmful (Michalik et al. 2003, Hauck et al. 2020), No association (Hauck et al. 2020) |
|  |  |  |  | *Ixodes inopinatus* | Harmful (Hauck et al. 2020), No association (Hauck et al. 2020) |
|  |  | Urban grassland | | *Ixodes ricinus* | Harmful (Hansford et al. 2021, Hansford et al. 2022) , No association (Hansford et al. 2022) |
|  |  | Scrubland | | *Amblyomma helvolum* | No association (Kwak et al. 2021) |
|  |  |  |  | *Dermacentor auratus* | No association (Kwak et al. 2021) |
|  |  |  |  | *Haemaphysalis semermis* | No association (Kwak et al. 2021) |
|  |  |  |  | *Ixodes granulatus* | No association (Kwak et al. 2021) |
|  |  | Presence of log and brush piles | | *Amblyomma americanum* | Harmful (Gregory et al. 2022) |
|  |  |  |  | *Ixodes scapularis* | Harmful (Gregory et al. 2022) |
|  |  |  |  | *Haemaphysalis longicornis* | Harmful (Gregory et al. 2022) |
|  |  | Vegetable of flower garden | | *Amblyomma americanum* | No association (Gregory et al. 2022) |
|  |  |  |  | *Haemaphysalis longicornis* | No association (Gregory et al. 2022) |
|  | Vegetation index | Urban canopy cover | | *Ixodes ricinus* | Harmful (Hansford et al. 2021) |
|  |  |  |  | *Ixodes scapularis* | Harmful (VanAcker et al. 2019, Gregory et al. 2022) |
