## Additional 8 Table S5 for "Green cities and the risk for vector-borne disease transmission for humans and animals: a scoping review"

| **TYPE OF DESCRIPTION** | **GROUP OF DESCRIPTION** | **CATEGORY** | **SUBCATEGORY** | **VECTOR BORNE DISEASE (Pathogen)** | **VECTOR SPECIES** | **EFFECT OF URBAN VEGETATION (and REFERENCES)** |
| --- | --- | --- | --- | --- | --- | --- |
| ***Urban green space typology*** | Connectivity | | | Lyme (*Borrelia*) | *Ixodes ricinus, Ixodes scapularis* | Harmful (VanAcker et al. 2019), No association (Heylen et al. 2019) |
|  |  |  |  | Anaplasmosis (*Anaplasma phagocytophilum*) | *Ixodes ricinus* | No association (Heylen et al. 2019) |
|  |  |  |  | Piroplasmosis (*Babesia*) | *Ixodes ricinus* | No association (Heylen et al. 2019) |
|  | Type of urban green space | Urban park | | Lyme (*Borrelia*) | *Ixodes ricinus, Ixodes hexagonus, Dermacentor reticulatus* | No association (Kybicova et al. 2017, Hornok et al. 2014, Levytska et al. 2021) |
|  |  |  |  | Anaplasmosis (*Anaplasma phagocytophilum*) | *Ixodes ricinus, Ixodes hexagonus, Dermacentor reticulatus* | Harmful (Kybicova et al. 2017, Hornok et al. 2014), Variable (Shorn et al. 2011), No association (Hornok et al. 2014, Levytska et al. 2021) |
|  |  |  |  | Rickettsiosis (*Rickettsia*) | *Ixodes ricinus, Ixodes hexagonus, Dermacentor reticulatus* | Variable (Schorn et al. 2011), No association (Levytska et al. 2021) |
|  |  |  |  | Babesiosis (*Babesia*) | *Ixodes ricinus, Ixodes hexagonus, Dermacentor reticulatus* | No association (Schorn et al. 2011, Levytska et al. 2021) |
|  |  |  |  | Bartonellosis (*Bartonella*) | *Ixodes ricinus, Ixodes hexagonus, Dermacentor reticulatus* | No association (Levytska et al. 2021) |
|  |  | Urban forest | | Anaplasmosis (*Anaplasma phagocytophilum*) | *Ixodes ricinus* | No association (Hornok et al. 2014, Welc-Falęciak et al. 2014) |
|  |  |  |  | Rickettsiosis (*Rickettsia)* | *Ixodes ricinus* | Variable (Welc-Falęciak et al. 2014) |
|  |  |  |  | Lyme (*Borrelia*) | *Ixodes ricinus* | Harmful (Hornok et al. 2014) |
|  |  | Residential areas | | Rickettsiosis (*Rickettsia*) | *Amblyomma americanum* | No association (Small et al. 2021) |
|  |  |  |  | Ehrlichiosis (*Ehrlichia*) | *Amblyomma americanum* | No association (Small et al. 2021) |
|  |  | Generic urban green spaces | | Anaplasmosis (*Anaplasma phagocytophilum*) | *Ixodes ricinus* | Variable (May et al. 2014) |
|  |  |  |  | Lyme (*Borrelia*) | *Ixodes ricinus* | No association (Kubiak et al. 2019), Beneficial (Gryczynska et al. 2021), Variable (Juntila et al. 1999) |
|  |  |  |  | Rickettsiosis (*Rickettsia*) | *Ixodes ricinus* | Variable (May et al. 2014, Overzier et al. 2013) |
|  | Surrounding land cover | % developed lands | | Ehrlichiosis (*Ehrlichia*) | *Amblyomma americanum* | Beneficial (Noden et al. 2022), No association (Noden et al. 2022) |
|  |  |  |  | Babesiosis (*Theileria cervi, Babesia*) | *Amblyomma americanum* | No association (Noden et al. 2022), Beneficial (Noden et al. 2022) |
|  |  |  |  | Lyme (*Borrelia*) | *Amblyomma americanum* | No association (Noden et al. 2022) |
|  |  |  |  | Rickettsiosis (*SPG-Rickettsia*) | *Amblyomma americanum* | No association (Noden et al. 2022) |
| ***Description of urban vegetation*** | Presence of vegetation | | | Lyme (*Borrelia*) | *Ixodes ricinus* | No association (Hansford et al. 2017) |
|  | Type of vegetation | Urban forest | Woodland | Lyme (*Borrelia*) | *Ixodes ricinus* | Harmful (Hansford et al. 2021), No association (Hansford et al. 2022) |
|  |  |  | Woodland edge | Lyme (*Borrelia*) | *Ixodes ricinus* | Harmful (Hansford et al. 2022), No association (Hansford et al. 2022) |
|  |  |  | Deciduous forest | Lyme (*Borrelia*) | *Ixodes ricinus* | No association (Michalik et al. 2003) |
|  |  |  | Mixed forest | Lyme (*Borrelia*) | *Ixodes ricinus* | No association (Michalik et al. 2003) |
|  |  | Urban grassland | | Lyme (*Borrelia*) | *Ixodes ricinus* | Harmful (Hansford et al. 2021) |
|  | Vegetation index | Canopy cover | | Lyme (*Borrelia*) | *Ixodes ricinus* | Harmful (Hansford et al. 2021, Adalsteinsson et al. 2012) |
|  | Litter composition | Woody debris | | Lyme (*Borrelia*) | *Ixodes dentatus, Ixodes scapularis, Haemaphysalis leporispalustris* | Harmful (Adalsteinsson et al. 2012) |
|  |  | Leaf litter | | Lyme (*Borrelia*) | *Ixodes dentatus, Ixodes scapularis, Haemaphysalis leporispalustris* | Beneficial (Adalsteinsson et al. 2012) |
|  | Plant species | *Rosa multiflora* | | Lyme (*Borrelia*) | *Ixodes dentatus, Ixodes scapularis, Haemaphysalis leporispalustris* | Harmful (Adalsteinsson et al. 2012) |
